# Causal feature selection using a knowledge graph combining structured knowledge from the biomedical literature and ontologies: a use case studying depression as a risk factor for Alzheimer’s disease

**DOI:** 10.1101/2022.07.18.500549

**Authors:** Scott A. Malec, Sanya B. Taneja, Steven M. Albert, C. Elizabeth Shaaban, Helmet T. Karim, Arthur S. Levine, Paul Munro, Tiffany J. Callahan, Richard D. Boyce

**Affiliations:** Department of Biomedical Informatics, School of Medicine, University of Pittsburgh, Pittsburgh, PA USA; Intelligent Systems Program, University of Pittsburgh, Pittsburgh, PA USA; Department of Behavioral and Community Health Sciences, School of Public Health, University of Pittsburgh, Pittsburgh, PA USA; Department of Epidemiology, School of Public Health, University of Pittsburgh, Pittsburgh, PA USA; Department of Psychiatry, University of Pittsburgh, Pittsburgh, PA USA; Department of Bioengineering, University of Pittsburgh, Pittsburgh, PA USA; Department of Neurobiology, School of Medicine, University of Pittsburgh, Pittsburgh, PA USA; The Brain Institute, School of Medicine, University of Pittsburgh, Pittsburgh, PA USA; School of Computing and Information, University of Pittsburgh, Pittsburgh, PA USA; Department of Biomedical informatics, Columbia University, New York, NY USA

**Keywords:** knowledge representation, management, or engineering, knowledge graphs, causal modeling, feature selection, Alzheimer’s disease, depression

## Abstract

**Background:** Causal feature selection is essential for estimating effects from observational data. Identifying confounders is a crucial step in this process. Traditionally, researchers employ content-matter expertise and literature review to identify confounders. Uncontrolled confounding from unidentified confounders threatens validity, conditioning on intermediate variables (mediators) weakens estimates, and conditioning on common effects (colliders) induces bias. Additionally, without special treatment, erroneous conditioning on variables combining roles introduces bias. However, the vast literature is growing exponentially, making it infeasible to assimilate this knowledge. To address these challenges, we introduce a novel knowledge graph (KG) application enabling causal feature selection by combining computable literature-derived knowledge with biomedical ontologies. We present a use case of our approach specifying a causal model for estimating the total causal effect of depression on the risk of developing Alzheimer’s disease (AD) from observational data.

**Methods:** We extracted computable knowledge from a literature corpus using three machine reading systems and inferred missing knowledge using logical closure operations. Using a KG framework, we mapped the output to target terminologies and combined it with ontology-grounded resources. We translated epidemiological definitions of confounder, collider, and mediator into queries for searching the KG and summarized the roles played by the identified variables. We compared the results with output from a complementary method and published observational studies and examined a selection of confounding and combined role variables in-depth.

**Results:** Our search identified 128 confounders, including 58 phenotypes, 47 drugs, 35 genes, 23 collider, and 16 mediator phenotypes. However, only 31 of the 58 confounder phenotypes were found to behave exclusively as confounders, while the remaining 27 phenotypes played other roles. Obstructive sleep apnea emerged as a potential novel confounder for depression and AD. Anemia exemplified a variable playing combined roles.

**Conclusion:** Our findings suggest combining machine reading and KG could augment human expertise for causal feature selection. However, the complexity of causal feature selection for depression with AD highlights the need for standardized field-specific databases of causal variables. Further work is needed to optimize KG search and transform the output for human consumption.

**Highlights:**

- Knowledge of causal variables and their roles is essential for causal inference.
- We show how to search a knowledge graph (KG) for causal variables and their roles.
- The KG combines literature-derived knowledge with ontology-grounded knowledge.
- We design queries to search the KG for confounder, collider, and mediator roles.
- KG search reveals variables in these roles for depression and Alzheimer’s disease.

**Graphical abstract:** 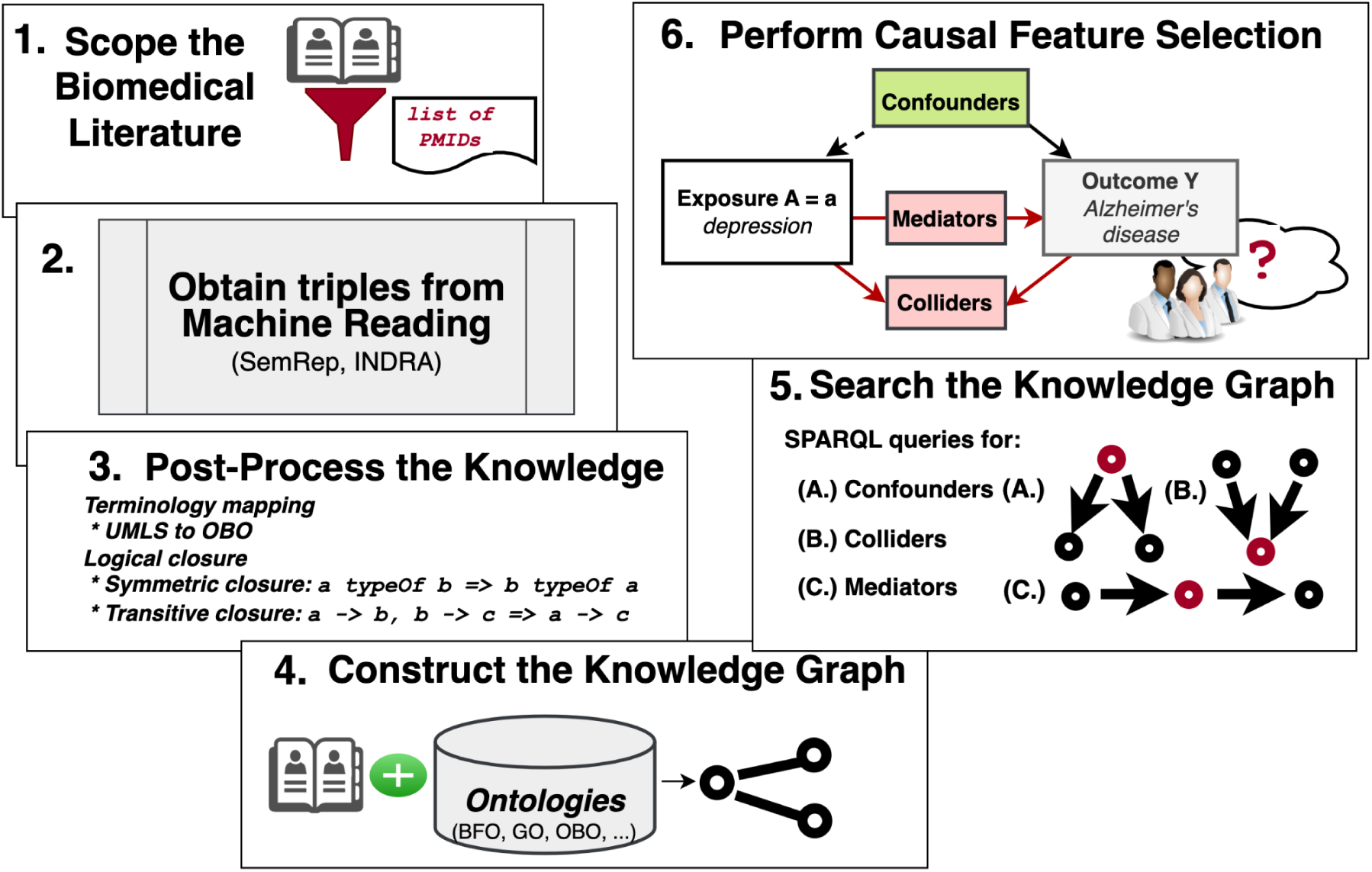

**Statement of Significance:** *Problem:* Extensive knowledge is required to identify confounders for estimating total effects in observational settings. The literature is too vast for humans to process. Bias may remain from not adjusting on unknown confounders or erroneously adjusting on a collider or mediator.

*What is already known:* Structured literature-derived knowledge is often useful but is noisy.

*What this papers adds:* We present a biomedical knowledge graph linking literature-derived and ontology-ground knowledge of biochemical processes with definitions of clinical disease. We search the KG to distill a model for estimating the (total) effect of depression on Alzheimer’s disease from observational data.

## 1. Introduction

Randomized Controlled Trials (RCTs) are widely considered to be the paramount method for determining the causal relationship between exposure and outcome in the medical field^1^. The supremacy of RCTs is due to the efficacy of randomization in eliminating bias by eliminating the relationship between exposure assignment and the health characteristics of study participants that may impact the outcome. However, there are instances where RCTs are not feasible due to ethical or pragmatic reasons. For instance, studying the risk factors for chronic diseases among older adults through RCTs can be impractical, as RCTs typically require short durations and healthy participants. Additionally, the external validity of RCTs can be problematic, as it may not be possible to generalize conclusions from the study to the target population^1^. Therefore, study designs have been developed to address these limitations to estimate the effects of non-randomized, observational data.

However, confounding bias is common when estimating causal effects in observational studies. Confounding bias occurs when a third unrelated factor is linked to both the exposure and the outcome of interest, distorting the true causal effect of the exposure. Researchers can use methods like matching, stratification, restriction, or statistical adjustment to reduce or eliminate this bias. Thus, identifying variables sufficient to control for confounding bias is essential to estimate the effects from observational data reliably^2^. This variable identification task, causal feature selection, is the present study’s focus.

Fortunately, the causal inference literature has produced exacting causal criteria to guide causal feature selection, i.e., to populate a causal model (often in a directed acyclic graph or DAG) ^2–4^. These causal criteria recommend which variables to include in an adjustment set. First and foremost of these criteria is the Principle of the Common Cause or PCC, first expressed by philosopher of science Hans Reichenbach^4^. Essentially, the PCC requires the inclusion of all common causes (confounders) of the exposure and the outcome. More recently, more advanced alternative criteria have been proposed for mitigating scenarios with unmeasured confounders^5^. Nevertheless, applying these causal feature selection criteria to real-world problems depends on accurate, granular knowledge about etiological relationships relating an exposure to an outcome.

Traditionally, the task of causal feature selection depends on prior knowledge of the subject matter through literature review, or expert consultation has fallen to human content-matter experts to identify and carefully select variables for which to control^2, 6, 7^ using a judicious mix of content-matter expertise, literature review, and guesswork^8–11^. However, the literature is vast (PubMed indexes over 1 million articles per year^12^), making this a difficult task for humans to carry out. Knowledge about the underlying causal structure is fragmentary and in constant revision.

The roles (confounder, collider, and mediator) played by variables etiologically related to an exposure and an outcome are depicted in *Figure 1*.

**Figure 1.**
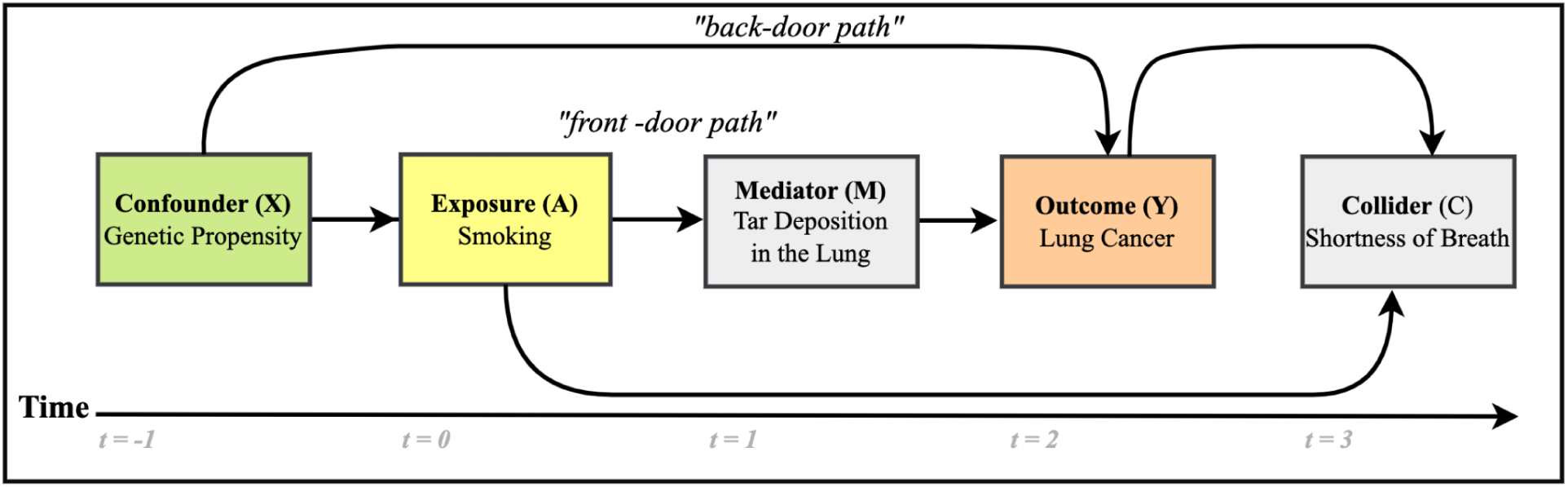
This causal diagram depicts exposure (A) and outcome (Y) in yellow and orange, respectively, along with a confounder denoted “X” with a green background, a mediator variable denoted “M” with a gray background, and a collider “C” also with a gray background. The green background for the confounder means that it is desirable to condition on such variables. Gray indicates that conditioning on that type of variable induces bias.

Causal variables do not always behave solely as confounders. For example, conventional regression-based estimates fail under *treatment confounder feedback*^13^, in which confounders act simultaneously as intermediate variables called mediators^14–16^. Conditioning on mediators is not recommended for estimating the total causal effect as mediators are on the causal path and tend to bias effect measures. Variables that act simultaneously as confounders and mediators are also called **c**onfounders **a**ffected by **p**rior **t**reatment, or CAPTs^17^. CAPTs should not be adjusted for unless estimation methods designed for this purpose are used.

Additionally, variables called colliders, the common effects of the exposure and outcome, should not be used for conditioning because they induce collider bias that distorts measures of effect. By introducing non-causal dependency, collider bias can inflate a null effect into a positive or negative effect or reverse the direction of the effect^18^. In a well-known example of collider bias, a strong association was detected between locomotor disease (exposure) and respiratory disease (outcome): odds ratio (OR) 4.09. In this case, collider-stratification bias resulted from looking only at hospitalized patients. When the data included a mix of hospitalized patients and the general population, this association was absent (OR 1.06)^19^. Thus, because variables may not act in well-defined categories, it is important to be aware of potentially ill-behaved covariates when selecting features so that variables with complex behavior can be dealt with optimally. In our Discussion, we address these and related issues in more depth.

The inability to ensure comprehensive knowledge can mean that important confounders are not controlled for^20, 21^ while inappropriate variables are^19, 22–24^. Fortunately, it may be possible to reduce confounding bias better and prevent inducing bias from the inappropriate selection of variables by considering a more comprehensive set of potential confounders using computational tools.

Harnessing computable causal knowledge made accessible by machine reading could improve the comprehensiveness of causal feature selection. Machine reading works by capturing concepts^25^, or units of meaning representing specific biomedical entities or ideas in a text, e.g., diseases, symptoms, treatments, but also how different concepts are related by automatically extracting knowledge from the biomedical literature in subject-predicate-object triples, also called *semantic predications*, e.g., “ibuprofen **TREATS** migraine.” These concepts are then represented in standardized terminologies and ontologies that describe and communicate healthcare-related information. There is substantial literature on workflows deploying triples from machine reading systems in a broad range of biomedical applications from pharmacovigilance^26–29^, cancer research^30, 31^, and drug repositioning^32, 33^. However, machine reading output has potential weaknesses. For instance, the recall of machine reading is low relative to other natural language processing tasks since machine reading systems often rely on overly precise manually-crafted patterns^34^. Furthermore, machine reading output may be incomplete. For instance, while a machine reading system may accurately detect the above predication, critical implied information may be missed.

For example, as human semantic reasoners proficient in English, it may be possible to infer that therapeutic potential extends to all NSAIDs, the broader category of drugs to which ibuprofen belongs, or that ibuprofen and the other NSAIDs could also treat migraine. Other weaknesses include that machine reading systems can produce erroneous output that is unsuitable *as is* for downstream applications, e.g., causal feature selection^35^. We address these challenges further in the Discussion.

Fortunately, biomedical ontologies contain authoritative, expert-curated representations of entities and relationships between those entities that are exploitable by computer programs. Causal knowledge in these ontologies has been used to inform causal models in AD research^36^ and elsewhere^37–41^. This knowledge could be used to filter out erroneous assertions from machine reading by distilling those assertions through the ontologies created by domain experts. However, like the literature, biomedical ontologies are incomplete due to the extensive manual effort to build and maintain them^42^.

The computable literature-derived knowledge from the literature and ontology-grounded resources are often consolidated into a knowledge graph (KG) representation. A KG is defined as a “graph-theoretic representation of human knowledge such that it can be ingested with semantics by a machine^43^.”In biomedical KGs, entities such as disease phenotypes, genes, enzymes, ligands, and diseases are represented as nodes, while edges (or arrows) denote relationships (predicates) between the nodes. Certain KG implementations can employ ontology-grounded information to prune erroneous inputs and thus may partially overcome the limitations of relying exclusively on structured information extracted from the literature.

This paper introduces a novel causal feature selection method using a KG application combining literature-derived structured knowledge with the precise semantics of ontology-grounded resources. ***Our approach entails constructing and searching a large biomedical KG and provides a proof of concept in a use case identifying and refining a set of confounders for estimating the total causal effects of depression on Alzheimer’s disease (AD)*.**

AD is the fourth most common cause of death for those 75 years of age or older^44^ and is a progressive, multifactorial neurodegenerative disease and the most common cause of dementia. Although most commonly associated with progressive memory loss, AD impacts cognitive ability across multiple domains^45^, including executive and language function, sleep, mood, and personality. Over time, it reduces a person’s autonomy and leads to death. Despite AD’s significant and growing burden, major knowledge gaps remain in AD diagnosis, treatment, and prevention, and only two FDA-approved disease-modifying therapies exist (Iacanumab and Aducanumab). Depression is a common mental illness where the person focuses on negative emotions with symptoms of restlessness, inability to focus, irritability, fatigue, and suicidal thoughts, and may be accompanied by coping behavior, e.g., overeating^46, 47^.

We chose to study the relationship between depression and AD as a use case for the following reasons. First, the two conditions are multifactorial and heterogeneous in nature and have overlapping clinical definitions, common risk factors, and etiological mechanisms^48–52^. Hence, we expect the structure of the causal neighborhood of variables related to both depression and AD to be inherently challenging and complex. We would expect to find variables that play multiple roles. Second, recent insights into the modifiable risk factors present unprecedented opportunities for preventive interventions to reduce AD’s disease burden dramatically. The Lancet Commission on dementia identified depression among eleven other modifiable risk factors, reporting that 40% of AD diagnoses are attributable to these modifiable risk factors^53^. Thus, if depression were truly an independent risk factor, reducing the prevalence of depression could also reduce the population-level disease burden of AD.

However, controversy surrounds depression as either an independent and modifiable risk factor or a prodrome of AD in late life, making depression an interesting condition to study. Lastly, depression is a highly prevalent disorder. In 2018, it was the third most common mental illness; by 2030, it is expected to become the number one most prevalent mental illness^47^. The prevalence of depression has also been exacerbated by current events, including the COVID-19 pandemic^54^. Thus, if the relationship between depression and AD is genuinely causal, intervening in depression could significantly impact public health. Taken together, the depression and AD relationship is a good fit as a use case since the objective of our causal feature selection method could address a critical need to clarify the etiological status of depression for its effect on AD risk in future studies by furnishing a more comprehensive set of confounders.

This work aims to advance the state-of-the-art causal feature selection method that combines literature-derived structured knowledge with the precise semantics of ontology-grounded resources. The essential idea of this study is to use information extracted from the literature to define the domain of discourse (here, the set of variables potentially related to depression and AD) and then filter out the variables based on biological plausibility using an ontology-grounded KG. We hypothesize that:

- It is possible to identify plausible confounders, colliders, and mediators from structured knowledge distilled from the literature through an ontology-grounded KG (**H1**). *Some of the variables identified by the KG search would also appear in the results of complementary methods and be reported in published observational studies*.
- There is a high degree of complexity regarding the variables linking depression and AD because of the bi-directional nature of the relationship between depression and AD (**H2**). Hence, *we expect many variables, including confounders, to play multiple causal roles*.
- The reasoning paths traversed searching the KG can produce reasonable hypotheses for the mechanisms underlying the relationships between the variables and clarify their meaning (**H3**).

In the next section, we describe the workflow for extracting and post-processing the structured knowledge from the literature, constructing the KG, and reporting and evaluating the KG search results. Then, we report the KG search results and evaluation. Finally, we interpret these results in the context of our use case and clarify the strengths and weaknesses of the approach to inform future work.

## 2. Methods and Materials

*Figure 2* shows the workflow for this study. We first delineated the scope of the corpus of articles in PubMed by their PubMed identifiers. Next, we extracted triples from the titles and abstracts from the articles in the corpus using machine reading systems (SemRep^55–57^ along with EIDOS and REACH in the INDRA^58^ ecosystem) and combined and harmonized the outputs. We refer to the graph containing the outputs, i.e., the structured knowledge derived from machine reading of the literature (and in the post-processing stages), as the *extraction graph*. Next, since there may be information logically entailed by the extracted information, we applied forward-chaining over the extracted triples to infer implied triples based on logical properties, e.g., transitivity, symmetry, asymmetry, and reflexivity, of predicates extracted by machine reading. The terminology-mapped and logically closed extraction graph was combined with an ontology-grounded KG constructed with the PheKnowLator workflow developed by Callahan et al.^59–61^. We then translated standard epidemiological criteria for confounders, colliders, and mediators into discovery patterns, which are semantic constraints over the relations between concepts^62^. We implemented those discovery patterns as SPARQL queries for searching the KG based on our previous research^63^. We analyzed the KG search results for evaluation by compiling the variables by their simple and combined roles and comparing these results with reported confounders from published observational studies that we manually collected from a pilot meta-review and the output of a complementary method (searching the Semantic MEDLINE database, or SemMedDB^64^). Finally, we manually inspected a selection of variables by inspecting the source sentences, interpreting explanatory reasoning paths produced by the search, and searching PubMed.

**Figure 2.**
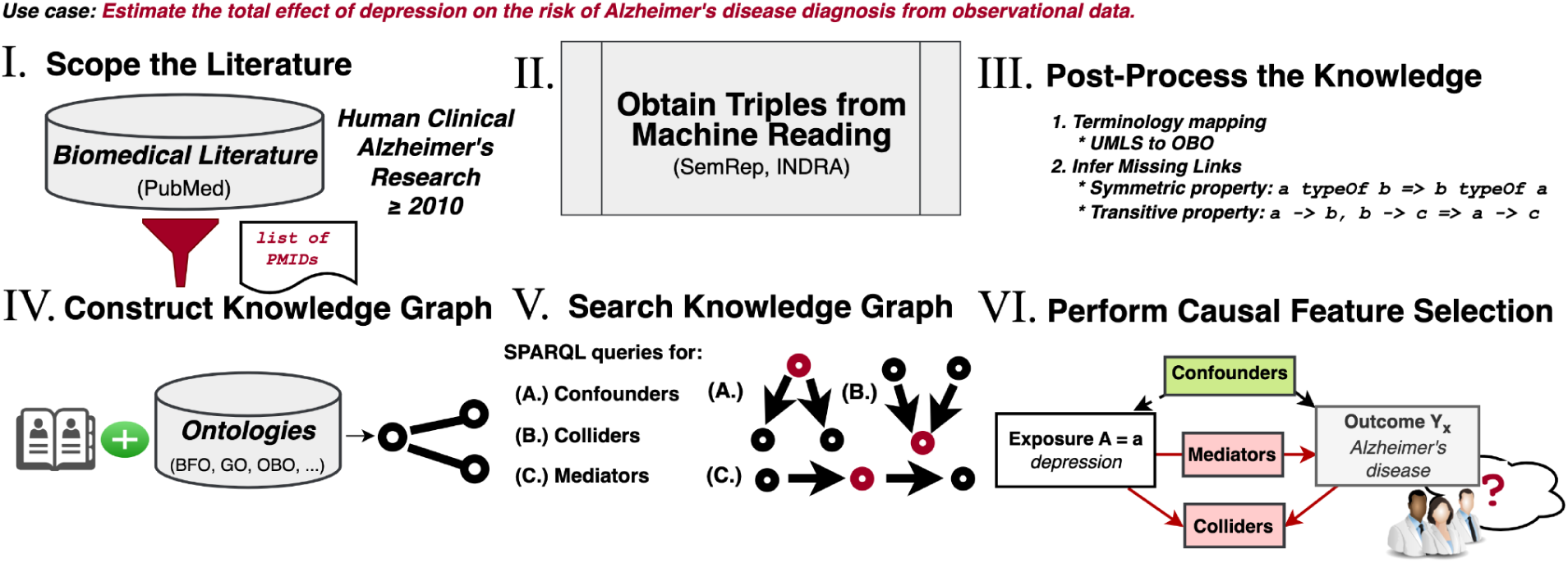
The principal stages of the workflow to construct the causal feature selection system. **I.** Scope the literature to clinical studies investigating AD, published in 2010 or after. **II.** Obtain triples from the machine reading systems. **III.** Post-process the knowledge by performing logical closure operations with the CLIPS production rule system^65, 66^ to infer missing edges and terminology mapping. **IV.** Construct the KG using the PheKnowLator platform, merging the output of the machine reading systems with ontology-grounded information. **V.** Search the KG to identify relevant variables. **VI.** Compile the search results into a causal model analyzing KG search result output and comparing that output with the structured knowledge in SemMedDB, the results of a pilot meta-review of reported confounders collected from observational studies, and PubMed.

We provide details for each component in the paragraphs below and document constructing the extraction graph and KG search. The code we produced for this work is available on the Zenodo project repository, https://doi.org/10.5281/zenodo.6785307. Documentation for how to use this code archive is available in Appendix C in the Supplementary Materials.

### 2.1 Scoping the literature

We consulted with a health sciences librarian to design a PubMed query (see **File II. (*Appendix_C.docx***), **Listing S1**) to limit the indexed literature’s scope to human research related to AD-related associations and risk factors. The idea is to ingest a significant fraction of the concepts in the AD domain while having a tractable data set. We limited the search to 10 years before the search date (July 11, 2020) to a subset of the literature related solely to AD. This is because different types of dementia may have different underlying etiological mechanisms and risk factors.

### 2.3 Obtaining triples from machine reading

#### SemRep

SemRep is a natural language processing tool developed by the National Library of Medicine (NLM) for extracting semantic triples from biomedical literature^55^. SemRep uses a named entity recognition system called MetaMap^67–69^. MetaMap transforms mentions of particular biomedical entities identified as components of semantic triples to “concepts” in the Unified Medical Language System (UMLS) metathesaurus^55^. Each concept in the UMLS metathesaurus is assigned a unique handle called a concept unique identifier or CUI. SemRep infers relations between entities encountered in the biomedical text using a part-of-speech tagger and a semantic network informed by semantic constraints and outputs semantic triples. SemRep has been evaluated extensively over its existence (now almost two decades). Evaluations focusing on SemRep’s intrinsic ability to extract semantic triples that accord with human judgment have yielded 0.55-0.9 precision, 0.24-0.42 recall, and 0.38-0.5 F_1_. For more extensive information on SemRep, see an overview of Semrep by Kilicoglu^55^.

We used the triples present in the Semantic MEDLINE database, SemMedDB. We downloaded SemMedDB version 40 (on June 1st, 2020). We searched SemMedDB for the triples associated with the PubMed identifiers resulting from the literature scoping query.

#### INDRA

The Integrated Network and Dynamical Reasoning Assembler, or INDRA, is an automated machine reading and model assembly software. We use two machine reading systems developed within the ecosystem, EIDOS and REACH. EIDOS is a rule-based open-domain machine reader that extracts causal and correlational relations between events mentioned in free-text^70, 71^. EIDOS extracts three predicate types - **`event`**, **`influence`**, and **`association`**. **`event`** signifies a change or modification of a concept identified by the machine reader. **`influence`** is a causal relationship between two events, while **`association`** represents a set of events without implying causality. ***<u>RE</u>****ading and **<u>A</u>**ssembling **<u>C</u>**ontextual and **<u>H</u>**olistic mechanisms from text*, or ***<u>R EACH</u>***, detects mentions of protein-modifying biochemical processes called post-translational modifications in text. Mass fluorescence spectroscopy has allowed for the precise characterization of different post-translational modifications and opened the possibility of discovering novel biomarkers, therapeutic strategies, or drug targets involving protein interactions. In addition, a burgeoning literature in AD research^79–81^ has revealed that many processes involving proteins related to AD’s pathogenesis are related to perturbations to processes involving post-translational modifications. REACH provides a window into these processes. *Table 1* lists the predicates detected by the machine reading systems we analyze in the paper.

**Table 1.** This table displays the subset of predicates we analyze in this paper, organized by machine reading system and semantic domain. These machine reading system predicates were chosen because they describe information essential for causal feature selection and modeling.

| **READER** | **Semantic Domains** | **Predicates** |
| --- | --- | --- |
| **SemRep** | Clinical Medicine | **TREATS, PREVENTS** |
|  | Molecular Interactions | **INTERACTS_WITH, INHIBITS, STIMULATES** |
|  | Disease Etiology | **ASSOCIATED_WITH, CAUSES, PRECEDES, PREDISPOSES** |
|  | Pharmacogenomics | **AFFECTS, AUGMENTS, COMPLICATES, DISRUPTS, INHIBITS, STIMULATES** |
|  | Static Relations | **PART_OF, COEXISTS_WITH** |
| **INDRA/EIDOS** | General Domain | **ASSOCIATION, INFLUENCE** |
| **INDRA/REACH** | Biochemical Reactions (involving covalent modification of a protein by attaching a functional group) | **PHOSPHORYLATION, UBIQUITINATION, HYDROXYLATION, SUMOYLATION, GLYCOSYLATION, ACETYLATION, FARNESYLATION, RIBOSYLATION, METHYLATION** |
|  | Nested Events / “Regulations” | **REGULATES, PROMOTES, INHIBITS, ACTIVATES** |

### 2.3 Knowledge post-processing

#### 2.3.1 Terminology mapping

The purpose of knowledge post-processing is to improve the accuracy and completeness of the literature-derived knowledge. In this step, we use knowledge hygiene via terminology mapping and automated reasoning to transform the output from machine reading into an extraction graph with inferred information from applying logic and with concepts mappable to an ontology-grounded KG. Knowledge hygiene is required to make the machine reading output more amenable to inference. Knowledge hygiene addresses issues with the completeness and validity of the extracted data, including redundancy.

We performed the following steps on the triples extracted by the machine reading systems. First, triples were removed with phrases in the subject or object position with more than three space-delimited tokens, e.g., “alpha-blockers aggravation cognitive dementia patients,” from the corpus since these phrases could not be condensed to a single concept. Second, we filtered out irrelevant predicates, e.g., **`location_of`**, and retained only the predicates listed in *Table 1*. We also removed duplicate triples. Third, we assigned a default probabilistic belief score of 0.8 to all SemRep triples. The belief score is a value that represents the level of confidence we place in the accuracy of the triple and ranges on a scale from 0 (no confidence) to 1.0 (certainty). The value of 0.8 was chosen because it is the average published performance characteristic across SemRep’s various predicates^64, 75^.

Fourth, we mapped the REACH predicates to **`ro:0002436 molecularly_interacts_with`**, except for phosphorylation, which was mapped to **`ro:0002447 phosphorylates`** in the Relation Ontology (RO)^76^. We used the built-in INDRA preassembly module to remove duplicate triples and assign belief scores for the extracted triples from REACH and EIDOS with the INDRA belief module. From EIDOS, we included all unique **`influence`** and **`association`** triples. We also mapped the biomedical entities appearing in the subject and object positions in the extracted INDRA triples to UMLS concepts using MetaMap^67, 68^ (version 1.8 with UMLS version 2018AA) and obtained the CUI, UMLS preferred name, and semantic type for the top-scoring match from MetaMap for each concept. All triples were removed from the corpus used to create the extraction graph where the subject, predicate, or object did not have a corresponding mapping in UMLS. Lastly, the triples from all three readers were converted to tab-separated files in a combined corpus that included one row for each subject, predicate, object (with UMLS mapping information), belief score, source sentence, and PubMed ID of the original text and combined the machine readers’ output into an extraction graph. The extraction graph contains only the literature-derived triples from each of the readers.

#### 2.3.2 Logical closure over the extraction graph

Machine reading systems typically do not consider logical entailment concerning the logical properties of the predicates they detect. Consequently, the extraction graph may be incomplete regarding knowledge that could be inferred by applying simple logic.

Therefore, it is essential to consider the semantic domains of a dataset and the different categories of meaning present within it. The predicates in *Table 1* can be subgrouped into broad semantic categories of causal, associative, and subsumptive predicates. For example, causal predicates such as **`causes`** describe a cause-and-effect relationship between two entities. In comparison, associative predicates such as **`associates_with`** and **`coexists_with`** describe a relationship in which two entities are frequently found together. On the other hand, subsumptive predicates such as **`part_of`** describe relationships in which one concept is in a hierarchical (“ad **IS_A** dementia”) or a mereological (from the Greek μερος, ‘part’) *part-whole* relationship^77, 78^ with another.

The logical properties of predicates are essential for knowledge representation and inference, especially in machine reading. Predicates belonging to the causal, associative, and subsumptive categories possess logically defined properties such as transitivity, symmetry, asymmetry, and reflexivity that enable the reasoning and inference of new knowledge about relationships between entities. We can derive additional knowledge from the data through inferences by utilizing these logical properties. Detailed operation definitions and examples of these properties may be found in File **I.**, **Background S2.** Logical properties.

To exploit these properties and logically infer entailed but missing triples, we mapped a subset of the machine readers’ predicates to their logical definition in terms of its logical properties using the logical definition from the object properties in the Relation Ontology (RO)^79^. The UMLS to RO mappings are available in file **II.**, **Table S1.** and the analyses of logical properties of the predicates in Table 1 are in file **IV *predicatePropertyAnalysis.xlsx*** for each machine reading system.

We then used the transitive, symmetric, asymmetric, reflexive, and inverse properties present in RO in a procedure to infer new triples entailed by the extraction graph^80, 81^. This procedure entails loading the initial extraction graph into the CLIPS forward-chaining production rule system ^82^ (https://clipsrules.net/) via the CLIPSPy Python library^66^. Simple rules using symmetric and transitive relationships belonging to RO predicates were applied to infer new triples and add them to the extraction graph. The CLIPS inference engine also tracked belief scores to manage uncertainty about the validity of inferred triples. New triples inferred from symmetric inference were assigned the same belief score as the asserted triple. The inference engine multiplied the belief score for each of the two asserted triples in transitive inference and assigned it to the inferred triple. All triples inferred from transitivity with a belief score of ≤ 0.6 were filtered out. The value 0.6 was chosen as a heuristic threshold to mimic a weakened transitivity property^83^, limiting transitive inference beyond one hop. For example, recalling that SemRep triples were assigned a belief score of 0.8, the first inferred triple from a transitive SemRep predicate would be retained because, through the application of the chain rule where each node is dependent on the previous nodes, 0.8 * 0.8 = 0.64. Subsequently, inferred triples would be filtered out since 0.8 * 0.8 * 0.8 < 0.6.

##### Listing 1. Ontologies merged into our implementation of PheKnowLator version 2.0.

Cell Line Ontology (CLO)
Chemical Entities of Biological Interest (ChEBI) Gene Ontology (GO)
Human Disease Ontology (DO)
Human Phenotype (HP)
Pathway (PW)
PRotein Ontology (PRO)
Relation Ontology (RO)
Sequence Ontology (SO)
<u>Uber</u>-anatomy Ontology (UBERON)
Vaccine Ontology (VO)

### 2.4 Constructing the KG

Phenotype Knowledge Translator (PheKnowLator) is an Ecosystem and Python 3 library designed to construct semantically rich, large-scale biomedical KGs from biomedical ontologies and complex heterogeneous data^84^. We cloned the PheKnowLator 2.0 code and ran the ontology hygiene and data preparation documented in the project’s Jupyter workbooks. PheKnowLator merged several ontologies represented using OWL^85^ related to genes, diseases, proteins, chemicals, vaccines, and other ontologies of biological interest. These ontologies are enumerated in *Listing 1*. Complete information about the resources used in PheKnowLator (version 2.0) is available on GitHub^86^. Each class in the ontologies merged by the PheKnowLator process formed nodes in the KG. Thus, the object properties assigned to each class formed an edge in the KG. The edges were extended by integrating data sources that connected classes using object properties provided by the RO^79^. Finally, the resulting KG was saved as an OWL ontology and a Python NetworkX graph object^87^. For more information on the Jupyter notebook, the mapping, and logical closure, see **File VII. *Appendix_C.docx***.

#### 2.4.1 Combining the Extraction Graph with PheKnowLator

Several steps were required to integrate the extraction graph with the PheKnowLator KG so that we could study its application in causal feature selection. The first step was to map the UMLS concept unique identifiers in the SemRep output to the OBO Foundry identifiers used by the PheKnowLator KG. We used existing mappings in the UMLS for the Gene Ontology and Human Phenotype Ontology classes supplemented by a query of the Bioportal Annotator service for other ontologies^81^. The REACH machine reader automatically generated mappings to OBO Foundry identifiers which we then added to a mapping table. We also mapped SemRep predicates to the RO version used by the PheKnowLator workflow as SemRep predicates when those predicates were not present in the RO.

#### 2.4.2 Reweighting the edges to support path search

The PheKnowLator KG had features deriving from the knowledge representation that made querying and path search challenging without further processing. The downstream use of the weighting scheme described in this section was to influence the shortest path and simple path searches so that they would include fewer edges and nodes that were present in the knowledge graph to support OWL semantics and more edges and nodes that were biologically interesting. The ontologies were written in the OWL description logic language, which uses logical statements to define the semantics of ontology entities. Our approach used a reweighting procedure to distill the KG into biologically meaningful entities useful for our study. Therefore, we used the following steps and heuristics to remove OWL semantic statements and reweight the edges of the KG:

1.) **Apply OWL-NETS to the KG to simplify Web Ontology Language (OWL)-encoded biomedical knowledge into an easier-to-use network representation.** OWL-NETS replaced the more complex logical statements with simpler triples without losing biologically relevant information (for examples, see https://github.com/callahantiff/PheKnowLator/wiki/OWL-NETS-2.0)^88^.
2.) **Remove all KG edges that specify a disjoint relationship between a subject and object in a triple.** A disjoint relationship in OWL specifies that the class extension of a class description has no members in common with the class extension of another class description^81^. While disjoint relationships are helpful for deductive closure of the KG using a description logic reasoner, they lead to uninformative results from a path search that seeks a mechanistic explanation.
3.) **Remove all edges in the KG that specify the domain of a relationship from the RO.** Many RO relations specify the ontology classes that comprise the relations’ domain and range. Like the disjoint relationships mentioned earlier, including these edges helps maintain logical consistency during deductive closure but leads to uninformative results from a path search that seeks a mechanistic explanation.
4.) **Reweight each edge in the graph to influence the path search on predicates useful for mechanistic explanations.** The weighting strategy is based on observing the method used in PheKnowLator to represent triples of value for constructing a mechanistic explanation explained below:
  a.) **Assign default edge weights.** Since the PheKnowLator KG is not weighted by default, start by assigning a default weight of 2 to all edges. The value of 2 has no special significance except a value greater than 0.
  b.) **Favor hierarchical relationships.** Hierarchical relationships can be very informative for mechanistic explanations. Most of the classes represented in the ontologies that have been merged are hierarchical and are represented using the `rdfs:subClassOf` predicate. If a predicate relates a subject to an object by `rdfs:subClassOf`: weight = 0. Assignment of subClasOf weights to zero lowers the cost of including a hierarchical edge in a path search through the KG, which increases the prevalence of hierarchical edges in path search results.
  c.) **Favor edges providing a mechanistic triples RO.** We developed a simple heuristic to produce mechanistic triples and avoid triples that provide meta-knowledge about entities in an ontology. If the predicate relates a subject to an object by owl:someValuesFrom: weight = 0. Similarly, if the *object* is a RO entity and the predicate is owl:onProperty: weight = 0.
  d.) **Disfavor subjects and edges that primarily describe metadata**. If the *subject* is a RO entity or the edge is `owl:`equivalentClass: weight = 5.

The procedure above aims to prune the KG components that are not useful for inferring biological mechanisms and to optimize the KG for providing explanations of links between biomedical entities in terms of the underlying molecular mechanisms based on information contained in the biomedical ontology inputs.

Finally, we created two copies of the PheKnowLator KG - one that was loaded into a graph data store (OpenLink Virtuoso version 7.20^89^) so that it could be queried using the SPARQL graph querying language, and the other that was saved as a Python NetworkX MultiDiGraph object for the application of path search algorithms. The extraction graph was loaded into the same graph data store as the PheKnowLator KG for querying.

### 2.5 Searching the KG

A primary objective of this paper is to identify possible confounders, colliders, and mediators from the KG search. Hence, identifying the variables that belong to each causal role constitute the competency questions. To answer these competency questions, we translated the standard epidemiological criteria for these variables into discovery patterns, or semantic constraints we use to identify variables of interest. We used these discovery patterns as requirements to construct queries implemented in the SPARQL graph querying language. The SPARQL queries are described in more detail in the paragraphs below. *Table 2* presents each causal role as a competency question with its associated discovery pattern. Note that each dyad in the discovery patterns also implies chains of causation, i.e.,”**A `interacts_with` B `causes_or_influences_condition` C**.”

**Table 2.** Summary of the competency questions and discovery pattern approach to finding answers via KG search. The left column enumerates the competency questions, and the right column describes the graph search for answering the competency questions identifying confounders, colliders, and mediators.

| Competency questions | Discovery patterns |
| --- | --- |
| 1. What are the potential confounders of depression and AD? | **confounder** CAUSES depression;<br>**confounder** CAUSES AD. |
| 2. What are the potential colliders of depression and AD? | depression CAUSES **collider**;<br>AD CAUSES **collider**. |
| 3. What are the potential mediators between depression and AD? | depression CAUSES **mediator**;<br>**mediator** CAUSES AD. |

KG search was implemented using Dijkstra’s shortest path search over the KG based on queries in the SPARQL graph querying language. Dijkstra’s shortest path algorithm was used to identify the shortest paths between nodes *of interest* in the KG. We chose Dijkstra’s shortest path algorithm as a simple way to test proof-of-concept, though there are many other graph search methods. The Dijkstra algorithm is a variant of the best-first search that prioritizes finding the shortest path between potential nodes. (Note that many paths are of the same length.) For more information on the SPARQL queries and to inspect the raw output from running the queries, see **File VII. *Appendix_C.docx***.

The KG search was comprised of two stages. In the first stage, the KG search was implemented as Dijkstra’s shortest path search over the KG based on a query in the SPARQL over the extraction graph. Recall that the extraction graph was the subgraph of the KG containing the literature-derived triples and edges inferred from the logical closure procedure. The purpose of the first stage is to use the concepts identified by machine reading plus the inferred edges to expand the breadth of variables, i.e., improve the completeness of the machine reading outputs.

In the second stage, the query uses the results from the first stage query are used (represented universal resource identifiers or URIs) as the start and end nodes for the input (as the start and end nodes) of the queries over the ontology-grounded KG. The purpose of this stage was to filter the literature-derived information through the ontology-grounded knowledge with the goal of improving accuracy over the literature-derived knowledge. Our discovery pattern-based KG search implementation is constrained to subsumptive and causal predicates. The causal predicates include **`causes_or_contributes_to_condition`**, **`genetically_interacts_with`**, and **`molecularly_interacts_with`**. These ontology-based predicates were chosen because they describe mechanistic causal links between disease and the genomic and biomolecular entities and processes essential for our use case.

To provide hypothetical explanations of the causal association between identified variables and depression and Alzheimer’s disease (AD), our KG search implementation traced the path of nodes and predicates traversed by Dijkstra’s shortest path search algorithm. These paths, which we refer to as reasoning paths, reveal the causal relationships and ontological links between concepts and allow us to derive an explanation linking biochemical processes with molecular definitions of disease from combined literature- and ontology-derived structured knowledge.

We created visualizations of the reasoning paths to facilitate the analysis and interpretation of the variables. As an illustration, *Figure 3* presents a sample reasoning path in its raw form as output from the KG search and in its transformed format that includes ontological and causal predicates. The transformed format provides a clear and intuitive representation of the relationships and causal links uncovered through the KG search process. These visualizations provide hypothetical mechanistic explanations for the causal role played by each variable in the identified associations according to these reasoning paths. Overall, by tracing the reasoning paths and creating visualizations, we can provide hypothetical explanations of the causal associations between the identified variables and depression and AD.

**Figure 3.**
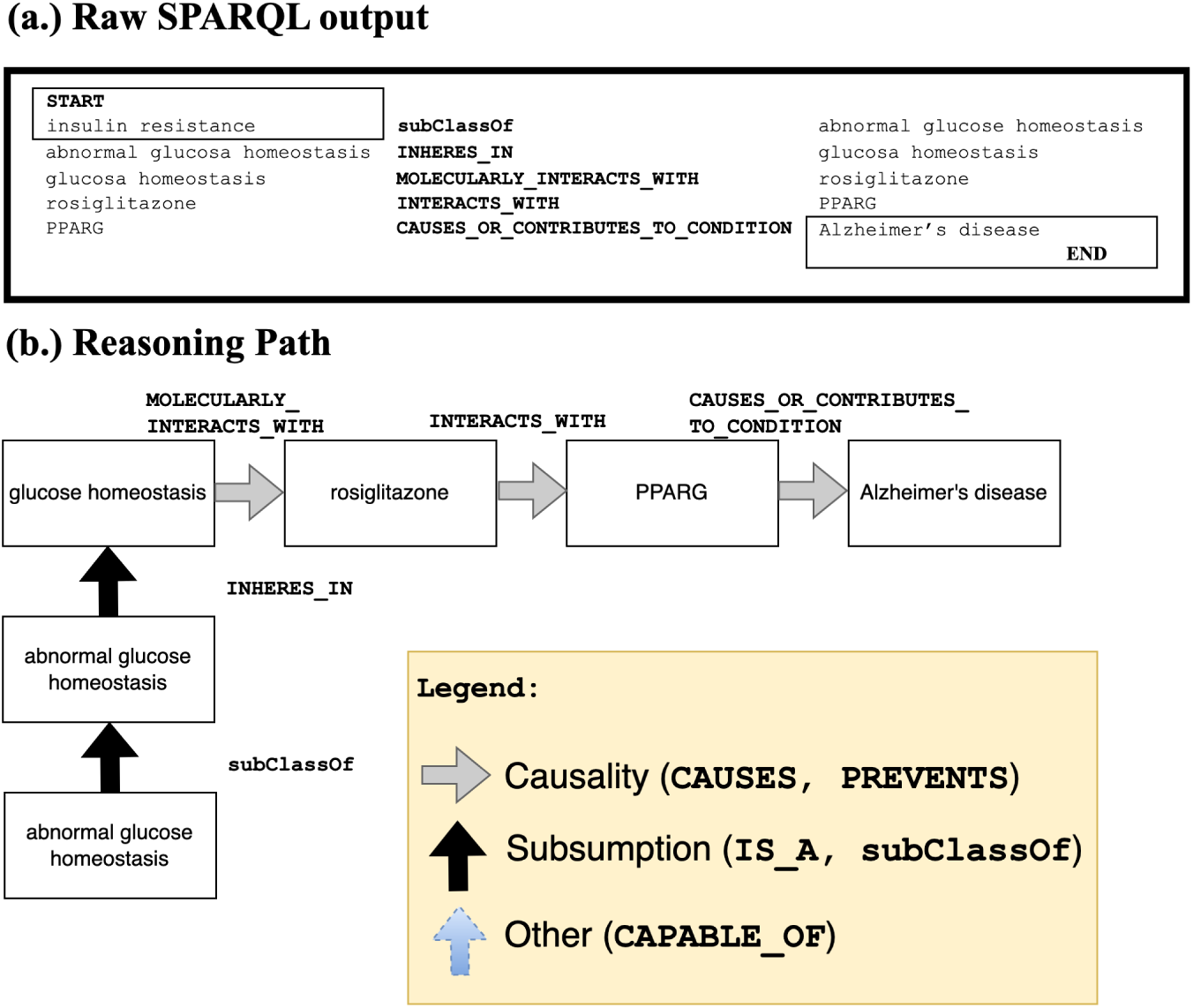
This figure shows two versions of a SPARQL query’s (partial) output. (a.) shows sample raw output from the SPARQL query, and (b.) shows a visualization we manually created from the output. Note that the visualization displays hierarchical (**`is_a`**), mereological (from the Greek μερος, ‘part’), e.g., **`part_of`, `has_part`**, relationships vertically, and causal relationships (**`interacts_with`**, **`causes_or_contributes_to_condition`**) vertically.

Throughout the article, we will present reasoning paths where they serve as useful visual representations of the relationships and causal links discovered in our KG search.

### 2.6 Causal Feature Selection (Evaluation)

Our evaluation procedure is summarized here and described in more detail in the paragraphs below:

1. **Present KG search results for confounder, collider, and mediator variables for depression and AD (***<u>partially answering</u>* **<u>H1</u>).** We conducted a KG search to identify the confounders, colliders, and mediators of depression and AD and reported the KG search results and sample reasoning paths.
2. **Display the KG search results as causal diagrams in two forms: simple roles and combination roles (***<u>partially answering</u>* **<u>H1</u>** *<u>and</u>* **<u>H2</u>).** In addition to the simple roles of confounder, collider, and mediator, our methodology allows for the identification of variables with complex behavior, referred to as combination roles, which are additional role categories for variables with more than one causal role, and include confounder/mediators, confounder/colliders, and confounder/mediator/colliders. These variables with combination roles were identified using set operations, such as set difference and intersection, on the list of variables for each role type.
3. **Compare the KG search results with SemMedDB and reported covariates in published observational studies using Venn diagrams (*<u>partially answering</u>* <u>H1 *and* H2</u>)**. We compared the KG search results with results from searching the SemMedDB_Scoped_ and SemMedDB_Complete_ databases to understand the differences between our novel feature selection and complementary methods for causal feature selection, including the consistency with which concepts are classified by their causal role(s). The search results for simple and combination roles were summarized. Additionally, we compared the KG and SemMedDB_Complete_ with confounders reported in a small database of confounders from a pilot-metareview. For more information on the SQL queries, see File VII. *Appendix_C.docx*.
4. **Prioritize and conduct in-depth analysis on a subset of variables (*<u>partially answering</u>* <u>H1, H*2, and* H3</u>).** A subset of the variables was selected for in-depth analysis to prioritize the results of the KG search. This in-depth analysis aimed to provide further insight into the search results’ veracity and the reasoning paths’ usefulness (**H3**).

As noted earlier, we compiled a database of confounders from a pilot meta-review of confounding control. More information, including details on the pilot meta-review, is provided on each step below.

#### 2.6.1 Confounders, colliders, and mediators KG search

We report the confounders, colliders, and mediators for depression and AD identified by searching the KG. To show the feasibility of our pilot project, KG search should be able to identify causal variables playing each role, and thereby demonstrate competency.

#### 2.6.2 Analysis of search results

In addition to the pre-defined, “simple” roles (confounder, collider, and mediator, mentioned earlier), our compilation procedures allow us to identify variables with complex behavior, i.e., “combination” roles, which we define as additional role categories for variables with more than one causal role. The combination roles included confounder/mediators, confounder/colliders, and confounder/mediator/colliders. Our rationale was to sort out the roles played by the variables and identify variables with complex behavior of which other researchers ought to be aware. The combination roles were identified using set operations, e.g., set difference and intersection, over the list of variables for each role type. For example, to identify “Confounders only” variables, we perform a set difference operation over the set of confounders minus the set of variables appearing in the collider, mediator, or both KG search result sets.

#### 2.6.3 Comparing search results

We sought to understand how our novel feature selection differs from complementary methods for causal feature selection using SQL queries to identify relevant variables from the literature explored in previous research and with confounders reported in published observational studies investigating depression as a risk factor for AD or dementia. To this end, we compared the KG search results with results from searching the original scoped predictions in MachineReadingDB_Scoped_ and SemMedDB_Complete_. MachineReadingDB_Scoped_ contained the original subset of structured knowledge restricted to human clinical studies on AD published in 2010 or after that we used to construct the extraction graph. SemMedDB_Complete_ contained the entire database of SemRep predicates extracted from titles and abstracts of articles published in 2010 or after. We summarized the search results for simple and complex roles as we did in the previous subsubsection. To gain insight into the veracity of our search results, we also conducted a comparison of search results for confounders with confounders reported in the literature. Such a comparison hinges on the existence of a database of reported confounders in the published literature. However, no such standardized and validated causal question-specific confounder database exists. Fortunately, however, we have been developing just such a confounder database in parallel research.

In the pilot meta-review, we manually collected from a small corpus of observational studies investigating depression as the exposure and dementia or AD as the outcome. We based on recommendations from the Cochrane Group^90^. Confounders appearing in the search results that were already recognized in the pilot meta-review of the literature partially validate the soundness of our methods. Information on confounders was manually collected from a small corpus of papers and entered into a spreadsheet. This small corpus included confounders reported in three papers studying depression cited by the Lancet Commission Report (LCR) on late-life modifiable risk factors for dementia^53^. We reason that papers cited in the LCR should be of relatively high quality. Additional reported confounders from four sporadically collected papers over the course of our research. We also collected information on each paper’s exposure and outcome definitions included in the corpus. In this way, we could ensure that only confounders specific to depression and dementia or depression and AD were included in our database. Hence, while the corpus of papers analyzed is small and the scoping strategy was not systematic, reflecting the preliminary stage of this research, which focused on the feasibility of building such a database rather than exhaustiveness, the confounders in the database can be assured to be accurate and reflective of confounders recognized as such by human content-matter experts.

Finally, we compiled the reported confounders into a set containing each confounder to execute the comparison.

Confounders appearing in search results that have already been recognized as such partially validate the soundness of our methods. Search results that do not appear in the literature may indicate that the search results are erroneous or that the search-identified confounder may have been overlooked. Alternatively, since our data collecting strategy was not systematic, our confounder database is not comprehensive. Hence, certain reported confounders may be missed, and potential confounders appearing in the search may not be novel confounders after all.

#### 2.6.4 In-depth analysis

In our study, we conducted several additional *posthoc* analyses on a set of variables obtained from a Knowledge Graph (KG) search. Due to the large number of variables identified through the KG search, we prioritized the subset of variables to be analyzed in-depth based on several criteria, including the potential to address confounding bias, the strength of the relationship as indicated by the strength of the association, scientific interest, and degree of surprise (a concept not present in pilot meta-review but present in the KG and SemMedDB_Complete_, or present but with conflicting assertions causal directionality).

One of the human experts who conducted the verification procedure (CES, with expertise in the population neuroscience of ADRD) was a content expert in Alzheimer’s Disease and Related Dementias (ADRD). The other human expert (HTK) has expertise in bioengineering and focuses on the biology of brain aging.

The human experts examined the source sentences from SemMedDB along with the reasoning paths produced by the KG search for those variables. The task was then to determine if the claims asserted by the KG and SemMedDB_Complete_, i.e., the source sentences from SemMedDB along with the reasoning paths produced by the KG search, were consistent with a combination of the human expert’s background knowledge and information gleaned performing additional searches in PubMed to fill in any gaps in their understanding.

In summary, after prioritizing a subset of variables for further examination, the *post hoc* analysis performed in our study aimed to verify the validity of selected confounders through a human expert-based verification procedure.

## 3. Results

*Figure 4* shows the result of applying post-processing procedures to the extracted triples. 13,365 PubMed-indexed articles were returned from PubMed, resulting in 226,997 triples from machine reading. The number of triples remaining after executing the knowledge hygiene procedure was 94,934, or 41.8% of the original triples. The subject and object entities of the entire set of triples included 10,020 unique UMLS CUIs, of which 2,504 were mapped to ontological identifiers and merged using the PheKnowLator workflow (2,110 from Bioportal, 844 from the Gene Ontology^91, 92^ and Human Phenotype Ontology^93^ mappings present in UMLS, and 394 from REACH). After filtering out the machine reading and inferred triples with a belief score of < 0.8 and mapping predicates to the RO, the total number of machine reading triples was 20,010, or 8.8% of the original total.

**Figure 4.**
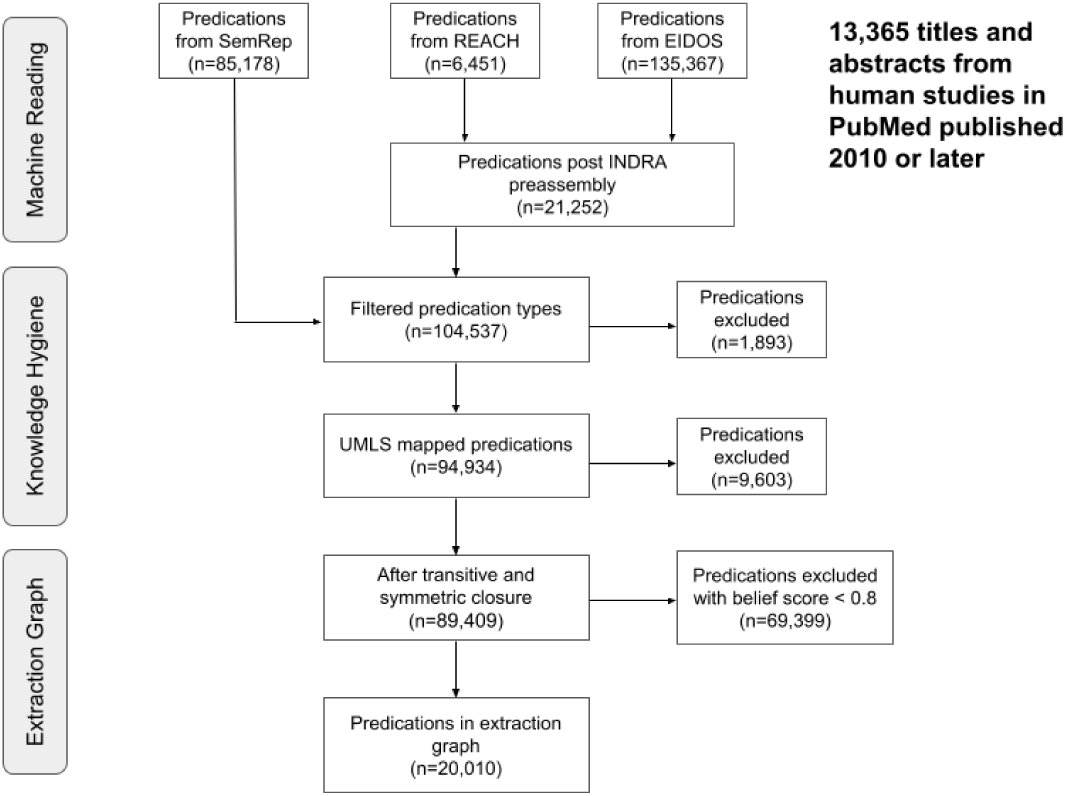
This figure illustrates merging the outputs from the machine reading systems, performing knowledge hygiene, and applying logical closure operations on the extraction graph.

The KG from the PheKnowLator workflow contained 18,869,980 triples. *Table 3* shows the counts of inferred edges by predicate type. The most significant gains by predicate type `causes_or_contributes_to_condition` yielded the most novel inferred causal edges, followed by `causal_relation_between_processes`. There were zero inferred edges for certain other predicate types because we did not define the logical properties of those predicates to allow for inference. *Table 4* shows the count of edges for four triple types relating genes or variants of genes to health conditions.

**Table 3.**
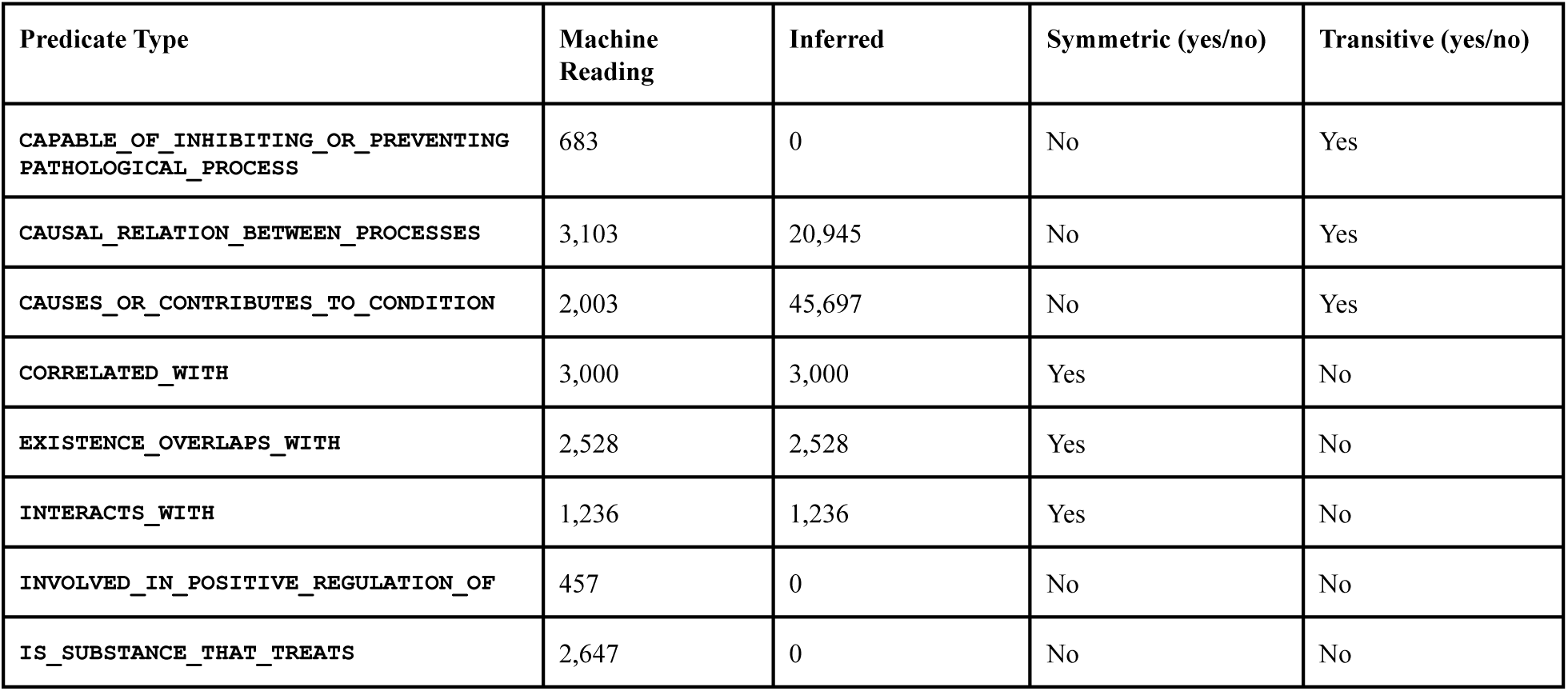
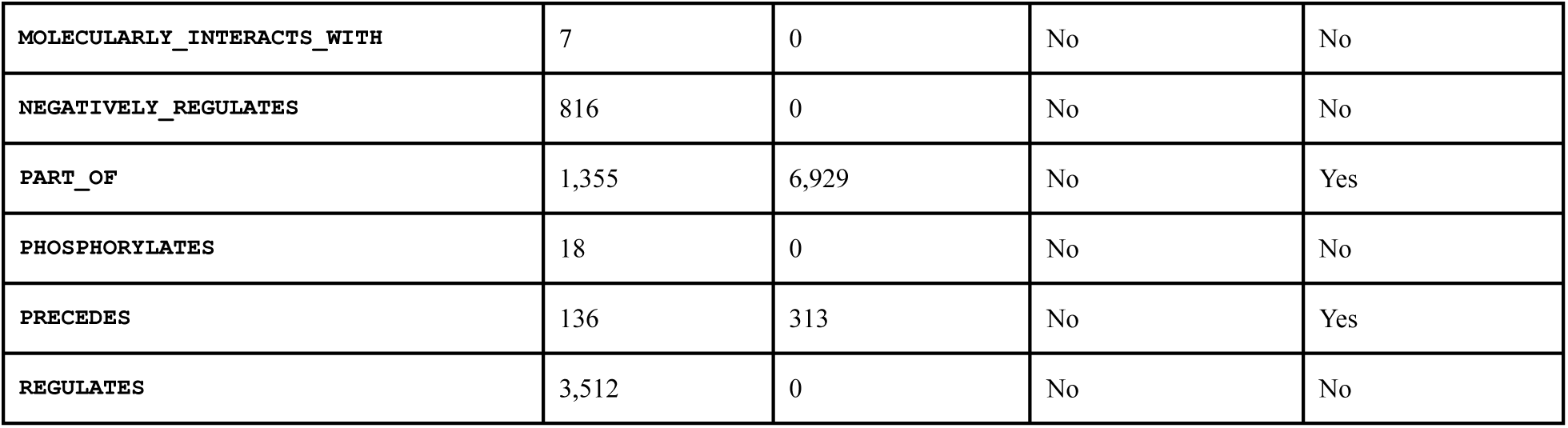
Counts of triples inferred using the logical closure operations given the symmetric and transitive properties of the literature-derived predicates from machine reading. Transitive and symmetric closure produced 73,046 triples after filtering out inferred triples with a belief score of < 0.6 and those without a mapping to the RO.

| Predicate Type | Machine Reading | Inferred | Symmetric (yes/no) | Transitive (yes/no) |
| --- | --- | --- | --- | --- |
| `CAPABLE_OF_INHIBITING_OR_PREVENTING_PATHOLOGICAL_PROCESS` | 683 | 0 | No | Yes |
| `CAUSAL_RELATION_BETWEEN_PROCESSES` | 3,103 | 20,945 | No | Yes |
| `CAUSES_OR_CONTRIBUTES_TO_CONDITION` | 2,003 | 45,697 | No | Yes |
| `CORRELATED_WITH` | 3,000 | 3,000 | Yes | No |
| `EXISTENCE_OVERLAPS_WITH` | 2,528 | 2,528 | Yes | No |
| `INTERACTS_WITH` | 1,236 | 1,236 | Yes | No |
| `INVOLVED_IN_POSITIVE_REGULATION_OF` | 457 | 0 | No | No |
| `IS_SUBSTANCE_THAT_TREATS` | 2,647 | 0 | No | No |
| MOLECULARLY_INTERACTS_WITH | 7 | 0 | No | No |
| NEGATIVELY_REGULATES | 816 | 0 | No | No |
| PART_OF | 1,355 | 6,929 | No | Yes |
| PHOSPHORYLATES | 18 | 0 | No | No |
| PRECEDES | 136 | 313 | No | Yes |
| REGULATES | 3,512 | 0 | No | No |

**Table 4.** The count of four causal predicates in the KG produced by PheKnowLator after running OWL-NETS.

| Predicate Type | # of triples |
| --- | --- |
| CAUSES_OR_CONTRIBUTES_TO_CONDITION | 113,837 |
| MOLECULARLY_INTERACTS_WITH | 5,678,829 |
| GENETICALLY_INTERACTS_WITH | 3,246,417 |

Note that we did not collect detailed, precise runtimes for these analyses. All analyses were performed on modest hardware configurations. The execution of the INDRA machine reading component took several days. The post-processing of the SemRep and INDRA outputs took minutes. The construction of the knowledge graph took days. Finally, the execution of all of the SPARQL queries took several hours on the modest hardware.

### 3.1 Answering competency questions

#### 3.1.1 Question #1 - What are the potential confounders of Depression and AD?

The KG search found 126 unique variables potentially confounding the association between depression and AD, including 47 conditions or phenotypes, 43 drugs, and 35 genes or enzymes. Examples included apolipoproteins, arachidonic acid, chronic infectious disease, encephalopathies, estradiol, hypercholesterolemia, hypoglycemia, inflammatory response, interleukin-6, myocardial infarction, and non-alcoholic fatty liver disease. (The KG search found 141. Fifteen of the 141 confounders appeared to be redundant, e.g., *vascular endothelial growth factor A* (PR_000017284) and *vascular endothelial growth factor A (human)* (PR_P15692)). However, according to PRO, these concepts are not the same. The human version (PR_P15692) is a child of (PR_000017284), so we retained both in the KG. For the purposes of reporting in the tables, we have merged these concepts.

The shortest path lengths returned by Dijkstra’s path search algorithm varied from 2 (15 paths from a confounder to depression / 16 paths from a confounder to AD) to 9 (2 paths, both from confounder to depression or confounder to AD). The paths for the confounder with the longest path lengths (*statin* (CHEBI_87631)) included paths that passed through the hub *molecular entity* (CHEBI_23367). The majority of confounders had paths to depression or AD with 4 or fewer path lengths (77 paths with <= 4 steps to depression and 80 paths with <= 4 steps to AD). *Figure 5* depicts the reasoning path for inflammatory response.

**Figure 5.**
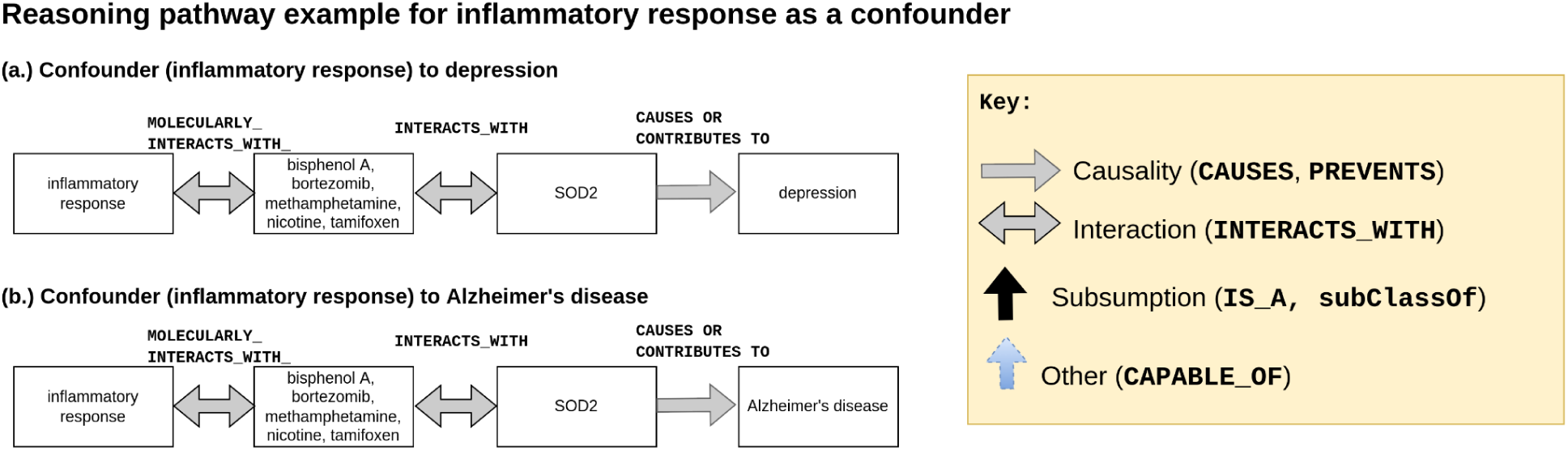
Sample reasoning pathway of inflammatory response, a confounder identified by KG search. Note that the y-axis is the **`type_of/`**subsumption axis. The x-axis is the **`causes/interacts_with`** axis. *SOD2* is superoxide dismutase 2, a gene that encodes a mitochondrial protein. *SOD2* is associated with diabetes and gastric cancer^94^. *SOD2* polymorphisms are associated with neurodegenerative diseases, including cognitive decline, stroke, and AD^95, 96^. *SOD2* is a gene that scavenges for reactive oxygen species (ROS) resulting from environmental exposures, e.g., bisphenol A, that result in inflammation^97^. Bisphenol A is found ubiquitously in plastics and is ingested from water bottles, packaged food, and many other sources^98–100^.

#### 3.1.2 Question #2 - What are potential colliders of Depression and AD?

The KG search found 28 unique potential colliders for depression and AD. These included amyotrophic lateral sclerosis, anemia, apraxias, atherosclerosis, brain hemorrhage, cerebral amyloid angiopathy, cerebrovascular accident, congestive heart failure, deglutition disorders, *<u>dementia</u>*, diabetes mellitus, diabetes mellitus, non-insulin-dependent, encephalitis, falls, *<u>frontotemporal dementia</u>*, ***homeostasis***, immune response, insulin resistance, ischemic stroke, malnutrition, neurofibrillary degeneration (morphologic abnormality), osteoporosis, parkinsonian disorders, pathogenesis, pneumonia, psychotic disorders, senile plaques, stroke, and tauopathies.

Note that in the list of colliders above, the KG search identified as potential colliders several variables we have emphasized by italicizing and underlining. The KG may have inferred dementia because AD is a specific dementia etiology. Homeostasis may have been included in the results as a generic biological process that was not filtered out. Finally, it would not make sense that AD would cause frontotemporal dementia, the most common dementia in patients younger than 60. We drop supercategories like dementia and generic processes like homeostasis for our analysis since these can be easily identified and explained away from context as spurious.

There were two colliders identified using SPARQL for which no path could be found from depression to the collider in the KG created using the PheKnowLator workflow (*falls* (HP_0002527) and *neurofibrillary tangles* (HP_0002185)). Two of the 26 colliders with paths from depression in KG had multiple identifiers (*cerebral amyloid angiopathy* (HP_0011970 and DOID_9246) and *frontotemporal dementia* (DOID_10216, DOID_9255, and HP_0002145)). Path search identified a total of 29 collider entities in KG.

For depression to the colliders, the path lengths of the shortest paths returned were 7 steps for 2 colliders, 8 for 26, and 9 for 1. The majority of colliders (24) resulted in 5 or more shortest paths returned. For AD to the colliders, the path step counts were slightly lower at 6 steps for 2 colliders, 7 for 26 colliders, and 8 for 1 collider. Similar to depression, 23 colliders have 5 or more shortest path lengths from AD.

As shown in *Figure 6* below, the typical path pattern for both cases involved passing through the hub node *nervous system* (UBERON_0001016) and *nervous system process* (GO_0050877) that molecularly interacts with a drug or chemical that also interacts with a gene that causes or contributes to AD.

**Figure 6.**
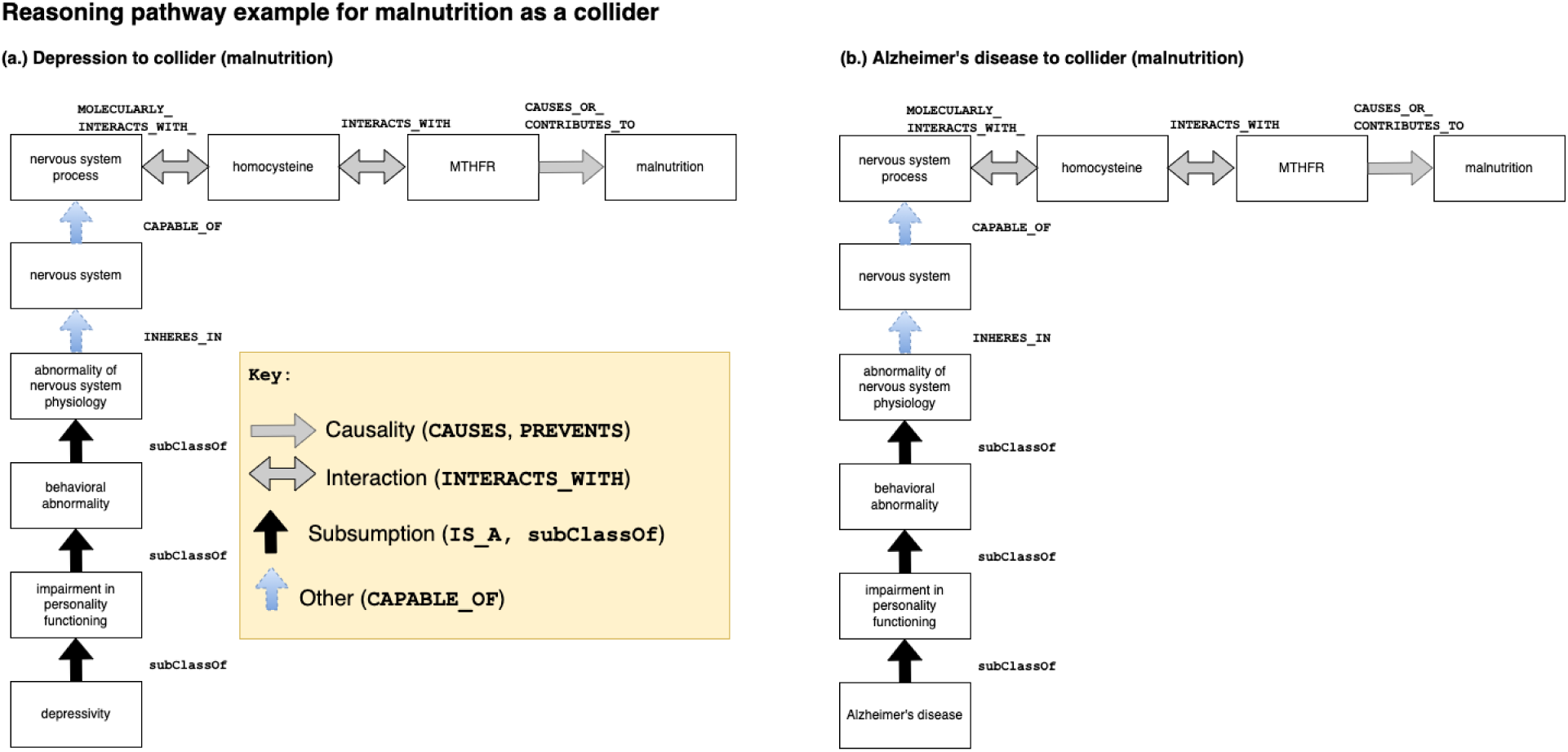
Sample reasoning pathway of malnutrition, a collider identified by KG search. The *MTHFR* gene is a gene that encodes the Methylenetetrahydrofolate reductase (MTHFR) enzyme that is critical for the metabolism of amino acids, including homocysteine^101^. *MTHFR* perturbations and mutations decrease the metabolism and elevate homocysteine. Elevated homocysteine is linked with blood clots and thrombosis events^102–104^. Homocysteine has been reported in the published literature as a confounder of depression and AD in at least one observational study^105^. Elevated homocysteine is also associated with low levels of vitamins B6 and B12 and folate.

#### 3.1.3 Question #3 - What are the potential mediators of Depression and AD?

The KG search found 18 unique potential mediators that were conditions or phenotypes for depression and AD. These included amyotrophic lateral sclerosis, anemia, atherosclerosis, cerebral amyloid angiopathy, cerebral hemorrhage, congestive heart failure, *<u>dementia</u>*, diabetes mellitus, dysphagia, insulin resistance, ischemic stroke, malnutrition, osteoporosis, parkinsonism, pathogenesis, stroke, tauopathy, type II diabetes mellitus. The shortest path lengths from Dijkstra’s path search algorithm were 8 for depression to mediators. There is more variety in path length from mediators to AD, with most path lengths between 4 and 6. As is shown in *Figure 7* below, the typical path pattern for both cases involves associating the disorder (depression or a mediator) with a nervous system process that molecularly interacts with a chemical that causes or contributes to AD.

**Figure 7.**
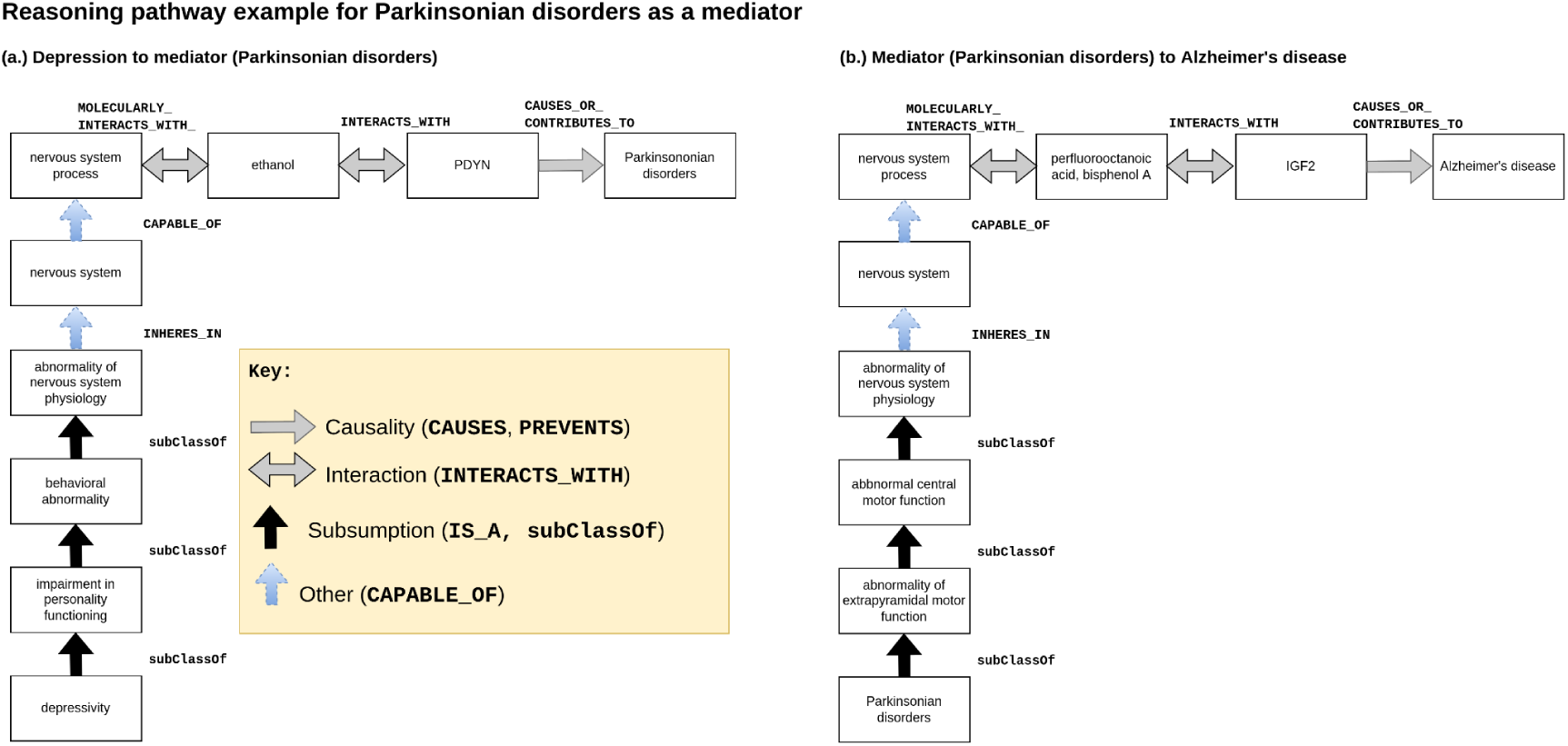
Sample reasoning pathway of Parkinsonian disorders, a potential mediator for AD identified by KG search. Prodynorphin (*PDYN*) is a gene that produces endogenous opioid peptides involved with motor control and movement^106^ implicated in neurodegenerative diseases that modulate response to psychoactive substances, including cocaine and ethanol^107^. The gene insulin growth factor 2 (*IGF2*) is associated with development and growth. Perfluorooctanoic acid and bisphenol A exposure is associated with neurodegeneration resulting in AD^108^. Hippocampal *IGF2* expression is decreased in AD-diagnosed patients^109^. Increased *IGF2* expression improves cognition in AD mice models^110^.

### 3.2 Analysis of search results by causal role

#### 3.2.1 Simple roles

*Figure 8* illustrates the phenotype variables produced from our KG search. Aside from the conditions we analyzed, the KG search also identified pharmaceutical substances, genes, and enzymes as confounders. However, the shortest path parameters produced no search results from our queries for colliders and mediators belonging to those semantic categories. Note also the same pattern was observed searching SemMedDB. We report the complete set of variables the KG identified in the spreadsheet in the Supplementary Materials in **File III.**, **Worksheet S2,** and **Worksheet S3**. (see the worksheets entitled “Substances” and “GenesEnzymes,” respectively), and in **File II. Appendix B. Figures S1 and S2.**

**Figure 8.**
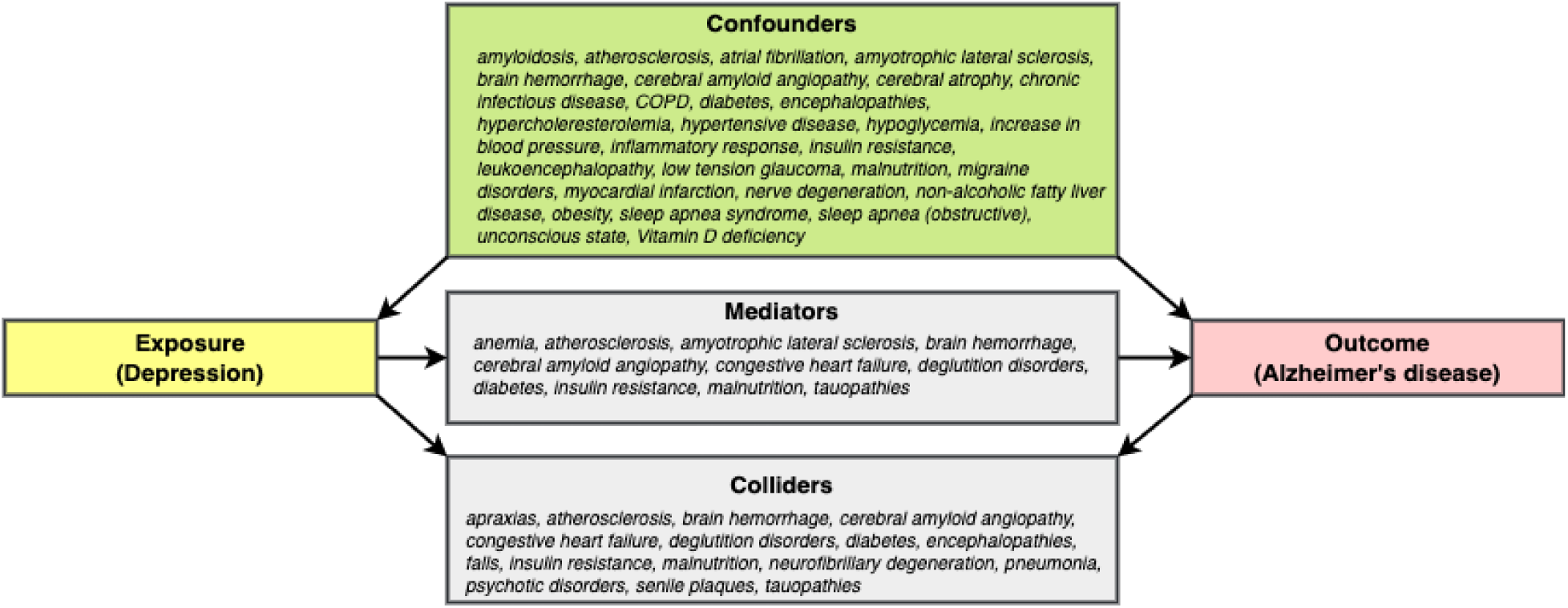
This diagram shows the variables identified using their causal roles relating to depression and AD by translating standard epidemiological definitions into patterns for querying the KG.

#### 3.2.2 Complex roles

*Figure 9* illustrates the same conditions from *Figure 8*, albeit in a more distilled form since we distinguish between variables playing single versus the different categories of combination roles. Twenty-two distinct confounders and nine colliders were found to behave solely in one role. The KG search could not find any variables that behave exclusively in the role of mediator. The rest of the variables fulfilled two or more causal roles. Of these, 11 variables fulfilled the criteria of all three causal roles.

**Figure 9.**
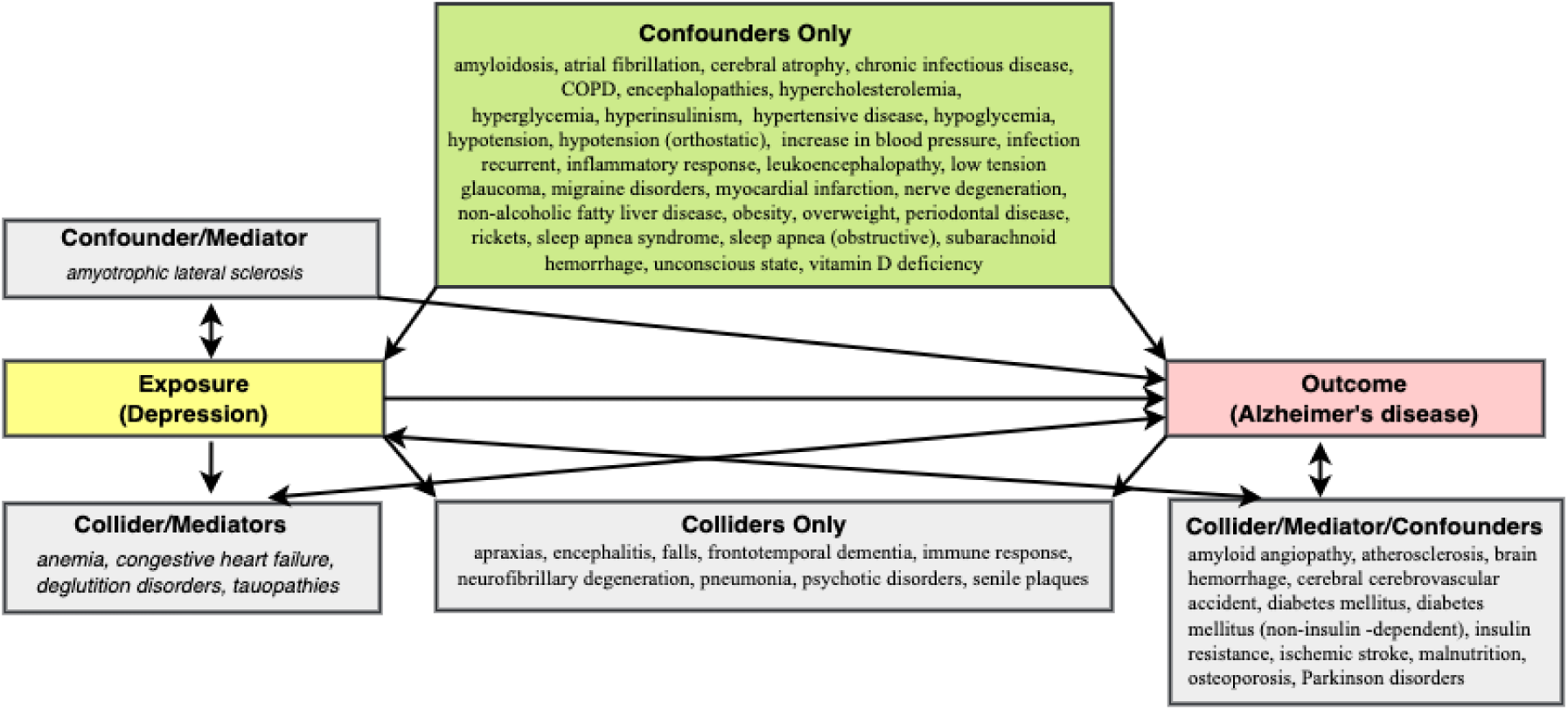
This diagram shows variables according to their roles, including single and hybrid role types for the relationship between depression (exposure) and AD (outcome). Each rectangular box in the figure lists all the conditions by single role (confounder, collider, or mediator) or combination roles (e.g., mediator/collider). For example, the box labeled “Confounders only” contains covariates that are exclusively confounders and were not found to act as a collider or a mediator.

Note that the graphs in *Figures 8* and *9* are not, strictly speaking, causal DAGs since they depict cycles. Since causal DAGs cannot represent cyclic relationships, new graph formalisms can represent latent variables and cycles (feedback loops) by generalizing the causal interpretation of DAGs, e.g., directed mixed graphs (DMGs)^111^, chain graphs^112^.

### 3.3 KG vs. SemMedDB and reported covariates in the published literature

*Table 5* shows the confounders from the pilot meta-review of reported confounders we gathered from human observational studies investigating depression as the exposure and dementia or Alzheimer’s disease as the outcome. About half of the studies were prospective cohort studies. The other study designs used were case-control and using twins for controls. We looked at studies at AD-specific studies, to see if there were differences between those used mainly from dementia-focused studies below, but could not find any major differences.

**Table 5.**
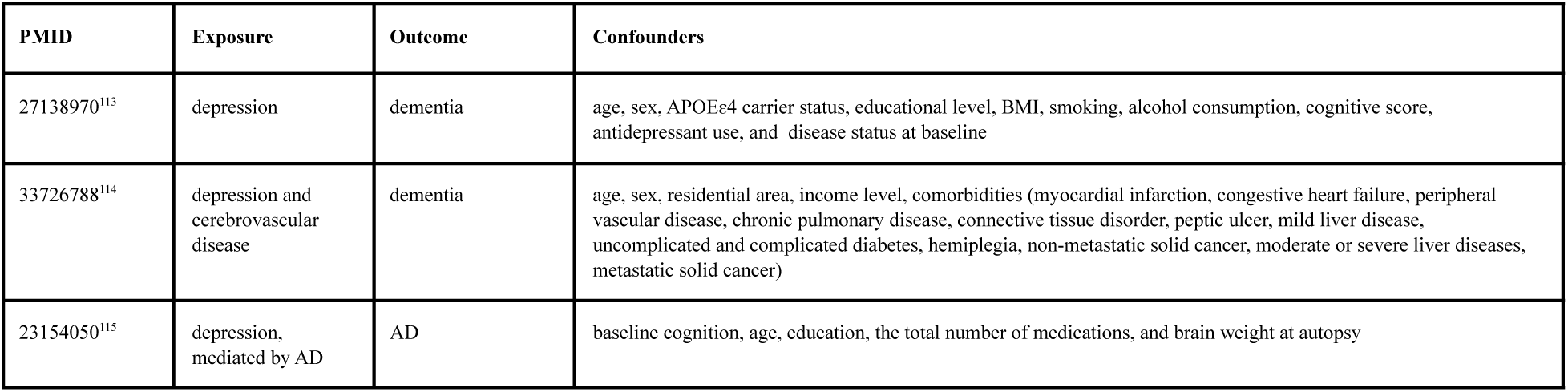
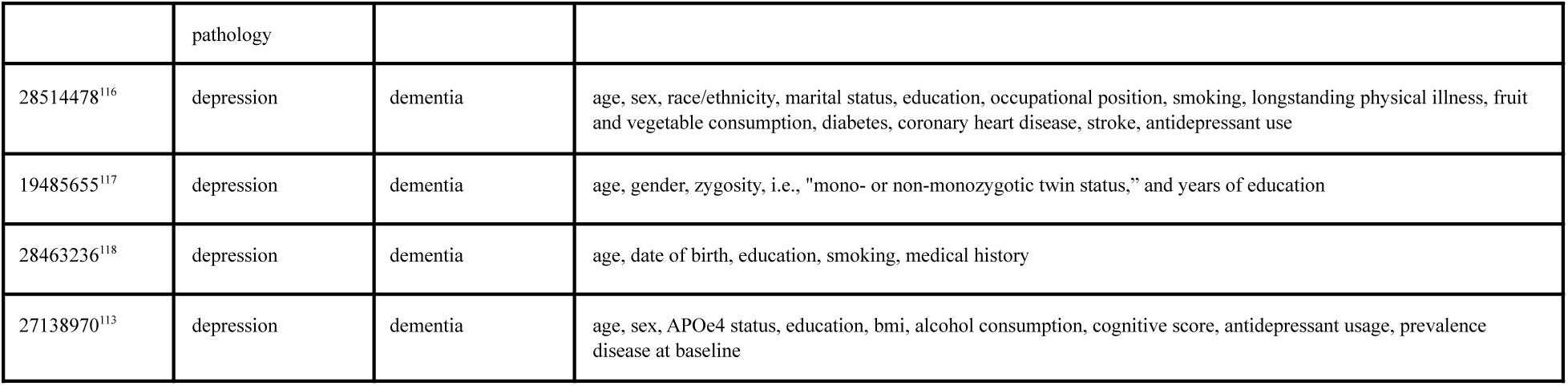
Reported confounders from published studies investigating depression and AD (or dementia) from our pilot meta-review.

| PMID | Exposure | Outcome | Confounders |
| --- | --- | --- | --- |
| 27138970<sup>113</sup> | depression | dementia | age, sex, APOEε4 carrier status, educational level, BMI, smoking, alcohol consumption, cognitive score, antidepressant use, and disease status at baseline |
| 33726788<sup>114</sup> | depression and cerebrovascular disease | dementia | age, sex, residential area, income level, comorbidities (myocardial infarction, congestive heart failure, peripheral vascular disease, chronic pulmonary disease, connective tissue disorder, peptic ulcer, mild liver disease, uncomplicated and complicated diabetes, hemiplegia, non-metastatic solid cancer, moderate or severe liver diseases, metastatic solid cancer) |
| 23154050<sup>115</sup> | depression, mediated by AD pathology | AD | baseline cognition, age, education, the total number of medications, and brain weight at autopsy |
| 28514478<sup>116</sup> | depression | dementia | age, sex, race/ethnicity, marital status, education, occupational position, smoking, longstanding physical illness, fruit and vegetable consumption, diabetes, coronary heart disease, stroke, antidepressant use |
| 19485655<sup>117</sup> | depression | dementia | age, gender, zygosity, i.e., "mono- or non-monozygotic twin status," and years of education |
| 28463236<sup>118</sup> | depression | dementia | age, date of birth, education, smoking, medical history |
| 27138970<sup>113</sup> | depression | dementia | age, sex, APOE4 status, education, bmi, alcohol consumption, cognitive score, antidepressant usage, prevalence disease at baseline |

While demographic variables such as age and sex are consistently reported and there is considerable overlap between some of the covariates including APOEɛ4 status and comorbidities related to kidneys, liver, and cardiovascular systems, there is inconsistency between studies, presumably depending on whether or not data were available for these variables, a common pattern in observational studies^119, 120^. Note that the second study in the third row of Table 5, with the many types of cancer listed as confounders, is something of an outlier.

We omitted columns for the INDRA machine reading systems (EIDOS and REACH) in *Figure 10* because there were no variables identified using information from those sources.

**Figure 10.**
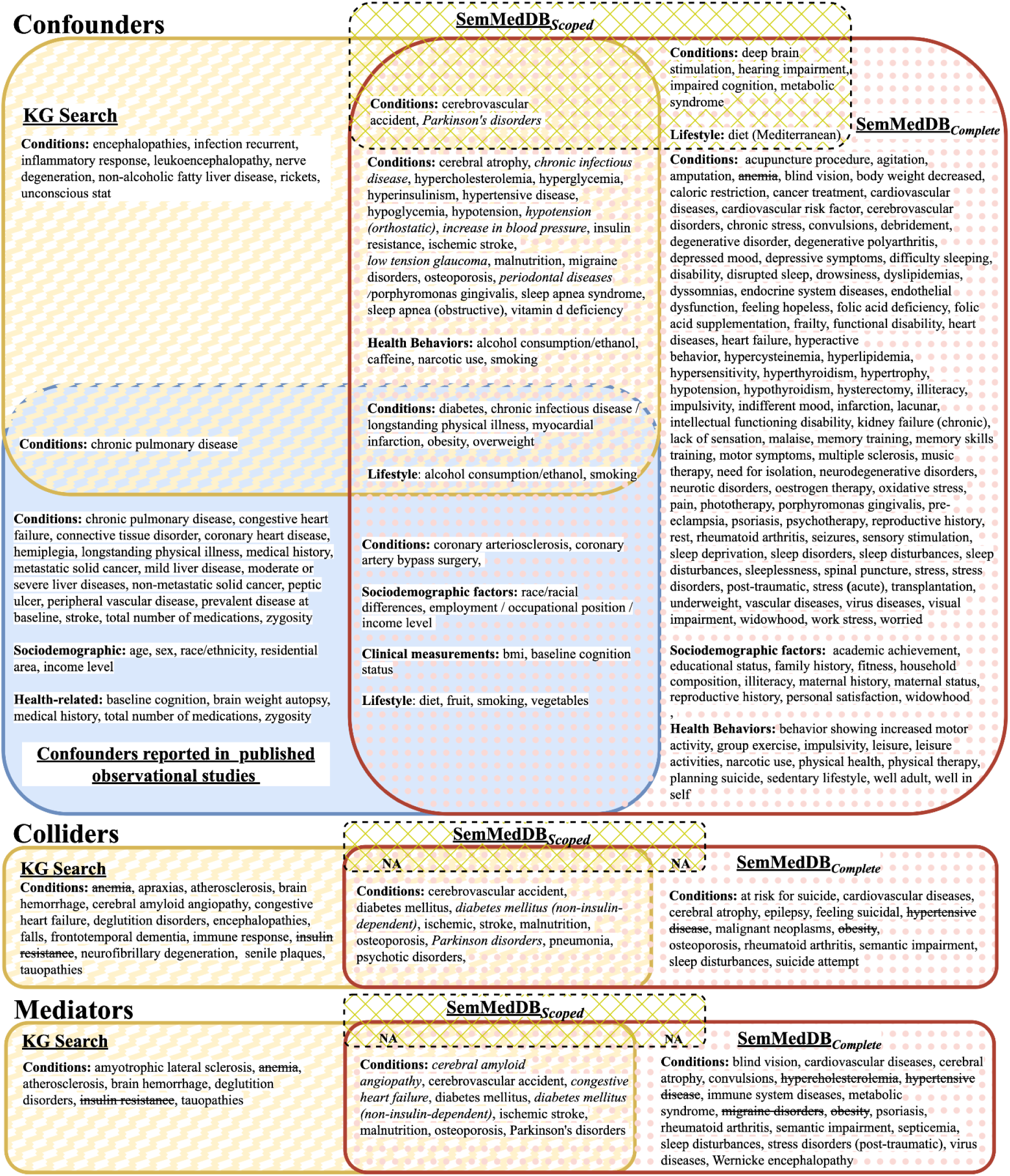
This figure shows (main) condition variables/concepts for simple (not combined) causal roles from KG search in yellow, SemMedDB_Scoped_ in green diagonal and SemMedDB_Complete_ in rose polka dot, and the confounders from the pilot-metareview in solid blue as a Venn diagram. See **Appendix B. Figure SI.** for the complete results, containing genes/enzymes/proteins and drugs/substances.

Only a small number of confounders appear in all sources, though many confounders appear in two sources. We see a moderately high concurrence between the KG and SemMedDB_Complete_, but there are curious and perhaps telling patterns. For instance, among the discordant results, there is a pattern that concerns the semantic categories of output. Specifically, SemRep had better recall performance for variables that are potentially more difficult to establish mechanistically, e.g., psychiatric symptoms (“agitation,” “dyssomnias,” “feeling hopeless”) or lifestyle factors (e.g., exercise, family history). Moreover, the concepts identified by KG search results are generally less redundant than those from searching the literature (e.g., diabetes, diabetes mellitus). Searching SemMedDB_Complete_ yielded the most search results. Searching the KG yielded fewer search results. Searching MachineReadingDB_Scoped_ yielded still fewer results. This is not surprising since the KG was harmonized while SemMedDB_Complete_ search was not, resulting in some concepts with multiple names.

Again, only a small number of confounders appear in all sources. Two primary patterns that stand out in *Figure 11* are the relatively high degree of concordance between the KG and SemMedDB_Complete_ regarding confounders and combination variables that play all three roles (confounder, collider, and mediator) on the one hand. Note that while there is a literature describing the three “simple” or core roles (confounder, collider, and mediator), the literature for confounder/mediator (also called time-varying confounding and treatment-confounder feedback) is the only type of combination or hybrid variable for which a literature exists^14^. From this point on, we call variables playing all three roles, such as stroke, chimeras. In Greek mythology, a chimera is a creature with a fire-breathing head of a lion, a goat’s body, and a snake’s tail. From the perspective of valid causal inference, the chimera is a dangerous variable category because conditioning on chimeras, like conditioning on mediators and colliders, can potentially distort measures of association or effect. Thus, chimeras are important to identify to avoid introducing bias. On the other hand, the recall performance for the other types of exclusive and the combination variable is poor. For example, anemia appears in “Mediator only” in the KG search results but in “Confounders only” in the SemMedDB_Complete_ search results.

**Figure 11.**
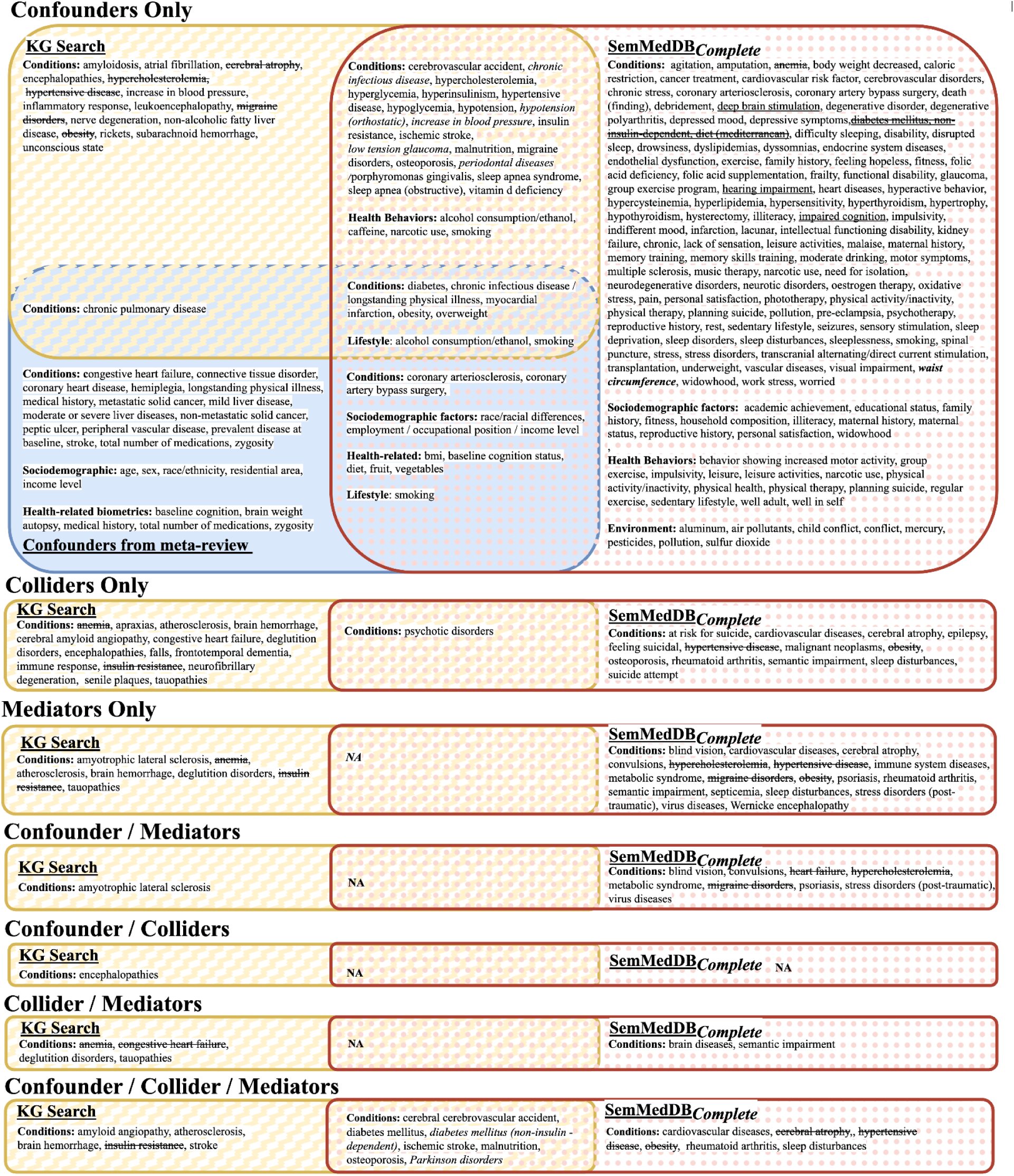
This figure shows the (mainly) condition variables/concepts for combination causal roles from KG search in yellow, SemMedDB_Complete_ in pink poka dot, and the confounders from the pilot-metareview in solid blue as a Venn diagram. See **Appendix B. Figure S2.** for the complete results, containing genes/enzymes/proteins and drugs/substances.

### 3.4 Manual qualitative evaluation of search results

In this section, we analyze five condition variables in-depth. We prioritized conditions for in-depth analysis for the following reasons. First and foremost, as noted before, conditions were the only category of biomedical entity variable identified as playing all three causal roles in both the KG search results and in SemMedDB search.

Hence, phenotypes were the only category of variable we could analyze from the standpoint of untangling the complexity of combined causal roles using the reasoning path outputs. Secondly, it would have been infeasible to analyze all the variables. We had to prioritize a subset of variables. Thirdly, conditions were the most frequently reported category of reported confounder in published observational studies after sociodemographic characteristics, health behaviors, and APOEε2/3/4 status. Fourthly, our use case focused on variables that would be commonly available in an electronic medical records system, which ruled out biomarkers.

#### 3.4.1 Confounders

The objective of this section was to provide an in-depth analysis of the evidence supporting the classification of a sample of potentially newly discovered confounders for depression and AD. The source sentences for review of three variables identified as being “confounders only” in both the KG and SemMedDB_Complete_: chronic infectious disease (CID), obstructive sleep apnea (OSA), and vitamin D deficiency (VDD) are available in the Supplementary Materials in **File V. *ConfounderSourceSentences.xlsx***. (Note that SQL code for running the queries with which to obtain sources sentences from SemMedDB is included in the final worksheet of **File V.**)

These variables were chosen for analysis because: (1.) they occurred in both KG and SemMedDB_Complete_, and the agreement was unanimous about their categorization (confirmation by two distinct methods); (2.) they had the most support in SemMedDB, and we could analyze the source sentences and publications supporting their categorization as confounders (amount of evidence, i.e., the # of retrieved and pertinent predications); and (3.) the practical usefulness for our use case, scientific interest, and surprise. In this case, OSA, CID, and VDD pertained directly to our use case, and had support in SemMedDB, yet, surprisingly, they did not appear in the published observational studies we examined, though our review was not comprehensive.

In the view of the content expert, all three variables seemed biologically plausible common causes of both depression and AD. OSA seemed to have the most (10 predications from 9 distinct publications supporting OSA→depression (where “→” means “**`causes`**”) and 22 predications from 21 distinct publications supporting OSA→AD) and the strongest evidence browsing PubMed. The 3 strongest “OSA→depression” predications and the 2 strongest “OSA→AD” predications are marked in the spreadsheet.

While there is a larger body of evidence on infections and AD, the evidence was less extensive overall for vitamin D deficiency (2 predications from 2 distinct publications supporting CID→depression and 4 predications from 4 distinct publications supporting CID→AD).

The sentences concerning VDD also supported it as a confounder (3 predications from 3 distinct publications supporting CID→depression and 9 predications from 9 distinct publications supporting CID→AD).

Notably, however, the expert noted the difficulty in assessing the evidence quality from one sentence, and the need to drill down into the article to determine if the causal assertion was transferable. When reviewing the OSA → MDD and OSA → AD assertions, the expert noted that the various studies were all plausible and included primary studies (e.g., OSA to MDD: supported by PMID 20939075, 27631236, 33158487, 33158487 and OSA→AD supported by PMID 27070140, 28329084). In general, OSA is a well-known risk factor for both MDD and AD both clinically and in research.

#### 3.4.2 Stroke as a chimeric variable

The source sentences from SemMedDB for stroke are included in the Supplementary Materials in **File VI. *ChimeraSourceSentences.xlsx***. (Note that SQL code for running the queries to obtain the source sentence from SemMedDB is included in the final worksheet of **File VI.**)

Stroke was selected for further analysis in this study due to its frequent appearance as a confounder in reported observational studies. Despite being commonly considered a confounder without referencing stroke-subtype (e.g., see the phenotype definition for stroke from Singh-Manoux et al.^116^), both the KG and SemMedDB_Complete_ search results indicated that the role of stroke might vary depending on its subtype, specifically between ischemic and hemorrhagic. These findings were further supported by the results of a PubMed search, which highlighted the potential difficulties in conditioning on stroke without considering its subtype.

##### Listing 2. Common ICD-10 codes used for stroke.

I60: Nontraumatic subarachnoid hemorrhage

I61: Nontraumatic intracerebral hemorrhage

I62: Other and unspecified nontraumatic intracranial hemorrhage

I63: Cerebral infarction

I64: Stroke, not specified as haemorrhage or infarction

Stroke was identified as a chimera by both the KG and SemMedDB_Complete_. The support for stroke as a chimera was substantive, though with a caveat where the interpretation of the meaning of stroke depends on the stroke subtype. To review the meaning of stroke, a stroke results from a reduced blood supply to the brain from either constriction, e.g., ischemic stroke, or hemorrhage, e.g., hemorrhagic stroke, or intracerebral hemorrhage. There are also less immediately devastating stroke-like events called transient ischemic attacks that presage future more serious stroke events. Ischemic stroke is a known risk factor for both depression^121–123^ and AD^124–126^. Although the KG search classified both stroke and the ischemic stroke subtype as chimeras, presumably including both subtypes under the umbrella term or hypernym of stroke, the evidence was stronger for ischemic stroke as a chimera. However, evidence for AD as a risk factor was weaker for ischemic than for hemorrhagic stroke^125^. In the meta-analysis of Waziri et al. based on four national studies and assessing the relative risk of hemorrhagic vs. ischemic stroke in AD vs. non-AD patients and applying matching criteria, the relative risk of AD patients vs. similar non-AD controls was 1.41 (95% CI 1.23-1.64) for hemorrhagic stroke but 1.15 (95% CI 0.89-1.48) for ischemic stroke. A PubMed search did not yield any specific information about the hemorrhagic stroke subtype as a confounder.

#### 3.4.3 Resolving contradictions between SemMedDB and KG search

Anemia was selected for further analysis in this study due to its complex and contradictory relationship with depression and Alzheimer’s disease. The supporting evidence from reasoning paths, source sentences, and PubMed searches was contradictory, making it challenging to determine the exact causal role of anemia. This complexity and contradiction provided an opportunity to achieve one of the sub-goals of the paper (**H3**), which was to examine the causal roles of identified variables through the interpretation and evaluation of reasoning paths.

Anemia is scientifically interesting as it has been reported to have complex relationships with both depression and Alzheimer’s disease. While anemia has been reported as a risk factor for depression, some studies have also suggested that depression may lead to anemia. Similarly, while anemia has been reported to increase the risk of Alzheimer’s disease, other studies have suggested that Alzheimer’s disease may lead to anemia.

In order to provide a more nuanced understanding of the relationship between anemia and depression, and Alzheimer’s disease, it was important to examine anemia in detail. This examination allowed for the exploration of the reasoning paths and the evaluation of the causal roles of anemia, depression, and Alzheimer’s disease, thereby contributing to the overall goal of the paper (**H3**) to provide mechanistic explanations of the causal roles of identified variables.

As noted previously in Results, anemia, a disease characterized by low hemoglobin levels, appears as a “mediator/collider” in the KG results but exclusively as a confounder in the SemMedDB_Complete_ results. Table 6 shows the support from SemMedDB for anemia causing depression was provided by two **`predisposes`** triples, and the support for anemia causing AD was supplied by a single **`causes`** triple. In terms of conveying causative force, **`predisposes`** is weaker than **`causes`** (see Rindflesch et al.^127^ for details) and implies that anemia may attenuated as a risk factor, or not well studied.

**Table 6.**
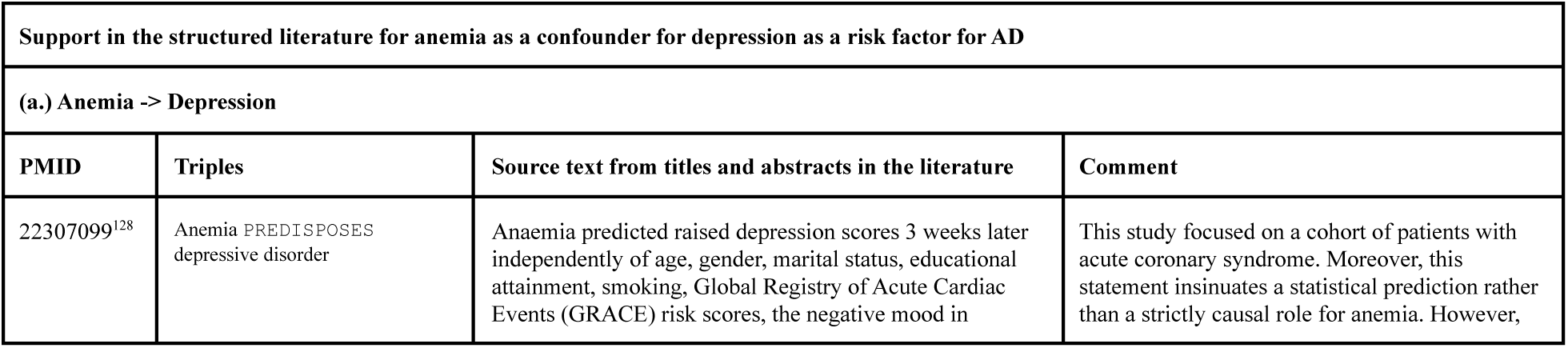
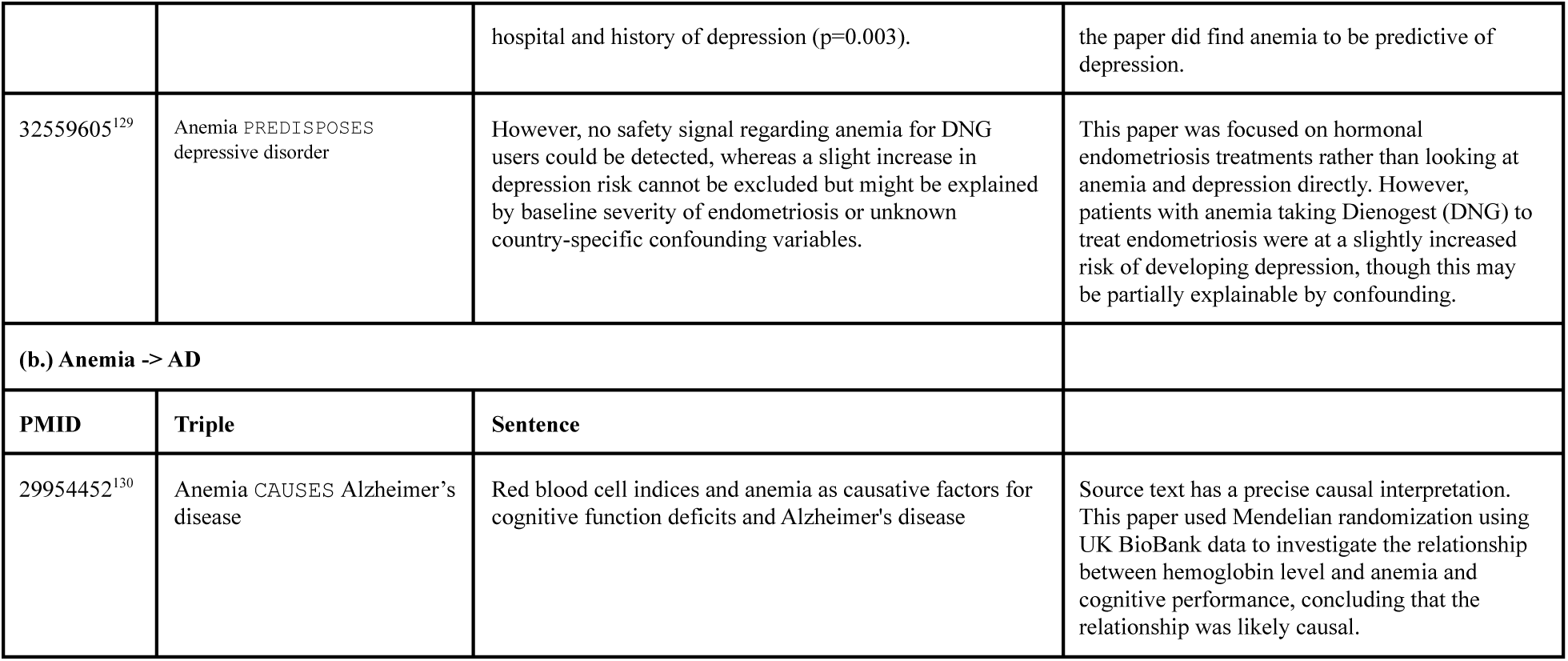
Support from SemMedDB for anemia as a confounder of depression as a risk factor for AD. The support is divided into two components: **(a.)** the anemia→ depression component and **(b.)** the anemia → AD component.

| Support in the structured literature for anemia as a confounder for depression as a risk factor for AD |  |  |  |
| --- | --- | --- | --- |
| (a.) Anemia -> Depression |  |  |  |
| PMID | Triples | Source text from titles and abstracts in the literature | Comment |
| 22307099<sup>128</sup> | Anemia **PREDISPOSES** depressive disorder | Anaemia predicted raised depression scores 3 weeks later independently of age, gender, marital status, educational attainment, smoking, Global Registry of Acute Cardiac Events (GRACE) risk scores, the negative mood in | This study focused on a cohort of patients with acute coronary syndrome. Moreover, this statement insinuates a statistical prediction rather than a strictly causal role for anemia. However, |
|  |  | hospital and history of depression (p=0.003). | the paper did find anemia to be predictive of depression. |
| 32559605<sup>129</sup> | Anemia PREDISPOSES depressive disorder | However, no safety signal regarding anemia for DNG users could be detected, whereas a slight increase in depression risk cannot be excluded but might be explained by baseline severity of endometriosis or unknown country-specific confounding variables. | This paper was focused on hormonal endometriosis treatments rather than looking at anemia and depression directly. However, patients with anemia taking Dienogest (DNG) to treat endometriosis were at a slightly increased risk of developing depression, though this may be partially explainable by confounding. |
| **(b.) Anemia -> AD** |  |  |  |
| **PMID** | **Triple** | **Sentence** |  |
| 29954452<sup>130</sup> | Anemia CAUSES Alzheimer's disease | Red blood cell indices and anemia as causative factors for cognitive function deficits and Alzheimer's disease | Source text has a precise causal interpretation. This paper used Mendelian randomization using UK BioBank data to investigate the relationship between hemoglobin level and anemia and cognitive performance, concluding that the relationship was likely causal. |

*Figures 12* and *13* present the support from the KG for anemia as a collider and mediator, respectively, of the depression and AD relationship.

**Figure 12.**
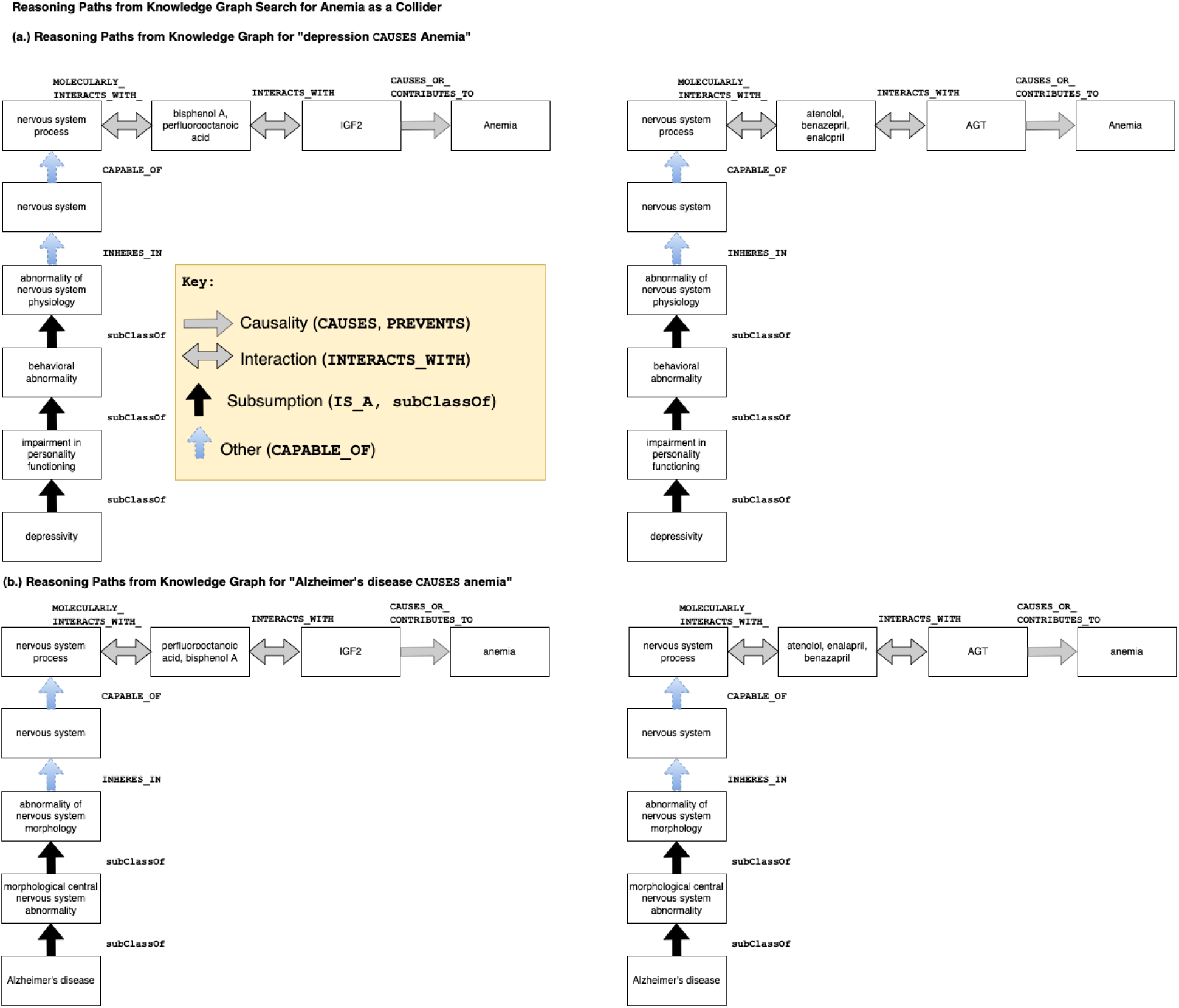
This figure shows the reasoning path support for depression causes anemia in (a.) and reasoning path support for AD causes anemia in (b.), and shows the mediating role of IGF2 and AGT in the relationship between depression, AD, and anemia. Reasoning path support in the KG for anemia as a common effect or collider for depression and AD. Both reasoning paths on the left **(a.)** and **(b.)** show how *IGF2* mediates between depression and environmental exposures (e.g., perfluorooctanoic acid and bisphenol A) on the one hand and anemia on the other. Angiotensinogen (*AGT*) is the name of a gene and enzyme coded by that gene involved in blood pressure and maintaining fluid and electrolyte homeostasis^131^. Both reasoning paths on the right **(a.)** and **(b.)** show how *AGT* mediates between AD and the drugs atenolol (a beta-blocker) and benazepril and enalapril are angiotensin-converting enzyme inhibitors, or (ACEIs), on the one hand, and anemia on the other. These drugs are known to cause anemia^132^. Numerous studies indicate AD as a potential cause of anemia^133, 134^.

**Figure 13.**
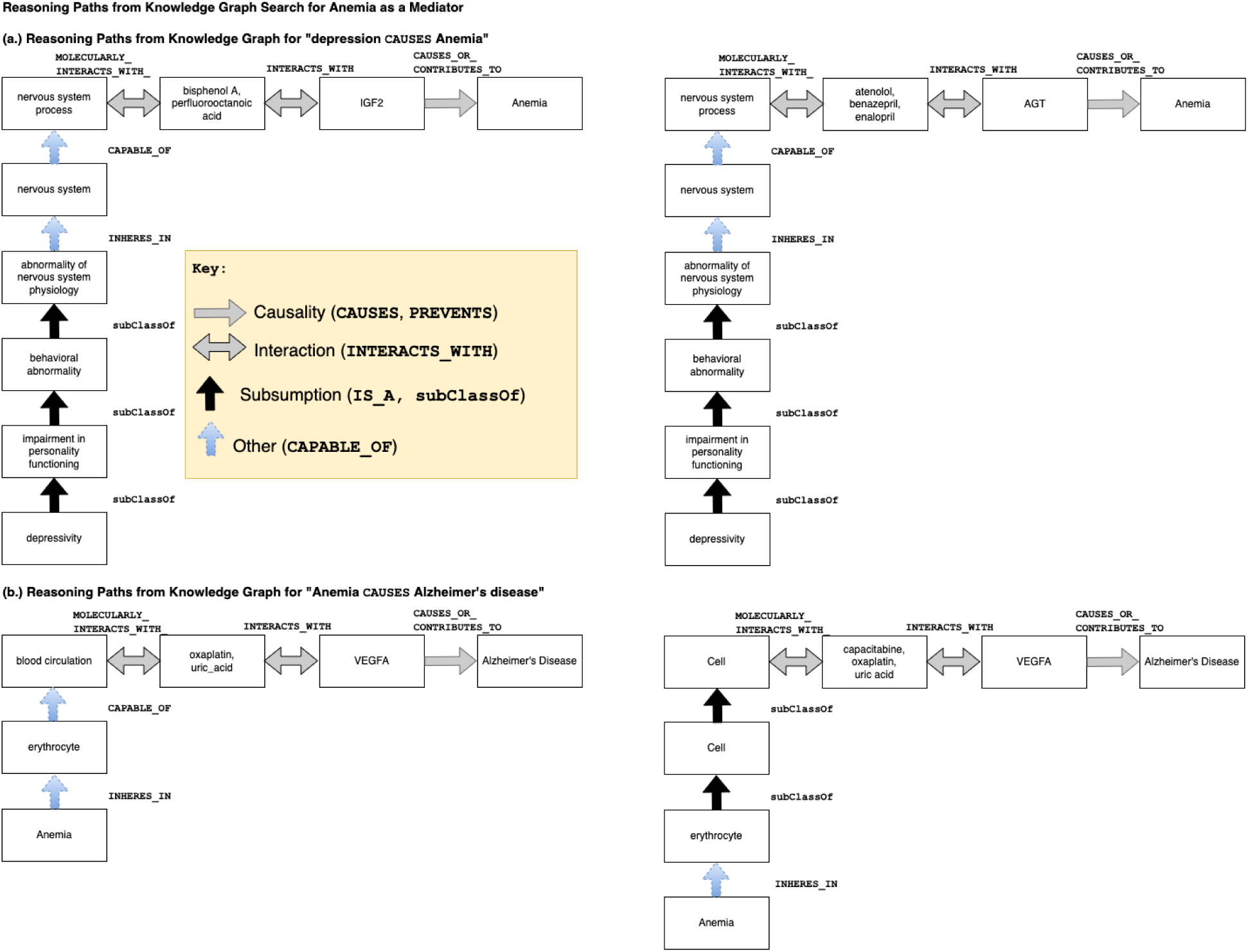
Support in the KG for anemia as a mediator for depression and AD. Note that **(a.)** contains the same information as Figure 12 **(a.)**. Vascular endothelial growth factor A (*VEGFA*) promotes the proliferation and migration of vascular endothelial cells and is upregulated in tumorogenesis^135^. *VEGFA* is also upregulated in AD^136^.

As shown in *Figure 13*, anemia is indeed in a complex relationship. Bisphenol A, an endocrine disruptor and an everyday exposure, disrupts the iron cycle. Low iron decreases dopamine which causes low mood, and depression causes brain atrophy associated with cognitive decline. Tentatively, our verdict is that anemia is a potential confounder and a mediator, though the evidence provided is not very strong.

Searching PubMed, we found support for AD causing anemia with an adjusted OR = 3.41 (95% CI 1.68, 6.92) reported in Faux et al.^133^, and was explained mechanistically with decreased plasma iron levels in AD^134^. We also found support for anemia causing AD. A recent paper using a robust study design explored the relationship between anemia and AD^137^. The potential causal relationship between anemia and depression is well known^138, 139^. Most surprisingly, we found support for depression causing anemia. Vulser et al. found a dose-response relationship between depression severity and anemia^140^.

## 4. Discussion

Our findings suggest that our novel causal feature selection method can identify plausible confounder, collider, and mediator variables for depression and AD (**H1**). A substantial number of variables, all conditions, were found to play multiple roles. Surprisingly, only conditions appeared in all three roles. The extent to which specific candidate confounders could behave in unexpected ways that violate the strict definition of confounding further confirms the complexity of variables required for studying the relationship between depression and AD (**H2**). KG search produced interpretable mechanistic explanations in the form of reasoning paths (**H3**). Finally, the KG and SemMedDB_Complete_ search results did not always agree about either the variables or the roles. There were potentially essential confounders identified by human content-matter experts that were not identified by our method. Note that SemMedDB is not a reference standard and that the status of the variables identified can only be determined by expert review. The clinical significance of identifying as yet unknown confounders is that these confounders may as yet be additional modifiable risk factors. For example, chronic infectious disease may be intervened upon via adult vaccination. There is a growing literature using large, retrospective reviews that adults vaccinated against common infectious agents (herpes zoster, hepatitis, tuberculosis) have a 25-45% lower risk of dementia^141–145^. The notion is that adult vaccination “tunes up” the immune response to whatever causes AD, even if it is non-specific. The risk reduction is robust, and from large reviews that appear to have been corrected for SES status, health care, and education. Vitamin D deficiency and sleep apnea are also amenable to treatment and could independently modify AD risk.

In contrast to systems based on assertions from the literature, where source sentences provide human-auditable provenance, the explanatory capability of KG search is provided by reasoning paths. The reasoning paths provide supporting evidence by containing a trace of the nodes and predicates transverse in the course of searching the KG executing the SPARQL query. The reasoning pathway thus furnishes a mechanistic explanation that is, in essence, a hypothesis or theory justifying the causal role(s) identified for that variable, filtered through ontology-grounded resources.

The reasoning paths, which informally represent mechanistic hypotheses, seemed biologically plausible upon inspection. Although there were problems with the reasoning paths from including generic nodes, we felt it had a strong explanation capability that could adequately clarify how variables were inferred from the source ontologies.

A review by a human expert in AD confirmed that the variables identified by both the KG and SemMedDB_Complete_, including chronic infectious disease, obstructive sleep apnea, and vitamin D deficiency, were likely confounders. However, in our inexhaustive, exemplary review of reported covariates from observational studies investigating depression and AD, these variables were not included. We interpret this to mean that the causal feature selection method achieved a major objective in discovering unknown confounders beyond those reported in existing observational studies. We also note the inconsistency in the reported covariates, indicating opportunities to standardize confounder adjustment. Often, researchers only condition on variables for which they have data. Suppose there are known confounders for which data has not been collected. In that case, the opportunity is often missed to inform future researchers of these variables is often missed since reporting on unmeasured confounders is not standard practice in AD research. However, reporting on unmeasured confounders is often a helpful value-added proposition if these variables are included visually in a DAG since such inclusion alerts future researchers of the need to consider these variables in their models.

In our comparative evaluation, comparing outputs between the KG and SemMedDB_Complete_, we found a relatively high concordance regarding the variables identified and the causal roles played by those variables. We found contradictions, and a significant portion of the KG and SemMedDB_Complete_ search results did not co-occur. A partial explanation is that the KG and the structured literature-derived knowledge derive from completely different processes, origins, and purposes. The KG uses information derived from granular, ontology-grounded knowledge, while SemMedDB relies on the output of advanced NLP systems. Some concepts appeared in the KG that did not appear in SemMedDB_Complete_, e.g., inflammatory response (although a literature search produces promising evidence supporting inflammatory response as a confounder^146–150^), and vice versa, e.g., leukoencephalopathy. SemMedDB_Complete_ had a better recall of lifestyle-related confounders. The KG excelled at biological plausibility. Arguably, the KG and SemMedDB_Complete_ complement one another in their coverage. SemMedDB_Complete_ will have a different set of assertions than what MachineReadingDB_Scoped_ has or that make sense through the distillation process searching the combined KG. These different sets of assertions can contradict one another, resulting in topologies for the variables/nodes that reflect those contradictory assertions. Of this information, it is not known what is erroneous or not until these components undergo verification by human experts.

Human experts suggested reasonable confounders not included in the KG search results. For example, two external correspondents with backgrounds in neurology noted two important confounders missing from the list of confounders identified by the KG and the literature: pesticides and adverse childhood experiences. The scoping of the literature also likely impacted the results. For example, the pesticide was present when searching SemMedDB_Complete_. In addition, childhood trauma has been noted as a potential risk factor for depression and AD^151^. However, “adverse childhood event” was missing from SemMedDB_Complete_, further suggesting the importance of a human-in-the-loop process for identifying potential confounders.

To the best of our knowledge, the literature has identified covariates that act as either confounder/mediators or confounder/colliders, but not all three roles, i.e., “the chimera.” We verified the plausibility of stroke as an example of a chimera for depression and AD. Since chimeras are a new category of variable, no work to our present knowledge has been performed to identify chimeras explicitly.

### 4.1 Related Work

There is vast, and still growing, literature on integrating and using forms of computable biomedical knowledge. For example, related work has been done in defining ontologies for representing knowledge for neurodegenerative diseases^78^, identifying erroneous information by applying transitive closure over causal predicates^78^, modeling argument structure^152^, detecting contradictions in statements extracted from the literature^153, 154^, drug repurposing pipelines^32, 33, 35, 155, 156^, and elucidating mechanisms in systems and molecular biology^40^. Simply put, the vast body of tangentially related work would be impossible to condense. However, in the interest of space, since that literature is so vast, we will confine the discussion to a review of related work regardless of application area on the problem of pruning spurious nodes resulting from erroneous information.

The uncritical use of computable knowledge can result in a causal graph with low precision^35^. Thus, the development of methods for filtering out erroneous information is arguably a major hindrance to automating causal feature selection and has spurred much creativity in developing novel approaches to address this problem. For example, Hu et al. used information from authoritative ontologies to orient the edges of a graphical causal model studying risk factors for AD^36^. In the approach of Nordon et al., edges were removed if the variables were not correlated in their corpus of electronic medical record data^35^. The incompleteness of knowledge was another major problem, and approaches exist that attempt to solve both problems by using concept similarity measures to filter out erroneous edges, while inferring new edges. Zhang et al. report using neural graph embedding method PubMedBioBert^33^. Malec et al. used cutoff scores for string-based and vector-based confounder search^63, 157^.

Our novel causal feature selection method builds on and differs from previous research in the following regards. In contrast to the previous work, our method combines the breadth of literature-derived information with the precision of ontology-grounded resources. In addition, we introduce refinements to confounder search that allows us to identify a comprehensive set of variables with which to adjust without inducing bias (by identifying and filtering out potential colliders and mediators).

The present work builds on the ideas from literature-based discovery, an area pioneered by Don Swanson and Neil R. Smallheiser^158–161^. Originating in the field of literature-based discovery, an essential notion of our approach relies on the idea of a discovery pattern, or the use of semantic constraints to yield potentially interesting hypotheses^158, 159^. The first discovery patterns were manually constructed patterns noticed by observing recurring relationships between concepts^62, 162^. More recently, new applications for discovery patterns have been identified, and methods have been developed to infer discovery patterns automatically^163, 164^. Discovery patterns can be and have been implemented in any query or knowledge representation language that considers how concepts relate to one another, including SPARQL^165^, SQL^166^, and advanced methods for querying distributed semantic representations^63, 157, 167, 168^ of structured knowledge.

Related to the present work, in its explanatory aspect, would be a paper by Elsworth et al. using literature-derived triples to identify intermediate concepts^169^. However, the objective of their paper was focused on identifying mediating mechanisms from a theoretical perspective rather than the context of identifying contextual knowledge relevant to data-driven studies. Other notable work related to an explanation would be the EpiphaNet project, an interactive knowledge discovery system developed by Cohen et al.^170^ EpiphaNet allows scientists to navigate relationships that interest them visually and interactively.

Other prior work has applied structure learning methods to observational data to recapitulate known and unknown edges between the biomedical entities under the models^171^, and provide automated cogent explanatory paths relating to drugs and adverse events^29^.

### 4.2 Strengths and limitations

A significant strength of the present work was utilizing ontology-grounded knowledge resources in the KG. The ontology-grounded KG helped prune erroneous information and provide mechanistic explanations in the form of reasoning paths that describe how the variables identified were related to depression and AD.

A methodological limitation may have arisen in our search procedure. In the interest of search efficiency, we constrained the max length of our search. The KG search also identified genes, enzymes, and drug exposures as potential confounders. However, the KG search only produced output for phenotypes playing collider and mediator roles. Hence, for illustration purposes, our analysis focused primarily on confounders that were conditions or phenotypes. We suspect that the threshold applied to limit search depth may have constrained the results, resulting in lower recall than a deeper search in these categories. Several variables in these categories were recapitulated in our literature review in *Table 5*, e.g., homocysteine, and APOEɛ4 carrier status.

The present study successfully incorporated output from different machine reading systems with distinct but complementary purposes, beginning with their respective focus domains. For example, SemRep excels at extracting pharmacogenomic^172^ and clinical knowledge, whereas INDRA/REACH excels with detailed mechanistic knowledge of proteins, thus facilitating the linkage between detailed mechanistic and clinical knowledge. However, we could not capitalize on this knowledge because of imperfect terminology mapping. This form of information loss arises when a concept appears in the source terminology that is missing in the target. As seen in the PRISMA diagram in *Figure 4*, many concepts identified in the literature were lost in translation undergoing the terminology mapping procedure. There was also information loss because of a lack of granularity in the mappings we were able to find. For example, all of the post-translational modifications were mapped to the predicate **`biomolecularly_interacts_with`**, which is not unidirectional in its meaning: molecularly interacts with is associational. As a result, we may have lost information about directionality in some of the predicates. In other cases, inferences may have been erroneous.

Erroneous inferences may arise from errors inherited from the machine reading systems or the ontology-grounded knowledge in the KG. This is because our causal feature selection approach relies on KG search and KG search depends on the information from the machine reading systems and the ontologies. However, note that the errors inherent in the machine reading systems are distinct from the errors in downstream applications, i.e., causal modeling. For more specifics on machine reading system-specific errors, Ahlers et al. discusses pharmacogenomic predications^172^, while Rindflesch discusses difficulties extracting predications concerning the etiology of disease^127^. For approaches for post-processing machine reading outputs that address erroneous outputs and other aspects of knowledge hygiene, see Holtsapple et al. ^33^ for a post-processing workflow for INDRA and Zhang et al.^33^ for a post-processing workflow for SemRep predications.

It is important to acknowledge the limitations of the mapping process and the need for further optimization to improve its performance in future studies. The mapping of INDRA concepts to UMLS using MetaMap resulted in a concept loss. Many unmapped concepts were non-biological terms or abbreviations that could not be disambiguated without the source texts. Reduced accuracy in the presence of ambiguity is a known limitation of MetaMap^68^.

We also speculate that the poor performance of this mapping process may be partially explained by a mismatch between the general-purpose biomedical free-text use case for MetaMap^68, 173^ and the unique sublanguage required for post-translational modifications (PTMs) of proteins. Drawing on the work of Zellig Harris^174^, Friedman defined a sublanguage as a language that is used in a specific domain or field of expertise, which may have its own unique vocabulary and syntax^175^. Similarly, Rindflesch and Fiszman discussed the importance of considering sublanguage when developing NLP tools for biomedical texts^176^. Hence, MetaMap was not as successful at recognizing and resolving semantically ambiguous concepts in the PTM domain as the INDRA/REACH NLP system, which was specifically designed to recognize concepts (and predicates) in the PTM domain.

In addition, the current implementation of PheKnowLator in mapping from UMLS to OBO ontologies (see *Listing 1*) may have also resulted in a loss of mappings, leading to less complete explanations and collider, confounder, and mediators in KG search results. This is because PheKnowLator only includes patterns that have OBO mappings in its path searches. While acceptable as a proof of concept, where the focus is on feasibility, this limitation highlights the need for further work to improve and expand the mappings and evaluate the performance characteristics of the mapping processes. A detailed list of triples and mappings is available for the interested reader for further examination. The semantic types of mapped and unmapped terms can be analyzed by comparing the mapping tables (see **File VII. *Appendix_C.docx***) and querying the UMLS MRSTY tables (available at this link: https://www.nlm.nih.gov/research/umls/licensedcontent/umlsknowledgesources.html).

Errors from the machine reading NLP can take different forms and result in causal models that are either incomplete, inaccurate or both, resulting from differential confounder misclassification^177^. Either inaccuracies or incompleteness of information can lead to bias from omitting an important confounder or adjusting for a combined role variable. For example, residual bias from not conditioning on an important confounder or omitted variable bias^20^, and differential conclusions. Similarly, a missing collider relationship can result in collider-stratification bias if the variable is adjusted for. In simulation studies, this bias can range from 10-15% (or more) in either direction^178^. On the other hand, if a variable is adjusted for as a confounder when its other role as a mediator relationship is not identified, overadjustment bias^179^ or treatment-confounder feedback bias may result. In this scenario, the measure of effect or association tends to be biased toward the null, since the mediator is an aspect of the causal effect of the exposure. Fortunately, appropriate estimation methods designed for these settings, such as G-methods^180, 181^ and IPTW^182–185^ can be utilized to address these challenges.

Finally, bias of unpredictable direction can arise from a combination of incompleteness and inaccuracies in machine reading and the ontologies, depending on the nature and strength of the variables. However, residual bias may remain from not adjusting for these problematic variables.

Science is not infallible, and establishing ground truth is a contentious process. Erroneous conclusions may be drawn from undiagnosed bias from confounding bias^7^, selection bias, measurement error bias^186^, missing data^21, 187^, or a combination of some or all of the above. For example, selection bias can result from restrictions on the study population, such as inclusion/exclusion criteria based on healthcare utilization (a collider),^188–190^ or on a prognostic test (even if the prognostic does not affect the outcome, and is not a collider).^191^

Another limitation that inheres in both forms of structured knowledge we used in this study. It is also essential to consider not just the accuracy and completeness of the structured knowledge for causal feature selection but also the transportability of the causal relationships identified when interpreting the results of a study^192–194^. In other words, it is also important to consider cases when the structured knowledge is accurate, but the causal relationship identified by the study is not transportable to the target population. For example, the relationship between a risk factor and a disease may be accurately identified in a study population but may not hold true in a population with different demographics, such as age, gender, or ethnic background^195, 196^. Similarly, the accuracy of a causal relationship may be impacted by differences in how the variables are measured and categorized. For example, a relationship between a risk factor and a disease may be identified in a study using self-reported data but may not hold true when the same relationship is assessed using objective measures. Furthermore, the non-transportability of causal relationships may also arise from differences in the underlying causal structure of the populations, giving rise to heterogeneous treatment effects^197^. For example, a causal relationship identified in a study population may result from a common cause that is not present in the target population, such as shared environmental exposure, or vice versa. Finally, factors such as the distribution of confounding variables, measurement and categorization of variables, and underlying causal structure should be carefully considered when extrapolating the findings to other populations.

Another limitation was the small size of the literature corpus and the small size likely had inadvertent negative side effects on the recall of variables and roles. Also, since we scoped the inputs from the literature-derived computable knowledge to focus exclusively on AD (the outcome) and the identification of confounders and other roles requires comprehensive knowledge of both the exposure (depression) and outcome mechanisms, we likely missed titles having to do with our exposure, depression, limiting the ability to identify confounders. A larger and more representative sample of the knowledge in the literature could partially address the missing bits of knowledge.

There is always the risk of latent and unmeasured confounding regardless of the causal feature selection method. Knowledge, like all things human, is never complete or perfect. Both in spite of and because of these caveats, computational tools for causal feature selection are critical for simplifying the task of exploring the causal structure underlying scientific questions and for identifying more comprehensive, robust, and plausibly valid feature sets than is traditionally possible.

### 4.3 Future directions

*Automated visualization of KG search output.* In the present work, we built the diagrams manually. In future work, we will develop tools that automatically compile and generate causal graphs and reasoning path diagrams after collecting requirements from human content experts. Explanatory power and ease of use are critical for engaging with and obtaining a clearer view of the status of human content-matter experts. Additional work is also needed regarding how to best gather input from human content-matter experts concerning variables identified via our workflow to derive a reference standard for a subset of modifiable risk factors. An automated explanation capability would significantly facilitate the investigation of causal feature selection methods to facilitate review.

The manual verification exercises demonstrated how KG and structured literature search could be fruitfully combined with manual review searching PubMed to adjudicate KG search output.

However, human experts would likely require more than just source sentences to classify variables by the causal roles of identified variables. To optimize the transportability of causal findings from the literature and ontology-grounded information to inform epidemiological studies, more work needs to be done to collect meta-data for causal assertions, including information about study characteristics, whether derived from observational studies, RCTs, or GWAS assays. The Cochrane Handbook for Systematic Reviews of Interventions provides recommendations on data collection of characteristics of observational studies to assess the transportability of their findings^198^. In this vain, recent work^199^ highlighted a workflow for assessing the quality of confounding control in observational studies included in meta-analyses. Efforts collecting a field-specific database of confounders are foundational for evaluating outputs from structured knowledge. However, it will also be important to gather information on mediators and colliders. Mediators are particularly important if they are also confounders, as such variables may be CAPTs^17^, mentioned earlier. However, few papers report these types of variables, unless the study is focusing on estimating effects in the context of time-varying confounders.

The estimation of multiple concurrent exposures is a slowly emerging topic^200–202^. In this paper, we focused on understanding the complexities of a single exposure, depression. We did not explore the causal feature selection for scenarios with multiple concurrent exposures, While there is some existing research on joint treatment effects, the emphasis has been on estimation procedures rather than causal feature selection. However, we have begun to consider the implications of considering complex exposure regimes in preliminary concurrent work.

We briefly describe a knowledge-based procedure for handling the technicalities of multiple concurrent exposures, a slowly emerging topic relevant to public health aspects of AD research. Returning to our original use case, consider depression and an anti-depressant treatment for depression to be the joint/multiple (concurrent) exposures with outcome AD. To estimate the effect of these exposures on AD, we first identify and adjust for confounders that affect each exposure and the outcome. Confounders for each exposure-outcome relationship should be selected separately by identifying variables that affect the exposure and outcome. For depression and AD, potential confounders could include age, sex, education, and genetic risk factors for AD. For anti-depressant use and AD, confounders could include the severity of depression, comorbidities, and prior medication use^203, 204^. We then combine both of these confounders into a single graph. Note that certain confounders will be unique to one exposure, and other variables will be common to both.

A popular way to handle confounders that are unique to each exposure when estimating causal effects is through marginal structural models (MSMs)^184, 205, 206^. Standard adjustment methods, including multivariate logistic regression, fail in such scenarios. MSMs allow for the estimation of the total effect of joint treatment while controlling for the confounding effects of each exposure. Time-varying confounders that may be uniquely associated with depression and AD could include changes in cognitive function^51, 207–209^ over time. Cognitive function could also be a time-varying confounder as it may both affect and be influenced by depression and, in turn, may affect treatment decisions and outcomes. In contrast, time-varying confounders that may be uniquely associated with anti-depressant use and AD could include changes in the severity of depression and medication adherence over time. Certain other confounders could be common to both exposures, e.g., sociodemographic characteristics like age, race, and gender. Both unique and common confounders should be included in the analysis to control for their effects. Unique confounders for each exposure can be handled by including them in the weighting function, which balances the distribution of the confounders across treatment groups. Time-fixed confounders, e.g., APOEε2/3/4 status^210–212^) can be included as covariates, while time-varying confounders can be handled using IPTW^15, 184, 213, 214^ or G-methods^15, 16, 180, 181, 215^. Readers interested in a more detailed discussion of diagnostic approaches for addressing joint exposures and time-varying confounding in research methods may refer to Jackson’s work^15^. By considering multiple exposures, we can capture a more comprehensive picture of the complex interplay between various risk factors and treatment outcomes, ultimately leading to more precise and effective interventions tailored to individual subgroups.

Returning to the public health question, more reliable estimation of joint treatment effects has the potential to significantly enhance our ability to identify heterogeneous treatment effects within subgroups and optimize interventions for their effect by subgroup on modifiable risk factors, and explored in greater detail by Barnes et al. ^216^. Finally, in parallel work, we are working on a method addressing the problem of “M”-bias. “M”-bias is a type of collider bias, wherein the bias is amplified by inducing a spurious dependency between the exposure and the outcome conditioning on a collider^217, 218^. In “M”-bias, there are confounders that are themselves effects of two or more other confounders. To reduce bias in this scenario, causal relationships between the confounders must be considered.

### 4.4 Practical Implications

The variables in the “Confounders only” category in *Figure 11* could be said to be a potentially strong starting point for adjustment sets. These concepts fulfill the definition of a confounder as a mutual cause of the exposure (depression) and the outcome (AD) of interest without being effects of either depression or AD and have been vetted by two distinct causal feature selection methods but require review by human content-matter experts.

Yet, all hope is not lost for confounders that play multiple roles. Here, we briefly consider how to handle confounders that also might play other roles, e.g., colliders or mediators. When a variable is both a confounder and a collider, adjusting for it based on data measured before the exposure may still result in collider-stratification bias^23, 219^. This occurs because adjusting for a collider affects the relationship between the exposure and the outcome and may introduce bias into the estimated effect. A common, though incorrect, approach to prevent collider bias is to include only covariates measured before the exposure. However, this approach may still not fully address the issue of collider-stratification bias^220^.

Instead, a more thorough and careful approach may be necessary, such as using causal inference techniques, such as propensity score matching methods, e.g., IPTW^182–185^, to adjust for the selection bias induced by the collider, or adjusting for other variables that can block the collider pathway. In the latter case, knowledge of the causal structure and measure of the other variable is required. In addition, sensitivity analyses may be conducted to assess the results’ robustness to the adjustment method. Nevertheless, the decision on whether to adjust for a variable that is both a confounder and a collider should be based on a thorough understanding of the underlying causal mechanisms and relationships between the variables and a critical evaluation of the potential sources of bias and confounding. Regardless of the estimation method, sensitivity analyses and diagnostics are recommended for the best results. Illuminating further discussion on sensitivity analysis when collider-stratification bias is suspected is discussed in a paper by Kennedy and Balakrishnan^7^.

Recent work has advocated for the development of processes for the systematic review of potential confounders by content-matter experts with the objective of the creation of field-specific databases of confounders^222^ and routinely reporting causal DAGs containing all known confounders in observational research^222^. Tennant et al. suggest reporting variables measured in observational studies and variables that researchers would have liked to have measured but for which they had not collected data in their current study^223^. They also suggest using a standardized format to report causal DAGs such as DAGitty^224^. Such reporting standards, employing FAIR (fair, accessible, interoperable, and reusable) data principles^225^, ensure shareability and, if adopted, could make observational studies more rigorous and reproducible and allow researchers to build more concretely on each other’s work, making advances with each iteration to answer difficult causal questions from large sets of non-randomized patient health data “across sites and studies” as suggested by Blum et al.^222^. Iteratively building on the knowledge of confounders could spur advance in AD studies, perform informed sensitivity analyses, and quantify the extent of “unknown unknowns” using the e-value metric^222, 226, 227^. The e-value metric essentially quantifies the prevalence the confounder would need to have to explain away an association, as per the classic study of Cornfield et al. on smoking and lung cancer^228–231^.

Observational studies that employ a more comprehensive, refined adjustment set and standardization of adjustment sets by expert review could improve the rigor of observational studies (by standardizing adjustment, a point raised above) and bolster the precision of causal effect estimates. This information could then be applied to facilitate causal inference at scale over large centralized and federated data stores, such as PCORnet^232, 233^ and *All_of_Us*^234^.

Lastly, the following bullet points represent the contributions of this work: (1.) contributes information on potential confounders to consider when designing rigorous observational studies of depression and AD; (2.) provides explanatory reasoning paths providing mechanistic hypotheses for the causal edges identified; and (3.) shows how semantic and causal inference can be partially combined.

## 5. Conclusion

In summary, our study has demonstrated the potential of utilizing knowledge graphs (KG) to identify potential confounding, colliding, and mediating variables in the relationship between depression as an exposure and AD as an outcome. Our methodology has partially addressed the limitations of literature-derived structured knowledge by distilling it through ontology-grounded knowledge in a KG. Our findings have revealed that KGs offer an efficient way to represent and automate reasoning over knowledge, leading to the generation of mechanistic hypotheses concerning biomolecular processes underlying confounders. Moreover, our research has highlighted that semantic inference can enhance causal inference, though it requires further investigation.

Ultimately, our work aims to answer epistemological questions. While depression and AD serve as a prime example in our study, it is not the sole reason for our research. More research needs to be conducted on our analytical strategy to reach a definitive conclusion for this specific case. We anticipate that this work and future studies will progress the rigor of observational studies.

The complexity of variables underscores the value of a combination of machine-human strategies, where machines can proficiently search for large-scale knowledge, linking clinical phenotypes with molecular processes and disease definitions. Human experts can focus on comprehending the appropriateness of identified variables in their context, i.e., transportability. Computational tools that simplify access to knowledge can help scientists conduct more meticulous observational studies and prioritize public health interventions with greater confidence. Future work is needed to enhance KG search for causal feature selection, and our methods can be effortlessly extended to other content areas.

## Supporting information

File I. Appendix_A.docx. Background S1. Logical properties.

File_II-AppendixB-Details

File III. AdjustmentSetDepression2AD.xlsx. Worksheet S1. Conditions. Worksheet S2. Substances. Worksheet S3. GenesEnzymes.

File IV. predicatePropertyAnalysis.xlsx. This file contains our analysis of the logical properties of symmetry and transitivity for the predicates

File V. ConfounderSourceSentences.xlsx. This file contains source sentences for three potential confounders that have not been reported as covariates

File VI. ChimeraSourceSentences.xlsx. This file contains source sentences for stroke as a chimeric variable for depression and AD.

File VII. Appendix_C.docx. This file contains documentation for access the Zenodo archive containing Jupyter notebooks for knowledge hygiene operation

File VIII. MDD2ADandDementia-v1.0.xlsx. This file contains a pilot metareview of confounders collected from reported confounders in published studies.

## Supplementary Information

Supplementary Information accompanies this paper as described below.

**Additional files: File I. *Appendix_A.docx***. **Background S1.** Logical properties. **File II. *Appendix_B.docx***. **Listing S1.** PubMed scoping query. **Table S1.** Mappings for UMLS CUIs to RO. **Table S2.** UMLS CUI definitions for depression and AD. **Table S3.** Complete analysis of KG search with SemMedDBs. **Table S4.** Comparison of confounders - search results belonging to the gene or enzyme, protein, or semantic categories. **Table S5.** Comparison of confounders - search results belonging to the drug or hormone semantic categories. **Table S6.** Comparison of phenotype/condition confounders playing simple roles. **Table S7.** Comparison of phenotype/condition confounders playing combined roles. **Figure S1.**Venn diagram of complete results for all semantic types for simple roles. **Figure S2.** Venn diagram of complete results for all semantic types for combined roles. **File III. *AdjustmentSetDepression2AD.xlsx***. **Worksheet S1.** Conditions. **Worksheet S2.** Substances. **Worksheet S3.** GenesEnzymes. **File IV. *predicatePropertyAnalysis.xlsx***. This file contains our analysis of the logical properties of symmetry and transitivity for the predicates from the machine reading systems. **File V. *ConfounderSourceSentences.xlsx***. This file contains source sentences for three potential confounders that have not been reported as covariates in published observational studies looking at depression and AD. **File VI. *ChimeraSourceSentences.xlsx***. This file contains source sentences for stroke as a chimeric variable for depression and AD. **File VII. *Appendix_C.docx***. This file contains documentation for access the Zenodo archive containing Jupyter notebooks for knowledge hygiene operations (terminology mapping and logical closure), the SPARQL queries and raw output from running the queries, and the SemMedDB queries. **File VIII. *MDD2ADandDementia-v1.0.xlsx***. This file contains a pilot metareview of confounders collected from reported confounders in published studies.

## Acknowledgments

The authors would like to thank the anonymous reviewers for helpful suggestions and challenging comments that improved the paper. We would also like to thank health sciences librarian Helena M. VonVille for assisting in designing the PubMed query to scope the literature. Previous versions of this paper benefited from the suggestions and comments of Professors Gregory F. Cooper, Andrew Stern, and Howard Aizenstein. We would also like to recognize Elizabeth Wu of Alzforum (http://www.alzforum.org) for discussion regarding previous versions of this paper. Thanks to Sarah Wenyon for proofreading. SAM takes responsibility for all errors.

## Availability of data and materials

The data supporting these findings are available from this project’s Zenodo repository (https://doi.org/10.5281/zenodo.6785307). No restrictions apply to the public availability of these data.

## Authors’ Contributions

SAM had full access to all of the data in this study and took full responsibility for the integrity and accuracy of the data analysis. RDB supervised the study. SBT and RDB conceived of the graph querying methods with SAM’s input. RDB and SAM conceived and executed the initial study design and SAM analyzed the study data, interpreted study findings, and drafted the manuscript along with RDB. TJC provided invaluable input in constructing the KG. SMA, CES, HTK, ASL, TJC, PWM, and RDB critically reviewed drafts of the manuscript. The corresponding author attests that all listed authors meet the authorship criteria and that no other meeting the criteria have been omitted. All authors read and approved the final manuscript.

## References

1 Cartwright N. Are RCTs the Gold Standard? BioSocieties [Internet]. 2007 Mar [cited 2017 Jul 21];2(1):11–20. Available from: http://www.palgrave-journals.com/doifinder/10.1017/S1745855207005029

2 VanderWeele TJ, Shpitser I. On the definition of a confounder. Ann Stat. 2013 Feb;41(1):196–220.

3 VanderWeele TJ. Principles of confounder selection. Eur J Epidemiol [Internet]. 2019 [cited 2019 Aug 20];34(3):211–9. Available from: https://www.ncbi.nlm.nih.gov/pmc/articles/PMC6447501/

4 Arntzenius F. Reichenbach’s Common Cause Principle. In: Zalta EN, editor. The Stanford Encyclopedia of Philosophy [Internet]. Fall 2010. Metaphysics Research Lab, Stanford University; 2010 [cited 2019 Dec 10]. p. 1. Available from: https://plato.stanford.edu/archives/fall2010/entries/physics-Rpcc/

5 VanderWeele TJ, Shpitser I. A new criterion for confounder selection. Biometrics [Internet]. 2011 Dec;67(4):1406–13. Available from: https://www.ncbi.nlm.nih.gov/pubmed/21627630

6 Groenwold RHH, Sterne JAC, Lawlor DA, Moons KGM, Hoes AW, Tilling K. Sensitivity analysis for the effects of multiple unmeasured confounders. Ann Epidemiol [Internet]. 2016 Sep 1 [cited 2020 Sep 20];26(9):605–11. Available from: http://www.sciencedirect.com/science/article/pii/S1047279716302253

7 Pearl J. Causality: Models, Reasoning, and Inference [Internet]. 2nd ed. Cambridge: Cambridge University Press; 2009 [cited 2017 Jul 21]. Available from: http://ebooks.cambridge.org/ref/id/CBO9780511803161

8 Patorno E, Glynn RJ, Hernandez-Diaz S, Liu J, Schneeweiss S. Studies with many covariates and few outcomes: selecting covariates and implementing propensity-score-based confounding adjustments. Epidemiol Camb Mass. 2014 Mar;25(2):268–78.

9 Talbot D, Diop A, Lavigne-Robichaud M, Brisson C. The change in estimate method for selecting confounders: A simulation study. Stat Methods Med Res. 2021 Sep;30(9):2032–44.

10 Variable Selection for Confounding Adjustment in High-dimensional Covariate Spaces When Analyzing Healthcare Databases. Vol. 28. United States; 2017.

11 Shpitser I, VanderWeele T, Robins JM. On the Validity of Covariate Adjustment for Estimating Causal Effects. :10.

12 MEDLINE PubMed Production Statistics [Internet]. U.S. National Library of Medicine; [cited 2022 Jul 3]. Available from: https://www.nlm.nih.gov/bsd/medline_pubmed_production_stats.html

13 Hernan MA, Robins JM. Causal Inference [Internet]. Taylor & Francis; 2017. (Chapman & Hall/CRC Monographs on Statistics & Applied Probab). Available from: https://books.google.com/books?id=_KnHIAAACAAJ

14 Karim ME, Tremlett H, Zhu F, Petkau J, Kingwell E. Dealing with Treatment-confounder Feedback and Sparse Follow-up in Longitudinal studies - Application of a Marginal Structural Model in a Multiple Sclerosis Cohort. Am J Epidemiol. 2020 Oct 30;

15 Jackson JW. Diagnostics for confounding of time-varying and other joint exposures. Epidemiol Camb Mass [Internet]. 2016 Nov [cited 2020 May 10];27(6):859–69. Available from: https://www.ncbi.nlm.nih.gov/pmc/articles/PMC5308856/

16 Robins J, Hernan M. Estimation of the causal effects of time-varying exposure. In: Longitudinal Data Analysis. 2008. p. 553–99.

17 Pega F, Blakely T, Glymour MM, Carter KN, Kawachi I. Using Marginal Structural Modeling to Estimate the Cumulative Impact of an Unconditional Tax Credit on Self-Rated Health. Am J Epidemiol. 2016 Feb 15;183(4):315–24.

18 Sackett DL. Bias in analytic research. J Chronic Dis. 1979;32(1–2):51–63.

19 Elwert F, Winship C. Endogenous Selection Bias: The Problem of Conditioning on a Collider Variable. Annu Rev Sociol [Internet]. 2014 Jul;40:31–53. Available from: https://www.ncbi.nlm.nih.gov/pubmed/30111904

20 Steiner PM, Kim Y. The Mechanics of Omitted Variable Bias: Bias Amplification and Cancellation of Offsetting Biases. J Causal Inference [Internet]. 2016 Sep;4(2):20160009. Available from: https://www.ncbi.nlm.nih.gov/pubmed/30123732

21 Abrahamowicz M, Bjerre LM, Beauchamp ME, LeLorier J, Burne R. The missing cause approach to unmeasured confounding in pharmacoepidemiology. Stat Med. 2016 Mar 30;35(7):1001–16.

22 Grätz M. When Less Conditioning Provides Better Estimates: Overcontrol and Collider Bias in Research on Intergenerational Mobility [Internet]. Working Paper Series. Stockholm University, Swedish Institute for Social Research; 2019 Jun [cited 2020 Jun 16]. (Working Paper Series). Report No.: 2/2019. Available from: https://ideas.repec.org/p/hhs/sofiwp/2019_002.html

23 Cole SR, Platt RW, Schisterman EF, Chu H, Westreich D, Richardson D, et al. Illustrating bias due to conditioning on a collider. Int J Epidemiol [Internet]. 2010 Apr;39(2):417–20. Available from: https://www.ncbi.nlm.nih.gov/pubmed/19926667

24 Banack HR, Kaufman JS. From bad to worse: collider stratification amplifies confounding bias in the “obesity paradox”. Eur J Epidemiol. 2015 Oct;30(10):1111–4.

25 Funk C, Baumgartner W, Garcia B, Roeder C, Bada M, Cohen KB, et al. Large-scale biomedical concept recognition: an evaluation of current automatic annotators and their parameters. BMC Bioinformatics [Internet]. 2014 Feb 26 [cited 2023 Feb 25];15(1):59. Available from: https://doi.org/10.1186/1471-2105-15-59

26 Mower J, Subramanian D, Shang N, Cohen T. Classification-by-Analogy: Using Vector Representations of Implicit Relationships to Identify Plausibly Causal Drug/Side-effect Relationships. AMIA Annu Symp Proc AMIA Symp. 2016;2016:1940–9.

27 Mower J, Cohen T, Subramanian D. Complementing Observational Signal with Literature-derived Distributed Representations for Post-marketing Drug Surveillance. Drug Saf [Internet]. 2020 Jan [cited 2020 Sep 17];43(1):67–77. Available from: https://www.ncbi.nlm.nih.gov/pmc/articles/PMC7243821/

28 Mower J, Subramanian D, Cohen T. Learning predictive models of drug side-effect relationships from distributed representations of literature-derived semantic predications. J Am Med Inform Assoc [Internet]. 2018 Jul 11;ocy077–ocy077. Available from: http://dx.doi.org/10.1093/jamia/ocy077

29 Shang N, Xu H, Rindflesch TC, Cohen T. Identifying plausible adverse drug reactions using knowledge extracted from the literature. J Biomed Inform. 2014 Dec;52:293–310.

30 Fathiamini S, Johnson AM, Zeng J, Araya A, Holla V, Bailey AM, et al. Automated identification of molecular effects of drugs (AIMED). J Am Med Inform Assoc JAMIA. 2016 Jul;23(4):758–65.

31 Cohen T, Widdows D, De Vine L, Schvaneveldt R, Rindflesch TC. Many Paths Lead to Discovery: Analogical Retrieval of Cancer Therapies. In: Busemeyer JR, Dubois F, Lambert-Mogiliansky A, Melucci M, editors. Quantum Interaction. Berlin, Heidelberg: Springer; 2012. p. 90–101. (Lecture Notes in Computer Science).

32 Nordon G, Koren G, Shalev V, Horvitz E, Radinsky K. Separating Wheat from Chaff: Joining Biomedical Knowledge and Patient Data for Repurposing Medications. Proc AAAI Conf Artif Intell [Internet]. 2019 Jul 17 [cited 2020 Mar 5];33(01):9565–72. Available from: https://aaai.org/ojs/index.php/AAAI/article/view/5017

33 Zhang R, Hristovski D, Schutte D, Kastrin A, Fiszman M, Kilicoglu H. Drug repurposing for COVID-19 via knowledge graph completion. J Biomed Inform. 2021 Mar;115:103696.

34 Müller HM, Kenny EE, Sternberg PW. Textpresso: An Ontology-Based Information Retrieval and Extraction System for Biological Literature. PLoS Biol [Internet]. 2004 Nov [cited 2022 Jul 4];2(11):e309. Available from: https://www.ncbi.nlm.nih.gov/pmc/articles/PMC517822/

35 Nordon G, Koren G, Shalev V, Kimelfeld B, Shalit U, Radinsky K. Building Causal Graphs from Medical Literature and Electronic Medical Records. Proc AAAI Conf Artif Intell [Internet]. 2019 Jul 17 [cited 2022 Jun 1];33(01):1102–9. Available from: https://ojs.aaai.org/index.php/AAAI/article/view/3902

36 Hu H, Kerschberg L. Improved Causal Models of Alzheimer’s Disease. In: 2021 IEEE 45th Annual Computers, Software, and Applications Conference (COMPSAC). 2021. p. 274–83.

37 Besnard P, Cordier M o, Moinard Y. Inferring Causal Explanations. Hunt Parsons Eds. 1999;99:55–67.

38 Friedman S, Magnusson I, Sarathy V, Schmer-Galunder S. From Unstructured Text to Causal Knowledge Graphs: A Transformer-Based Approach. ArXiv220211768 Cs [Internet]. 2022 Feb 23 [cited 2022 Feb 25]; Available from: http://arxiv.org/abs/2202.11768

39 Thomas PD, Hill DP, Mi H, Osumi-Sutherland D, Van Auken K, Carbon S, et al. Gene Ontology Causal Activity Modeling (GO-CAM) moves beyond GO annotations to structured descriptions of biological functions and systems. Nat Genet [Internet]. 2019 Oct [cited 2021 Aug 15];51(10):1429–33. Available from: https://www.nature.com/articles/s41588-019-0500-1

40 Tripodi IJ, Callahan TJ, Westfall JT, Meitzer NS, Dowell RD, Hunter LE. Applying knowledge-driven mechanistic inference to toxicogenomics. Toxicol Vitro Int J Publ Assoc BIBRA. 2020 Aug;66:104877.

41 Sarvet AL, Stensrud MJ. Without Commitment to an Ontology, There Could Be No Causal Inference. Epidemiology [Internet]. 2022 May [cited 2022 Jun 5];33(3):372–8. Available from: https://journals.lww.com/epidem/Citation/2022/05000/Without_Commitment_to_an_Ontology,_There_Could_Be.9.aspx

42 Liu K, Hogan WR, Crowley RS. Natural Language Processing methods and systems for biomedical ontology learning. J Biomed Inform. 2011 Feb;44(1):163–79.

43 Kejriwal M. Domain-Specific Knowledge Graph Construction [Internet]. Springer International Publishing; 2019. Available from: https://books.google.com/books?id=naHrvwEACAAJ

44 Vos T, Lim SS, Abbafati C, Abbas KM, Abbasi M, Abbasifard M, et al. Global burden of 369 diseases and injuries in 204 countries and territories, 1990–2019: a systematic analysis for the Global Burden of Disease Study 2019. The Lancet [Internet]. 2020 Oct 17 [cited 2021 Sep 29];396(10258):1204–22. Available from: https://www.sciencedirect.com/science/article/pii/S0140673620309259

45 Komarova NL, Thalhauser CJ. High Degree of Heterogeneity in Alzheimer’s Disease Progression Patterns. PLOS Comput Biol [Internet]. 2011 Nov 3 [cited 2019 Oct 2];7(11):e1002251. Available from: https://journals.plos.org/ploscompbiol/article?id=10.1371/journal.pcbi.1002251

46 Association AP. DSM 5 Diagnostic and statistical manual of mental disorders. DSM 5 Diagn Stat Man Ment Disord [Internet]. 2013 [cited 2022 Mar 13];947 p.-947 p. Available from: https://pesquisa.bvsalud.org/portal/resource/pt/psa-52826

47 Bains N, Abdijadid S. Major Depressive Disorder. In: StatPearls [Internet]. Treasure Island (FL): StatPearls Publishing; 2022 [cited 2022 Jun 5]. Available from: http://www.ncbi.nlm.nih.gov/books/NBK559078/

48 Cai N, Choi KW, Fried EI. Reviewing the genetics of heterogeneity in depression: operationalizations, manifestations and etiologies. Hum Mol Genet [Internet]. 2020 Sep 30 [cited 2021 Sep 28];29(R1):R10–8. Available from: https://doi.org/10.1093/hmg/ddaa115

49 Karim HT. The Elusive “White Whale” of Treatment Response Prediction: Leveraging the Curse of Heterogeneity in Late-Life Depression. Am J Geriatr Psychiatry Off J Am Assoc Geriatr Psychiatry. 2021 Apr 9;S1064-7481(21)00288–8.

50 Devi G, Scheltens P. Heterogeneity of Alzheimer’s disease: consequence for drug trials? Alzheimers Res Ther [Internet]. 2018 Dec 19 [cited 2019 Oct 2];10. Available from: https://www.ncbi.nlm.nih.gov/pmc/articles/PMC6300886/

51 Butters MA, Young JB, Lopez O, Aizenstein HJ, Mulsant BH, Reynolds CF, et al. Pathways linking late-life depression to persistent cognitive impairment and dementia. Dialogues Clin Neurosci. 2008;10(3):345–57.

52 Koenig AM, Bhalla RK, Butters MA. Cognitive functioning and late-life depression. J Int Neuropsychol Soc JINS. 2014 May;20(5):461–7.

53 Livingston G, Huntley J, Sommerlad A, Ames D, Ballard C, Banerjee S, et al. Dementia prevention, intervention, and care: 2020 report of the Lancet Commission. Lancet Lond Engl. 2020 Aug 8;396(10248):413–46.

54 COVID-19 Mental Disorders Collaborators. Global prevalence and burden of depressive and anxiety disorders in 204 countries and territories in 2020 due to the COVID-19 pandemic. Lancet Lond Engl. 2021 Nov 6;398(10312):1700–12.

55 Kilicoglu H, Rosemblat G, Fiszman M, Shin D. Broad-coverage biomedical relation extraction with SemRep. BMC Bioinformatics [Internet]. 2020 May 14 [cited 2020 May 26];21(1):188. Available from: https://doi.org/10.1186/s12859-020-3517-7

56 Kilicoglu H, Rosemblat G, Fiszman M, Rindflesch TC. Constructing a semantic predication gold standard from the biomedical literature. BMC Bioinformatics. 2011 Dec 20;12:486.

57 Rindflesch TC, Blake CL, Fiszman M, Kilicoglu H, Rosemblat G, Schneider J, et al. Informatics Support for Basic Research in Biomedicine. ILAR J [Internet]. 2017 Jul 1 [cited 2019 Aug 5];58(1):80–9. Available from: https://www.ncbi.nlm.nih.gov/pmc/articles/PMC5886329/

58 Gyori BM, Bachman JA, Subramanian K, Muhlich JL, Galescu L, Sorger PK. From word models to executable models of signaling networks using automated assembly. Mol Syst Biol [Internet]. 2017 Nov 24 [cited 2020 Aug 14];13(11). Available from: https://www.ncbi.nlm.nih.gov/pmc/articles/PMC5731347/

59 VanderWeele T. Explanation in Causal Inference: Methods for Mediation and Interaction [Internet]. Oxford University Press; 2015. Available from: https://books.google.com/books?id=K6cgBgAAQBAJ

60 Aalen OO, Roysland K, Gran JM, Ledergerber B. Causality, mediation and time: a dynamic viewpoint. J R Stat Soc Ser A Stat Soc. 2012 Oct;175(4):831–61.

61 Richiardi L, Bellocco R, Zugna D. Mediation analysis in epidemiology: methods, interpretation and bias. Int J Epidemiol. 2013 Oct;42(5):1511–9.

62 Hristovski D, Friedman C, Rindflesch TC, Peterlin B. Exploiting semantic relations for literature-based discovery. AMIA Annu Symp Proc AMIA Symp. 2006;349–53.

63 Malec SA, Wei P, Bernstam EV, Boyce RD, Cohen T. Using computable knowledge mined from the literature to elucidate confounders for EHR-based pharmacovigilance. J Biomed Inform [Internet]. 2021 May 1 [cited 2021 Mar 31];117:103719. Available from: https://www.sciencedirect.com/science/article/pii/S1532046421000484

64 Kilicoglu H, Shin D, Fiszman M, Rosemblat G, Rindflesch TC. SemMedDB: a PubMed-scale repository of biomedical semantic predications. Bioinforma Oxf Engl. 2012 Dec 1;28(23):3158–60.

65 Cafasso M. noxdafox/clipspy [Internet]. 2020 [cited 2020 Aug 8]. Available from: https://github.com/noxdafox/clipspy

66 CLIPS: A Tool for Building Expert Systems [Internet]. [cited 2022 Mar 20]. Available from: http://www.clipsrules.net/

67 Aronson AR. Effective mapping of biomedical text to the UMLS Metathesaurus: the MetaMap program. Proc AMIA Symp. 2001;17–21.

68 Aronson AR, Lang FM. An overview of MetaMap: historical perspective and recent advances. J Am Med Inform Assoc [Internet]. 2010 May [cited 2017 Jul 21];17(3):229–36. Available from: https://academic.oup.com/jamia/article-lookup/doi/10.1136/jamia.2009.002733

69 Semantic Types and Groups - MetaMap documentation [Internet]. [cited 2022 Feb 21]. Available from: https://lhncbc.nlm.nih.gov/ii/tools/MetaMap/documentation/SemanticTypesAndGroups.html

70 Sharp R, Pyarelal A, Gyori B, Alcock K, Laparra E, Valenzuela-Escárcega MA, et al. Eidos, INDRA, & Delphi: From Free Text to Executable Causal Models. Proc 2019 Conf North Am Chapter Assoc Comput Linguist Demonstr. 2019;6.

71 clulab/eidos [Internet]. Computational Language Understanding Lab (CLU Lab) at University of Arizona; 2020 [cited 2020 Aug 14]. Available from: https://github.com/clulab/eidos

72 Bazrgar M, Khodabakhsh P, Mohagheghi F, Prudencio M, Ahmadiani A. Brain microRNAs dysregulation: Implication for missplicing and abnormal post-translational modifications of tau protein in Alzheimer’s disease and related tauopathies. Pharmacol Res. 2020;155:104729.

73 Marcelli S, Corbo M, Iannuzzi F, Negri L, Blandini F, Nistico R, et al. The Involvement of Post-Translational Modifications in Alzheimer’s Disease. Curr Alzheimer Res. 2018 22;15(4):313–35.

74 Schaffert LN, Carter WG. Do Post-Translational Modifications Influence Protein Aggregation in Neurodegenerative Diseases: A Systematic Review. Brain Sci [Internet]. 2020 Apr 11 [cited 2020 Oct 13];10(4). Available from: https://www.ncbi.nlm.nih.gov/pmc/articles/PMC7226274/

75 Rindflesch TC, Fiszman M. The interaction of domain knowledge and linguistic structure in natural language processing: interpreting hypernymic propositions in biomedical text. J Biomed Inform [Internet]. 2003 Dec [cited 2020 May 28];36(6):462–77. Available from: https://linkinghub.elsevier.com/retrieve/pii/S1532046403001175

76 Relation Ontology [Internet]. [cited 2022 Mar 20]. Available from: https://obofoundry.org/ontology/ro.html

77 Fine K. Towards a Theory of Part. J Philos [Internet]. 2010 Nov 1 [cited 2023 Feb 25];107(11):559–89. Available from: https://www.pdcnet.org/pdc/bvdb.nsf/purchase?openform&fp=jphil&id=jphil_2010_0107_0011_0559_0589

78 Kahn CE. Transitive closure of subsumption and causal relations in a large ontology of radiological diagnosis. J Biomed Inform [Internet]. 2016 Jun 1 [cited 2021 Nov 12];61:27–33. Available from: https://www.sciencedirect.com/science/article/pii/S1532046416300065

79 oborel/obo-relations [Internet]. oborel; 2020 [cited 2020 Aug 8]. Available from: https://github.com/oborel/obo-relations

80 Horrocks I, Patel-schneider PF. Knowledge Representation and Reasoning on the Semantic Web: OWL.

81 OWL Web Ontology Language Reference [Internet]. [cited 2022 Mar 28]. Available from: https://www.w3.org/TR/owl-ref/

82 NASA Technical Reports Server (NTRS) [Internet]. [cited 2020 Aug 8]. Available from: https://ntrs.nasa.gov/citations/19910014730

83 Beckers SV. The Transitivity and Asymmetry of Actual Causation. Open Access J Philos [Internet]. 2017;4. Available from: http://hdl.handle.net/2027/spo.12405314.0004.001

84 Callahan T. PheKnowLator [Internet]. 2019. Available from: https://doi.org/10.5281/zenodo.3401437

85 OWL - Semantic Web Standards [Internet]. [cited 2020 Aug 20]. Available from: https://www.w3.org/OWL/

86 v2 Data Sources · callahantiff/PheKnowLator Wiki [Internet]. GitHub. [cited 2022 Jun 1]. Available from: https://github.com/callahantiff/PheKnowLator

87 NetworkX — NetworkX documentation [Internet]. [cited 2020 Aug 21]. Available from: https://networkx.github.io/

88 Callahan TJ, Baumgartner WA, Bada M, Stefanski AL, Tripodi I, White EK, et al. OWL-NETS: Transforming OWL Representations for Improved Network Inference. Pac Symp Biocomput Pac Symp Biocomput [Internet]. 2018 [cited 2021 Aug 9];23:133–44. Available from: https://www.ncbi.nlm.nih.gov/pmc/articles/PMC5737627/

89 OpenLink Software: Virtuoso Homepage [Internet]. [cited 2022 Jun 7]. Available from: https://virtuoso.openlinksw.com/

90 13.5.2.2 Confounding and adjustment [Internet]. [cited 2023 Feb 15]. Available from: https://handbook-5-1.cochrane.org/chapter_13/13_5_2_2_confounding_and_adjustment.htm

91 Ashburner M, Ball CA, Blake JA, Botstein D, Butler H, Cherry JM, et al. Gene ontology: tool for the unification of biology. The Gene Ontology Consortium. Nat Genet. 2000 May;25(1):25–9.

92 Gene Ontology Resource [Internet]. Gene Ontology Resource. [cited 2022 Mar 20]. Available from: http://geneontology.org/

93 Human Phenotype Ontology [Internet]. [cited 2022 Mar 20]. Available from: https://hpo.jax.org/app/

94 SOD2 superoxide dismutase 2 [Homo sapiens (human)] - Gene - NCBI [Internet]. [cited 2022 May 28]. Available from: https://www.ncbi.nlm.nih.gov/gene/6648

95 Flynn JM, Melov S. SOD2 in Mitochondrial Dysfunction and Neurodegeneration. Free Radic Biol Med [Internet]. 2013 Sep [cited 2022 May 27];62:10.1016/j.freeradbiomed.2013.05.027. Available from: https://www.ncbi.nlm.nih.gov/pmc/articles/PMC3811078/

96 Wiener HW, Perry RT, Chen Z, Harrell LE, Go RCP. A polymorphism in SOD2 is associated with development of Alzheimer’s disease. Genes Brain Behav. 2007 Nov;6(8):770–5.

97 Nguyen M, Sabry R, St. John. Elizabeth J, Favetta LA. Bisphenol A and S, but Not F, Alter Oxidative Stress Levels in Spermatozoa. J Endocr Soc [Internet]. 2021 May 1 [cited 2022 Jun 5];5(Supplement_1):A483–4. Available from: https://doi.org/10.1210/jendso/bvab048.989

98 Braun: Early-life exposure to EDCs: role in childhood … - Google Scholar [Internet]. [cited 2022 Jun 6]. Available from: https://scholar.google.com/scholar_lookup?title=Early-life%20exposure%20to%20EDCs%3A%20role%20in%20childhood%20obesity%20and%20neurodevelopment&publication_year=2017&author=J.M.%20Braun

99 Choi YJ, Lee YA, Hong YC, Cho J, Lee KS, Shin CH, et al. Effect of prenatal bisphenol A exposure on early childhood body mass index through epigenetic influence on the insulin-like growth factor 2 receptor (IGF2R) gene. Environ Int [Internet]. 2020 Oct 1 [cited 2022 Jun 6];143:105929. Available from: https://www.sciencedirect.com/science/article/pii/S0160412020318845

100 Bisphenol A: How the Most Relevant Exposure Sources Contribute to Total Consumer Exposure - Von Goetz - 2010 - Risk Analysis - Wiley Online Library [Internet]. [cited 2022 Jun 6]. Available from: https://onlinelibrary.wiley.com/doi/full/10.1111/j.1539-6924.2009.01345.x?casa_token=g9CVKHGnY7EAAAAA%3ARUTZPsV8Fmu_6Qz259EJKD5PsDX5fbWC7wUCDA3sCcCZ3WY9rPuD46POLpkGjM26Ed6ndOSs28RA

101 MTHFR methylenetetrahydrofolate reductase [Homo sapiens (human)] - Gene - NCBI [Internet]. [cited 2022 May 28]. Available from: https://www.ncbi.nlm.nih.gov/gene/4524

102 Dwyer R, Skrobot OA, Dwyer J, Munafo M, Kehoe PG. Using Alzgene-like approaches to investigate susceptibility genes for vascular cognitive impairment. J Alzheimers Dis JAD. 2013;34(1):145–54.

103 Lemche E. Early Life Stress and Epigenetics in Late-onset Alzheimer’s Dementia: A Systematic Review. Curr Genomics [Internet]. 2018 Nov [cited 2021 Oct 2];19(7):522–602. Available from: https://www.ncbi.nlm.nih.gov/pmc/articles/PMC6194433/

104 Varga EA, Sturm AC, Misita CP, Moll S. Homocysteine and MTHFR Mutations. Circulation [Internet]. 2005 May 17 [cited 2022 May 28];111(19):e289–93. Available from: https://www.ahajournals.org/doi/10.1161/01.cir.0000165142.37711.e7

105 Saczynski JS, Beiser A, Seshadri S, Auerbach S, Wolf PA, Au R. Depressive symptoms and risk of dementia: the Framingham Heart Study. Neurology. 2010 Jul 6;75(1):35–41.

106 Sgroi S, Capper-Loup C, Paganetti P, Kaelin-Lang A. Enkephalin and dynorphin neuropeptides are differently correlated with locomotor hypersensitivity and levodopa-induced dyskinesia in parkinsonian rats. Exp Neurol. 2016 Jun;280:80–8.

107 PubChem. PDYN - prodynorphin (human) [Internet]. [cited 2022 May 29]. Available from: https://pubchem.ncbi.nlm.nih.gov/gene/PDYN/human

108 Kwintkiewicz J, Nishi Y, Yanase T, Giudice LC. Peroxisome Proliferator–Activated Receptor-γ Mediates Bisphenol A Inhibition of FSH-Stimulated IGF-1, Aromatase, and Estradiol in Human Granulosa Cells. Environ Health Perspect [Internet]. 2010 Mar [cited 2022 Feb 23];118(3):400–6. Available from: https://www.ncbi.nlm.nih.gov/pmc/articles/PMC2854770/

109 Pascual-Lucas M, Viana da Silva S, Di Scala M, Garcia-Barroso C, González-Aseguinolaza G, Mulle C, et al. Insulin-like growth factor 2 reverses memory and synaptic deficits in APP transgenic mice. EMBO Mol Med. 2014 Oct;6(10):1246–62.

110 Mellott TJ, Pender SM, Burke RM, Langley EA, Blusztajn JK. IGF2 ameliorates amyloidosis, increases cholinergic marker expression and raises BMP9 and neurotrophin levels in the hippocampus of the APPswePS1dE9 Alzheimer’s disease model mice. PloS One. 2014;9(4):e94287.

111 Forré P, Mooij JM. Markov Properties for Graphical Models with Cycles and Latent Variables [Internet]. arXiv; 2017 Oct [cited 2022 Jun 5]. Report No.: arXiv:1710.08775. Available from: http://arxiv.org/abs/1710.08775

112 Lauritzen SL, Richardson TS. Chain graph models and their causal interpretations. J R Stat Soc Ser B Stat Methodol [Internet]. 2002 [cited 2021 Oct 2];64(3):321–48. Available from: https://onlinelibrary.wiley.com/doi/abs/10.1111/1467-9868.00340

113 Mirza SS, Wolters FJ, Swanson SA, Koudstaal PJ, Hofman A, Tiemeier H, et al. 10-year trajectories of depressive symptoms and risk of dementia: a population-based study. Lancet Psychiatry. 2016 Jul;3(7):628–35.

114 Jang YJ, Kang C, Myung W, Lim SW, Moon YK, Kim H, et al. Additive interaction of mid- to late-life depression and cerebrovascular disease on the risk of dementia: a nationwide population-based cohort study. Alzheimers Res Ther [Internet]. 2021 [cited 2022 Mar 13];13. Available from: https://www.ncbi.nlm.nih.gov/labs/pmc/articles/PMC7968260/

115 Royall DR, Palmer RF. Alzheimer’s disease pathology does not mediate the association between depressive symptoms and subsequent cognitive decline. Alzheimers Dement J Alzheimers Assoc. 2013 May;9(3):318–25.

116 Singh-Manoux A, Dugravot A, Fournier A, Abell J, Ebmeier K, Kivimäki M, et al. Trajectories of Depressive Symptoms Before Diagnosis of Dementia. JAMA Psychiatry [Internet]. 2017 Jul [cited 2022 Aug 23];74(7):712–8. Available from: https://www.ncbi.nlm.nih.gov/pmc/articles/PMC5710246/

117 Brommelhoff JA, Gatz M, Johansson B, McArdle JJ, Fratiglioni L, Pedersen NL. Depression as a risk factor or prodromal feature for dementia? Findings in a population-based sample of Swedish twins. Psychol Aging. 2009 Jun;24(2):373–84.

118 Almeida OP, Hankey GJ, Yeap BB, Golledge J, Flicker L. Depression as a modifiable factor to decrease the risk of dementia. Transl Psychiatry [Internet]. 2017 May [cited 2022 Mar 13];7(5):e1117. Available from: https://www.ncbi.nlm.nih.gov/labs/pmc/articles/PMC5534958/

119 Stang PE, Ryan PB, Overhage JM, Schuemie MJ, Hartzema AG, Welebob E. Variation in choice of study design: findings from the Epidemiology Design Decision Inventory and Evaluation (EDDIE) survey. Drug Saf. 2013 Oct;36 Suppl 1:S15–25.

120 Patel CJ, Burford B, Ioannidis JPA. Assessment of vibration of effects due to model specification can demonstrate the instability of observational associations. J Clin Epidemiol. 2015 Sep;68(9):1046–58.

121 Das J, G K R. Post stroke depression: The sequelae of cerebral stroke. Neurosci Biobehav Rev. 2018 Jul;90:104–14.

122 Lewin-Richter A, Volz M, Jöbges M, Werheid K. Predictivity of Early Depressive Symptoms for Post-Stroke Depression. J Nutr Health Aging. 2015 Aug;19(7):754–8.

123 Medeiros GC, Roy D, Kontos N, Beach SR. Post-stroke depression: A 2020 updated review. Gen Hosp Psychiatry. 2020 Oct;66:70–80.

124 Lo Coco D, Lopez G, Corrao S. Cognitive impairment and stroke in elderly patients. Vasc Health Risk Manag. 2016;12:105–16.

125 Zhou J, Yu JT, Wang HF, Meng XF, Tan CC, Wang J, et al. Association between stroke and Alzheimer’s disease: systematic review and meta-analysis. J Alzheimers Dis JAD. 2015;43(2):479–89.

126 Vijayan M, Reddy PH. Stroke, Vascular Dementia, and Alzheimer’s Disease: Molecular Links. J Alzheimers Dis JAD. 2016 Sep 6;54(2):427–43.

127 Rindflesch TC, Libbus B, Hristovski D, Aronson AR, Kilicoglu H. Semantic Relations Asserting the Etiology of Genetic Diseases. AMIA Annu Symp Proc [Internet]. 2003 [cited 2023 Feb 14];2003:554–8. Available from: https://www.ncbi.nlm.nih.gov/pmc/articles/PMC1480275/

128 Steptoe A, Wikman A, Molloy GJ, Kaski JC. Anaemia and the development of depressive symptoms following acute coronary syndrome: longitudinal clinical observational study. BMJ Open. 2012;2(1):e000551.

129 Moehner S, Becker K, Lange JA, von Stockum S, Heinemann K. Risk of depression and anemia in users of hormonal endometriosis treatments: Results from the VIPOS study. Eur J Obstet Gynecol Reprod Biol. 2020 Aug;251:212–7.

130 Winchester LM, Powell J, Lovestone S, Nevado-Holgado AJ. Red blood cell indices and anaemia as causative factors for cognitive function deficits and for Alzheimer’s disease. Genome Med. 2018 Jun 28;10(1):51.

131 AGT angiotensinogen [Homo sapiens (human)] - Gene - NCBI [Internet]. [cited 2022 Jun 6]. Available from: https://www.ncbi.nlm.nih.gov/gene?Db=gene&Cmd=DetailsSearch&Term=183

132 Ajmal A, Gessert CE, Johnson BP, Renier CM, Palcher JA. Effect of angiotensin converting enzyme inhibitors and angiotensin receptor blockers on hemoglobin levels. BMC Res Notes [Internet]. 2013 Nov 4 [cited 2022 Jun 9];6:443. Available from: https://www.ncbi.nlm.nih.gov/pmc/articles/PMC4228356/

133 Faux NG, Rembach A, Wiley J, Ellis KA, Ames D, Fowler CJ, et al. An anemia of Alzheimer’s disease. Mol Psychiatry. 2014 Nov;19(11):1227–34.

134 Hare DJ, Doecke JD, Faux NG, Rembach A, Volitakis I, Fowler CJ, et al. Decreased plasma iron in Alzheimer’s disease is due to transferrin desaturation. ACS Chem Neurosci. 2015 Mar 18;6(3):398–402.

135 Mao Y, Liu X, Song Y, Zhai C, Zhang L. Data from: VEGF-A/VEGFR-2 and FGF-2/FGFR-1 but not PDGF-BB/PDGFR-β play important roles in promoting immature and inflammatory intraplaque angiogenesis [Internet]. Dryad; 2019 [cited 2022 Jun 9]. p. 266587 bytes. Available from: http://datadryad.org/stash/dataset/doi:10.5061/dryad.8tt0411

136 Harris R, Miners JS, Allen S, Love S. VEGFR1 and VEGFR2 in Alzheimer’s Disease. J Alzheimers Dis JAD. 2018;61(2):741–52.

137 Jeong SM, Shin DW, Lee JE, Hyeon JH, Lee J, Kim S. Anemia is associated with incidence of dementia: a national health screening study in Korea involving 37,900 persons. Alzheimers Res Ther. 2017 Dec 6;9(1):94.

138 Shafi M, Taufiq F, Mehmood H, Afsar S, Badar A. Relation between Depressive Disorder and Iron Deficiency Anemia among Adults Reporting to a Secondary Healthcare Facility: A Hospital-Based Case Control Study. J Coll Physicians Surg--Pak JCPSP. 2018 Jun;28(6):456–559.

139 Hosseini SR, Zabihi A, Ebrahimi SH, Jafarian Amiri SR, Kheirkhah F, Bijani A. The Prevalence of Anemia and its Association with Depressive Symptoms among Older Adults in North of Iran. J Res Health Sci [Internet]. 2018 Dec 3 [cited 2022 Jul 6];18(4):e00431. Available from: https://www.ncbi.nlm.nih.gov/pmc/articles/PMC6941631/

140 Vulser H, Wiernik E, Hoertel N, Thomas F, Pannier B, Czernichow S, et al. Association between depression and anemia in otherwise healthy adults. Acta Psychiatr Scand. 2016 Aug;134(2):150–60.

141 Linard M, Bezin J, Hucteau E, Joly P, Garrigue I, Dartigues JF, et al. Antiherpetic drugs: a potential way to prevent Alzheimer’s disease? Alzheimers Res Ther [Internet]. 2022 Jan 7 [cited 2022 Jul 19];14(1):3. Available from: https://doi.org/10.1186/s13195-021-00950-0

142 Moir RD, Lathe R, Tanzi RE. The antimicrobial protection hypothesis of Alzheimer’s disease. Alzheimers Dement J Alzheimers Assoc. 2018 Dec;14(12):1602–14.

143 Kumar DKV, Eimer WA, Tanzi RE, Moir RD. Alzheimer’s disease: the potential therapeutic role of the natural antibiotic amyloid-β peptide. Neurodegener Dis Manag. 2016 Oct;6(5):345–8.

144 Vijaya Kumar DK, Moir RD. The Emerging Role of Innate Immunity in Alzheimer’s Disease. Neuropsychopharmacol Off Publ Am Coll Neuropsychopharmacol. 2017 Jan;42(1):362.

145 Eimer WA, Vijaya Kumar DK, Shanmugam NKN, Rodriguez AS, Mitchell T, Washicosky KJ, et al. Alzheimer’s Disease-Associated β-Amyloid Is Rapidly Seeded by Herpesviridae to Protect against Brain Infection. Neuron [Internet]. 2018 Jul 11 [cited 2022 Jul 19];99(1):56–63.e3. Available from: https://www.ncbi.nlm.nih.gov/pmc/articles/PMC6075814/

146 Abbott A. Depression: the radical theory linking it to inflammation. Nature [Internet]. 2018 May 29 [cited 2022 Feb 28];557(7707):633–4. Available from: https://www.nature.com/articles/d41586-018-05261-3

147 Lee CH, Giuliani F. The Role of Inflammation in Depression and Fatigue. Front Immunol [Internet]. 2019 Jul 19 [cited 2022 Feb 28];10:1696. Available from: https://www.ncbi.nlm.nih.gov/pmc/articles/PMC6658985/

148 Kinney JW, Bemiller SM, Murtishaw AS, Leisgang AM, Salazar AM, Lamb BT. Inflammation as a central mechanism in Alzheimer’s disease. Alzheimers Dement Transl Res Clin Interv [Internet]. 2018 Sep 6 [cited 2022 Feb 28];4:575–90. Available from: https://www.ncbi.nlm.nih.gov/pmc/articles/PMC6214864/

149 Hampel H, Caraci F, Cuello AC, Caruso G, Nisticò R, Corbo M, et al. A Path Toward Precision Medicine for Neuroinflammatory Mechanisms in Alzheimer’s Disease. Front Immunol [Internet]. 2020 Mar 31 [cited 2021 May 12];11. Available from: https://www.ncbi.nlm.nih.gov/pmc/articles/PMC7137904/

150 Lyra e Silva NM, Gonçalves RA, Pascoal TA, Lima-Filho RAS, Resende E de PF, Vieira ELM, et al. Pro-inflammatory interleukin-6 signaling links cognitive impairments and peripheral metabolic alterations in Alzheimer’s disease. Transl Psychiatry [Internet]. 2021 Apr 28 [cited 2022 Feb 27];11(1):1–15. Available from: https://www.nature.com/articles/s41398-021-01349-z

151 Tani Y, Fujiwara T, Kondo K. Association Between Adverse Childhood Experiences and Dementia in Older Japanese Adults. JAMA Netw Open. 2020 Feb 5;3(2):e1920740.

152 Ciccarese P, Wu E, Wong G, Ocana M, Kinoshita J, Ruttenberg A, et al. The SWAN biomedical discourse ontology. J Biomed Inform. 2008 Oct;41(5):739–51.

153 Rosemblat G, Fiszman M, Shin D, Kilicoglu H. Towards a characterization of apparent contradictions in the biomedical literature using context analysis. J Biomed Inform. 2019;98:103275.

154 Alamri A. The Detection of Contradictory Claims in Biomedical Abstracts [Internet] [phd]. University of Sheffield; 2016 [cited 2019 Aug 15]. Available from: http://etheses.whiterose.ac.uk/15893/

155 Nordon G, Gottlieb L, Radinsky K. Chemical and Textual Embeddings for Drug Repurposing. :6.

156 Hsieh KL, Plascencia-Villa G, Lin KH, Perry G, Jiang X, Kim Y. Deep Learning for Alzheimer’s Disease Drug Repurposing using Knowledge Graph and Multi-level Evidence [Internet]. medRxiv; 2021 [cited 2022 May 2]. p. 2021.12.03.21267235. Available from: https://www.medrxiv.org/content/10.1101/2021.12.03.21267235v1

157 Malec SA, Wei P, Xu H, Bernstam EV, Myneni S, Cohen T. Literature-Based Discovery of Confounding in Observational Clinical Data. AMIA Annu Symp Proc AMIA Symp. 2016;2016:1920–9.

158 Swanson DR. Fish oil, Raynaud’s syndrome, and undiscovered public knowledge. Perspect Biol Med. 1986 Autumn;30(1):7–18.

159 Smalheiser NR. Rediscovering Don Swanson: the Past, Present and Future of Literature-Based Discovery. J Data Inf Sci Wars Pol [Internet]. 2017 Dec;2(4):43–64. Available from: http://www.ncbi.nlm.nih.gov/pmc/articles/PMC5771422/

160 Smalheiser NR, Swanson DR. Using ARROWSMITH: a computer-assisted approach to formulating and assessing scientific hypotheses. Comput Methods Programs Biomed. 1998 Nov;57(3):149–53.

161 Smalheiser NR. Literature-based discovery: Beyond the ABCs. J Am Soc Inf Sci Technol [Internet]. 2012 Feb [cited 2017 Jul 21];63(2):218–24. Available from: http://doi.wiley.com/10.1002/asi.21599

162 Ahlers CB, Hristovski D, Kilicoglu H, Rindflesch TC. Using the literature-based discovery paradigm to investigate drug mechanisms. AMIA Annu Symp Proc AMIA Symp. 2007 Oct 11;6–10.

163 Cohen T, Widdows D, Schvaneveldt RW, Davies P, Rindflesch TC. Discovering discovery patterns with Predication-based Semantic Indexing. J Biomed Inform. 2012 Dec;45(6):1049–65.

164 Cohen T, Widdows D, Schvaneveldt R, Rindflesch TC. Finding Schizophrenia’s Prozac Emergent Relational Similarity in Predication Space. In: Song D, Melucci M, Frommholz I, Zhang P, Wang L, Arafat S, editors. Quantum Interaction. Berlin, Heidelberg: Springer Berlin Heidelberg; 2011. p. 48–59.

165 Hristovski D, Kastrin A, Dinevski D, Rindflesch TC. Constructing a Graph Database for Semantic Literature-Based Discovery. Stud Health Technol Inform. 2015;216:1094.

166 Malec SA, Bernstam EV, Wei P, Boyce RD, Cohen T. Using computable knowledge mined from the literature to elucidate confounders for EHR-based pharmacovigilance. medRxiv [Internet]. 2020; Available from: https://www.medrxiv.org/content/early/2020/07/10/2020.07.08.20113035.1

167 Cohen T, Widdows D. Embedding of semantic predications. J Biomed Inform. 2017 Apr;68:150–66.

168 Cohen T, Schvaneveldt RW, Rindflesch TC. Predication-based Semantic Indexing: Permutations as a Means to Encode Predications in Semantic Space. AMIA Annu Symp Proc [Internet]. 2009;2009:114–8. Available from: http://www.ncbi.nlm.nih.gov/pmc/articles/PMC2815384/

169 Elsworth B, Dawe K, Vincent EE, Langdon R, Lynch BM, Martin RM, et al. MELODI: Mining Enriched Literature Objects to Derive Intermediates. Int J Epidemiol [Internet]. 2018 Apr 1 [cited 2020 Aug 14];47(2):369–79. Available from: https://academic.oup.com/ije/article/47/2/369/4803214

170 Cohen T, Whitfield GK, Schvaneveldt RW, Mukund K, Rindflesch T. EpiphaNet: An Interactive Tool to Support Biomedical Discoveries. J Biomed Discov Collab [Internet]. 2010;5:21–49. Available from: http://www.ncbi.nlm.nih.gov/pmc/articles/PMC2990276/

171 the Alzheimer’s Disease Neuroimaging Initiative, Shen X, Ma S, Vemuri P, Simon G. Challenges and Opportunities with Causal Discovery Algorithms: Application to Alzheimer’s Pathophysiology. Sci Rep [Internet]. 2020 Dec [cited 2020 Jun 1];103(1):2975. Available from: http://www.nature.com/articles/s41598-020-59669-x

172 Ahlers CB, Fiszman M, Demner-Fushman D, Lang FM, Rindflesch TC. Extracting semantic predications from Medline citations for pharmacogenomics. Pac Symp Biocomput Pac Symp Biocomput. 2007;209–20.

173 “MetaMap Team.” MetaMap - A Tool For Recognizing UMLS Concepts in Text [Internet]. 2015 [cited 2015 Jun 4]. Available from: http://metamap.nlm.nih.gov/

174 Harris ZS. A theory of language and information: a mathematical approach [Internet]. Clarendon Press; 1991. Available from: https://books.google.com/books?id=eT5iAAAAMAAJ

175 Friedman C, Kra P, Rzhetsky A. Two biomedical sublanguages: a description based on the theories of Zellig Harris. J Biomed Inform [Internet]. 2002 Aug 1 [cited 2023 Feb 15];35(4):222–35. Available from: https://www.sciencedirect.com/science/article/pii/S1532046403000121

176 Bernhardt PJ, Humphrey SM, Rindflesch TC. Determining Prominent Subdomains in Medicine. AMIA Annu Symp Proc [Internet]. 2005 [cited 2023 Feb 16];2005:46–50. Available from: https://www.ncbi.nlm.nih.gov/pmc/articles/PMC1560510/

177 Di Martino M, Fusco D, Colais P, Pinnarelli L, Davoli M, Perucci CA. Differential misclassification of confounders in comparative evaluation of hospital care quality: caesarean sections in Italy. BMC Public Health [Internet]. 2014 Oct 8 [cited 2023 Feb 14];14:1049. Available from: https://www.ncbi.nlm.nih.gov/pmc/articles/PMC4210510/

178 Liu W, Brookhart MA, Schneeweiss S, Mi X, Setoguchi S. Implications of M bias in epidemiologic studies: a simulation study. Am J Epidemiol. 2012 Nov 15;176(10):938–48.

179 VanderWeele TJ. On the relative nature of overadjustment and unnecessary adjustment. Epidemiol Camb Mass. 2009 Jul;20(4):496–9.

180 Naimi AI, Cole SR, Kennedy EH. An introduction to g methods. Int J Epidemiol [Internet]. 2017 Apr 1 [cited 2020 Jun 6];46(2):756–62. Available from: https://academic.oup.com/ije/article/46/2/756/2760169

181 Robins J. A new approach to causal inference in mortality studies with a sustained exposure period—application to control of the healthy worker survivor effect. Math Model [Internet]. 1986 Jan 1;7(9):1393–512. Available from: http://www.sciencedirect.com/science/article/pii/0270025586900886

182 Chesnaye NC, Stel VS, Tripepi G, Dekker FW, Fu EL, Zoccali C, et al. An introduction to inverse probability of treatment weighting in observational research. Clin Kidney J. 2022 Jan;15(1):14–20.

183 Austin PC, Stuart EA. Moving towards best practice when using inverse probability of treatment weighting (IPTW) using the propensity score to estimate causal treatment effects in observational studies. Stat Med [Internet]. 2015 Dec 10;34(28):3661–79. Available from: http://www.ncbi.nlm.nih.gov/pmc/articles/PMC4626409/

184 Cole SR, Hernán MA. Constructing Inverse Probability Weights for Marginal Structural Models. Am J Epidemiol [Internet]. 2008 Sep 15 [cited 2021 Jul 31];168(6):656–64. Available from: https://www.ncbi.nlm.nih.gov/pmc/articles/PMC2732954/

185 Tchetgen Tchetgen EJ, Glymour MM, Weuve J, Robins J. Specifying the correlation structure in inverse-probability-weighting estimation for repeated measures. Epidemiol Camb Mass. 2012 Jul;23(4):644–6.

186 Adams R, Ji Y, Wang X, Saria S. Learning Models from Data with Measurement Error: Tackling Underreporting. In: Proceedings of the 36th International Conference on Machine Learning [Internet]. PMLR; 2019 [cited 2023 Feb 9]. p. 61–70. Available from: https://proceedings.mlr.press/v97/adams19a.html

187 Lin HW, Chen YH. Adjustment for missing confounders in studies based on observational databases: Am J Epidemiol. 2014 Aug 1;180(3):308–17.

188 Munafò MR, Tilling K, Taylor AE, Evans DM, Davey Smith G. Collider scope: when selection bias can substantially influence observed associations. Int J Epidemiol [Internet]. 2018 Feb [cited 2022 Mar 10];47(1):226–35. Available from: https://www.ncbi.nlm.nih.gov/pmc/articles/PMC5837306/

189 Weuve J, Tchetgen Tchetgen EJ, Glymour MM, Beck TL, Aggarwal NT, Wilson RS, et al. Accounting for bias due to selective attrition: the example of smoking and cognitive decline. Epidemiol Camb Mass. 2012 Jan;23(1):119–28.

190 de Beurs E, Warmerdam L, Twisk J. Bias through selective inclusion and attrition: Representativeness when comparing provider performance with routine outcome monitoring data. Clin Psychol Psychother. 2019 Jul;26(4):430–9.

191 Hernán MA. Invited Commentary: Selection Bias Without Colliders. Am J Epidemiol [Internet]. 2017 Jun 1 [cited 2023 Feb 9];185(11):1048–50. Available from: https://www.ncbi.nlm.nih.gov/pmc/articles/PMC6664806/

192 Schwartz S, Gatto NM, Campbell UB. Transportability and causal generalization. Epidemiol Camb Mass. 2011 Sep;22(5):745–6.

193 Bareinboim E, Pearl J. Causal inference and the data-fusion problem. Proc Natl Acad Sci [Internet]. 2016 Jul 5 [cited 2020 Oct 12];113(27):7345–52. Available from: https://www.pnas.org/content/113/27/7345

194 Hripcsak G, Levine ME, Shang N, Ryan PB. Effect of vocabulary mapping for conditions on phenotype cohorts. J Am Med Inform Assoc JAMIA [Internet]. 2018 Nov 3 [cited 2020 Feb 20];25(12):1618–25. Available from: https://www.ncbi.nlm.nih.gov/pmc/articles/PMC6289550/

195 Lesko CR, Ackerman B, Webster-Clark M, Edwards JK. Target validity: Bringing treatment of external validity in line with internal validity. Curr Epidemiol Rep. 2020 Sep;7(3):117–24.

196 Eng CW, Glymour MM, Gilsanz P, Mungas DM, Mayeda ER, Meyer OL, et al. Do the Benefits of Educational Attainment for Late-life Cognition Differ by Racial/Ethnic Group?: Evidence for Heterogenous Treatment Effects in the Kaiser Healthy Aging and Diverse Life Experience (KHANDLE) Study. Alzheimer Dis Assoc Disord. 2021 Jun 1;35(2):106–13.

197 Vable AM, Nguyen TT, Rehkopf D, Glymour MM, Hamad R. Differential associations between state-level educational quality and cardiovascular health by race: Early-life exposures and late-life health. SSM - Popul Health. 2019 Aug;8:100418.

198 Sterne JA, Hernán MA, McAleenan A, Reeves BC, Higgins JP. Assessing risk of bias in a non-randomized study. In: Cochrane Handbook for Systematic Reviews of Interventions [Internet]. John Wiley & Sons, Ltd; 2019 [cited 2023 Feb 15]. p. 621–41. Available from: https://onlinelibrary.wiley.com/doi/abs/10.1002/9781119536604.ch25

199 Petersen JM, Barrett M, Ahrens KA, Murray EJ, Bryant AS, Hogue CJ, et al. The confounder matrix: A tool to assess confounding bias in systematic reviews of observational studies of etiology. Res Synth Methods. 2022 Mar;13(2):242–54.

200 Greenland S. Methods for epidemiologic analyses of multiple exposures: A review and comparative study of maximum-likelihood, preliminary-testing, and empirical-bayes regression. Stat Med [Internet]. 1993 [cited 2023 Feb 14];12(8):717–36. Available from: https://onlinelibrary.wiley.com/doi/abs/10.1002/sim.4780120802

201 Murata S, Ono R, Yasuda H, Tanemura R, Kido Y, Kowa H. Effect of a Combined Exercise and Cognitive Activity Intervention on Cognitive Function in Community-dwelling Older Adults: A Pilot Randomized Controlled Trial. Phys Ther Res. 2021;24(2):112–9.

202 Tager IB, Haight T, Sternfeld B, Yu Z, van Der Laan M. Effects of Physical Activity and Body Composition on Functional Limitation in the Elderly: Application of the Marginal Structural Model. Epidemiology [Internet]. 2004 Jul [cited 2023 Feb 14];15(4):479. Available from: https://journals.lww.com/epidem/Fulltext/2004/07000/Effects_of_Physical_Activity_and_Body_Composition.16.aspx

203 Kessing LV, Andersen PK. Does the risk of developing dementia increase with the number of episodes in patients with depressive disorder and in patients with bipolar disorder? J Neurol Neurosurg Psychiatry. 2004 Dec;75(12):1662–6.

204 Kessing LV, Forman JL, Andersen PK. Does lithium protect against dementia? Bipolar Disord. 2010 Feb;12(1):87–94.

205 Robins JM, Hernán MÁ, Brumback B. Marginal Structural Models and Causal Inference in Epidemiology. Epidemiology [Internet]. 2000 Sep [cited 2022 Mar 13];11(5):550–60. Available from: https://journals.lww.com/epidem/fulltext/2000/09000/marginal_structural_models_and_causal_inference_in.11.aspx

206 Shinozaki T, Suzuki E. Understanding Marginal Structural Models for Time-Varying Exposures: Pitfalls and Tips. J Epidemiol [Internet]. 2020 Sep 5 [cited 2023 Feb 15];30(9):377–89. Available from: https://www.ncbi.nlm.nih.gov/pmc/articles/PMC7429147/

207 Bhalla RK, Butters MA, Becker JT, Houck PR, Snitz BE, Lopez OL, et al. Patterns of mild cognitive impairment after treatment of depression in the elderly. Am J Geriatr Psychiatry Off J Am Assoc Geriatr Psychiatry. 2009 Apr;17(4):308–16.

208 Butters MA, Whyte EM, Nebes RD, Begley AE, Dew MA, Mulsant BH, et al. The nature and determinants of neuropsychological functioning in late-life depression. Arch Gen Psychiatry. 2004 Jun;61(6):587–95.

209 Ganguli M. Depression, cognitive impairment and dementia: Why should clinicians care about the web of causation? Indian J Psychiatry [Internet]. 2009 Jan [cited 2020 Mar 1];51(Suppl1):S29–34. Available from: https://www.ncbi.nlm.nih.gov/pmc/articles/PMC3038544/

210 Butters MA, Sweet RA, Mulsant BH, Ilyas Kamboh M, Pollock BG, Begley AE, et al. APOE is associated with age-of-onset, but not cognitive functioning, in late-life depression. Int J Geriatr Psychiatry. 2003;18(12):1075–81.

211 Safieh M, Korczyn AD, Michaelson DM. ApoE4: an emerging therapeutic target for Alzheimer’s disease. BMC Med [Internet]. 2019 Mar 20;17(1):64. Available from: https://doi.org/10.1186/s12916-019-1299-4

212 Qiu WQ, Zhu H, Dean M, Liu Z, Vu L, Fan G, et al. Amyloid-associated depression and ApoE4 allele: longitudinal follow-up for the development of Alzheimer’s disease. Int J Geriatr Psychiatry [Internet]. 2016 Mar [cited 2021 Sep 30];31(3):316–22. Available from: https://www.ncbi.nlm.nih.gov/pmc/articles/PMC4840849/

213 Hernán MA, Brumback BA, Robins JM. Estimating the causal effect of zidovudine on CD4 count with a marginal structural model for repeated measures. Stat Med. 2002 Jun 30;21(12):1689–709.

214 Chesnaye NC, Stel VS, Tripepi G, Dekker FW, Fu EL, Zoccali C, et al. An introduction to inverse probability of treatment weighting in observational research. Clin Kidney J. 2022 Jan;15(1):14–20.

215 Robins JM, Rotnitzky A, Zhao LP. Estimation of Regression Coefficients When Some Regressors are not Always Observed. J Am Stat Assoc [Internet]. 1994 Sep 1 [cited 2020 Nov 4];89(427):846–66. Available from: https://doi.org/10.1080/01621459.1994.10476818

216 Barnes DE, Yaffe K. The projected effect of risk factor reduction on Alzheimer’s disease prevalence. Lancet Neurol. 2011 Sep;10(9):819–28.

217 Ding P, Miratrix L. To Adjust or Not to Adjust? Sensitivity Analysis of M-Bias and Butterfly-Bias. ArXiv14080324 Math Stat [Internet]. 2014 Aug 1 [cited 2019 Oct 19]; Available from: http://arxiv.org/abs/1408.0324

218 Flanders WD, Ye D. Limits for the Magnitude of M-bias and Certain Other Types of Structural Selection Bias. Epidemiol Camb Mass. 2019 Jul;30(4):501–8.

219 Greenland S. Quantifying biases in causal models: classical confounding vs collider-stratification bias. Epidemiol Camb Mass. 2003 May;14(3):300–6.

220 Groenwold RHH, Palmer TM, Tilling K. To Adjust or Not to Adjust? When a “Confounder” Is Only Measured After Exposure. Epidemiol Camb Mass [Internet]. 2021 Mar [cited 2021 Oct 11];32(2):194–201. Available from: https://www.ncbi.nlm.nih.gov/pmc/articles/PMC7850592/

221 Kennedy EH, Balakrishnan S. Discussion of “Data-driven confounder selection via Markov and Bayesian networks” by Jenny Häggström. Biometrics [Internet]. 2018 [cited 2022 Aug 1];74(2):399–402. Available from: https://onlinelibrary.wiley.com/doi/abs/10.1111/biom.12787

222 Ioannidis JPA, Tan YJ, Blum MR. Limitations and Misinterpretations of E-Values for Sensitivity Analyses of Observational Studies. Ann Intern Med [Internet]. 2019 Jan 1 [cited 2021 Apr 18];170(2):108–11. Available from: http://www.acpjournals.org/doi/10.7326/M18-2159

223 Tennant PWG, Murray EJ, Arnold KF, Berrie L, Fox MP, Gadd SC, et al. Use of directed acyclic graphs (DAGs) to identify confounders in applied health research: review and recommendations. Int J Epidemiol [Internet]. 2021 Apr 1 [cited 2021 May 30];50(2):620–32. Available from: https://doi.org/10.1093/ije/dyaa213

224 Textor J, Zander B van der, Ankan A. dagitty: Graphical Analysis of Structural Causal Models [Internet]. 2021 [cited 2021 Aug 3]. Available from: https://CRAN.R-project.org/package=dagitty

225 Wise J, de Barron AG, Splendiani A, Balali-Mood B, Vasant D, Little E, et al. Implementation and relevance of FAIR data principles in biopharmaceutical R&D. Drug Discov Today [Internet]. 2019 Apr 1 [cited 2021 Jan 10];24(4):933–8. Available from: http://www.sciencedirect.com/science/article/pii/S1359644618303039

226 Fang Y, He W, Hu X, Wang H. A method for sample size calculation via E-value in the planning of observational studies. Pharm Stat [Internet]. [cited 2020 Sep 13];n/a(n/a). Available from: http://onlinelibrary.wiley.com/doi/abs/10.1002/pst.2064

227 Localio AR, Stack CB, Griswold ME. Sensitivity Analysis for Unmeasured Confounding: E-Values for Observational Studies. Ann Intern Med. 2017 Aug 15;167(4):285–6.

228 von Elm E, Altman DG, Egger M, Pocock SJ, Gøtzsche PC, Vandenbroucke JP, et al. The Strengthening the Reporting of Observational Studies in Epidemiology (STROBE) statement: guidelines for reporting observational studies. Lancet Lond Engl. 2007 Oct 20;370(9596):1453–7.

229 Cornfield J. A Statistical Problem Arising from Retrospective Studies. Proc Third Berkeley Symp Math Stat Probab Vol 4 Contrib Biol Probl Health [Internet]. 1956 Jan 1 [cited 2021 May 1]; 135–48. Available from: https://www.projecteuclid.org/ebooks/berkeley-symposium-on-mathematical-statistics-and-probability/Proceedings-of-the-Third-Berkeley-Symposium-on-Mathematical-Statistics-and/chapter/A-Statistical-Problem-Arising-from-Retrospective-Studies/bsmsp/1200502552

230 Vandenbroucke JP. Commentary: ‘Smoking and lung cancer’—the embryogenesis of modern epidemiology. Int J Epidemiol [Internet]. 2009 Oct 1 [cited 2021 May 10];38(5):1193–6. Available from: https://doi.org/10.1093/ije/dyp292

231 VanderWeele TJ. Are Greenland, Ioannidis and Poole opposed to the Cornfield conditions? A defence of the E-value. Int J Epidemiol [Internet]. 2021 Oct 13 [cited 2022 Jul 10];51(2):364–71. Available from: https://www.ncbi.nlm.nih.gov/pmc/articles/PMC9082787/

232 Forrest CB, McTigue KM, Hernandez AF, Cohen LW, Cruz H, Haynes K, et al. PCORnet® 2020: current state, accomplishments, and future directions. J Clin Epidemiol. 2021 Jan;129:60–7.

233 Fleurence RL, Curtis LH, Califf RM, Platt R, Selby JV, Brown JS. Launching PCORnet, a national patient-centered clinical research network. J Am Med Inform Assoc [Internet]. 2014 Jul 1 [cited 2017 Jul 2];21(4):578–82. Available from: https://academic.oup.com/jamia/article/21/4/578/2909226/Launching-PCORnet-a-national-patient-centered

234 All of Us Research Program Investigators, Denny JC, Rutter JL, Goldstein DB, Philippakis A, Smoller JW, et al. The “All of Us” Research Program. N Engl J Med. 2019 Aug 15;381(7):668–76.

