## Supplementary material for "Causal feature selection using a knowledge graph combining structured knowledge from the biomedical literature and ontologies: a use case studying depression as a risk factor for Alzheimer’s disease": File I. Appendix_A.docx. Background S1. Logical properties.

### Supplementary Material Appendix A.

**File I.** *Appendix_A.docx*. **Background S2.** Logical properties.

#### Background S2. Logical properties.

For instance, the transitive property applies to predicates that express a relationship between three or more entities. If a predicate has the *transitive* property, a relationship between A and B, and B and C, implies a relationship between A and C. The **CAUSES** predicate has the transitive property. If A **CAUSES** B, and B **CAUSES** C, we can infer that A **CAUSES** C. Similarly, the **PART_OF** predicate also has the transitive property. If A is part of B, and B is part of C, we can infer that A is part of C.

Symmetry applies to predicates that express a bidirectional relationship between two entities. If a predicate has the *symmetrical* property, the relationship between A and B is the same as the relationship between B and A. For example, the **COEXISTS_WITH**, predicate has the symmetrical property. If A coexists with B, we can infer that B **COEXISTS_WITH** A.

Reflexivity applies to predicates where an entity is related to itself. The asymmetric property applies to predicates that express a relationship between two entities, but the relationship is not bidirectional. If a predicate has the *asymmetric property*, the relationship between A and B is not the same as the relationship between B and A. For example, the **PRECEDES** predicate has the *asymmetric property*. If A **PRECEDES** B, we cannot infer that B **PRECEDES** A.

If a predicate has the reflexive property, every entity is related to itself. For instance, the **PART_OF** predicate has the reflexive property. Every entity is part of itself.

The inverse property in terms of causal, associative, and subsumptive relationships allows us to express a relationship between two entities in the opposite direction. For example, the inverse property of the **CAUSES** predicate is the **CAUSED_BY** predicate, which expresses the inverse relationship. If "A **CAUSES** B," then we can also say that "B **CAUSED_BY** A". Likewise, for the **PART_OF** predicate, the inverse property is the **HAS_PART** predicate. If "A **PART_OF** B," then we can also say that "B **HAS_PART** A," which means that "B" has "A" as a part.

However, in certain cases, a predicate is its own inverse predicate. Consider the symmetrical predicate **COEXISTS_WITH**, which expresses the inverse relationship between two entities that are frequently found together and is itself its own predicate.

**References**

1. Petersen ML, van der Laan MJ. Causal Models and Learning from Data. Epidemiol Camb Mass [Internet]. 2014 May [cited 2020 Sep 14];25(3):418–26. Available from: https://www.ncbi.nlm.nih.gov/pmc/articles/PMC4077670/
2. Grätz M. When Less Conditioning Provides Better Estimates: Overcontrol and Collider Bias in Research on Intergenerational Mobility [Internet]. Working Paper Series. Stockholm University, Swedish Institute for Social Research; 2019 Jun [cited 2020 Jun 16]. (Working Paper Series). Report No.: 2/2019. Available from: https://ideas.repec.org/p/hhs/sofiwp/2019_002.html
3. Schisterman EF, Cole SR, Platt RW. Overadjustment Bias and Unnecessary Adjustment in Epidemiologic Studies. Epidemiol Camb Mass. 2009 Jul;20(4):488–95.
4. Karim ME, Tremlett H, Zhu F, Petkau J, Kingwell E. Dealing with Treatment-confounder Feedback and Sparse Follow-up in Longitudinal studies - Application of a Marginal Structural Model in a Multiple Sclerosis Cohort. Am J Epidemiol. 2020 Oct 30;
5. Kennedy EH, Balakrishnan S. Discussion of “Data-driven confounder selection via Markov and Bayesian networks” by Jenny H\"aggstr\"om. 2017 Oct 31 [cited 2022 Aug 1]; Available from: https://arxiv.org/abs/1710.11566v1
6. Hernan MA, Hernandez-Diaz S, Werler MM, Mitchell AA. Causal knowledge as a prerequisite for confounding evaluation: an application to birth defects epidemiology. Am J Epidemiol. 2002 Jan 15;155(2):176–84.
7. VanderWeele TJ, Shpitser I. On the definition of a confounder. Ann Stat. 2013 Feb;41(1):196–220.
8. Greenland S, Morgenstern H. Confounding in health research. Annu Rev Public Health. 2001;22:189–212.
9. Greenland S, Robins JM. Identifiability, exchangeability, and epidemiological confounding. Int J Epidemiol. 1986 Sep;15(3):413–9.
10. VanderWeele TJ. Principles of confounder selection. Eur J Epidemiol [Internet]. 2019 [cited 2019 Aug 20];34(3):211–9. Available from: https://www.ncbi.nlm.nih.gov/pmc/articles/PMC6447501/
11. Pearl J. The Mathematics of Causal Inference. In: Proceedings of the 17th ACM SIGKDD International Conference on Knowledge Discovery and Data Mining [Internet]. New York, NY, USA: ACM; 2011 [cited 2018 Jun 16]. p. 5–5. (KDD ’11). Available from: http://doi.acm.org/10.1145/2020408.2020416
12. Reichenbach H, Reichenbach M. The Direction of Time [Internet]. University of California Press; 1991. (Philosophy (University of California (Los Ángeles)))). Available from: https://books.google.com/books?id=LTaU5JTj3mUC
13. Hernán MA, Robins JM. Causal Inference: What If. :310.
14. VanderWeele T. Explanation in Causal Inference: Methods for Mediation and Interaction [Internet]. Oxford University Press; 2015. Available from: https://books.google.com/books?id=K6cgBgAAQBAJ
15. Baron RM, Kenny DA. The moderator–mediator variable distinction in social psychological research: Conceptual, strategic, and statistical considerations. J Pers Soc Psychol. 1986;51(6):1173–82.
16. Sackett DL. Bias in analytic research. J Chronic Dis. 1979;32(1–2):51–63.
17. Elwert F, Winship C. Endogenous Selection Bias: The Problem of Conditioning on a Collider Variable. Annu Rev Sociol [Internet]. 2014 Jul;40:31–53. Available from: https://www.ncbi.nlm.nih.gov/pubmed/30111904
18. Williams TC, Bach CC, Matthiesen NB, Henriksen TB, Gagliardi L. Directed acyclic graphs: a tool for causal studies in paediatrics. Pediatr Res [Internet]. 2018 Oct [cited 2021 Feb 17];84(4):487–93. Available from: https://www.nature.com/articles/s41390-018-0071-3

19. Krieger N, Davey Smith G. The tale wagged by the DAG: broadening the scope of causal inference and explanation for epidemiology. Int J Epidemiol. 2016 Dec 1;45(6):1787–808.
20. Tennant PWG, Murray EJ, Arnold KF, Berrie L, Fox MP, Gadd SC, et al. Use of directed acyclic graphs (DAGs) to identify confounders in applied health research: review and recommendations. Int J Epidemiol [Internet]. 2021 Apr 1 [cited 2021 May 30];50(2):620–32. Available from: https://doi.org/10.1093/ije/dyaa213
21. Pearl J. Causal Diagrams for Empirical Research. Biometrika [Internet]. 1995 [cited 2020 May 17];82(4):669–88. Available from: www.jstor.org/stable/2337329
