## Supplementary material for "Causal feature selection using a knowledge graph combining structured knowledge from the biomedical literature and ontologies: a use case studying depression as a risk factor for Alzheimer’s disease": File_II-AppendixB-Details

#### Supplementary Material Appendix B.

**File** II. ***Appendix_B.docx***.

### Contents:

**Listing S1.** PubMed scoping query.

**Table S1.** Mapping of machine reading predicates to RO and BFO ontology targets.

**Table S2.** UMLS CUI definitions for depression and AD.

**Table S3.** Breakdown of Concepts (Count) by Ontology.

**Table S4.** Comparison of confounders - search results belonging to the gene or enzyme, protein, or semantic categories. Comparison of search results belonging to the gene or enzyme semantic categories. Bold indicates an exact match between the knowledge graph and the complete literature. Italicized bold signifies a closely related synonym. Italicized bold with an asterisk means that it was identified but that the match is in the opposite table (e.g., that a geneProtein mapping has a match in the drug table).

**Table S5.** Comparison of confounders - search results belonging to the drug or hormone semantic categories.

**Table S6.** Comparison with variables organized in rows by role (confounder, collider, or mediator) and columns organized by (knowledge graph (KG) and the two subsets of SemMedDB. The subsets (restricted to articles published in 2010 or after) include MachineReadingDB$_{\text{Scoped}}$ and SemMedDB$_{\text{Complete}}$. Concepts in bold appear in both the KG and SemMedDB$_{\text{Complete}}$. Underlined concepts appear in both SemMedDBs. To save space, we restrict our analysis of concepts retrieved to the semantic categories relating to findings and phenotypes. Strikethrough indicates that the variable appears in either the KG or SemMedDB$_{\text{Complete}}$ but not in the identical role(s).

**Table S7.** Comparison of search results belonging to the semantic categories of findings and phenotypes organized in rows by variable role, including role combinations ("Confounders only," "Colliders only," or "Mediators only") and columns organized by (knowledge graph (KG) and SemMedDB$_{\text{Complete}}$. We ignore MachineReadingDB$_{\text{Scoped}}$ for this analysis since it did not yield colliders or mediators. Concepts in bold appear in both the KG and SemMedDB$_{\text{Complete}}$. Closely related hypernymic/hyponymic concepts (near matched) are italicized in bold. Concepts are underlined if they appear in both SemMedDBs. A strikethrough (e.g., ~~anemia~~) indicates disagreement between knowledge bases.

**Figure S1.** Venn diagram for simple (not combined) showing complete KG and SemMedDB results.

**Figure S2.** Venn diagram for combined roles showing complete KG and SemMedDB results.

—------------------

**Listing S1.** PubMed scoping query.

The query used to search PubMed for the scoped subset of literature was as follows:

> ("alzheimer disease"[MeSH Terms] OR ("alzheimer"[All Fields] AND "disease"[All Fields]) OR "alzheimer disease"[All Fields] OR ("alzheimer's"[All Fields] AND "disease"[All Fields]) OR "alzheimer's disease"[All Fields]) AND (("risk"[MeSH Terms] OR "risk"[All Fields]) OR ("protective factors"[MeSH Terms] OR ("protective"[All Fields] AND "factors"[All Fields]) OR "protective factors"[All Fields]) OR ("risk assessment"[MeSH Terms] OR ("risk"[All Fields] AND "assessment"[All Fields]) OR "risk assessment"[All Fields]) OR ("adverse outcome pathways"[MeSH Terms] OR ("adverse"[All Fields] AND "outcome"[All Fields] AND "pathways"[All Fields]) OR "adverse outcome pathways"[All Fields]) OR ("risk factors"[MeSH Terms] OR ("risk"[All Fields] AND "factors"[All Fields]) OR "risk factors"[All Fields]) OR ("uncertainty"[MeSH Terms] OR "uncertainty"[All Fields]) OR ("prevention and control"[Subheading] OR ("prevention"[All Fields] AND "control"[All Fields]) OR "prevention and control"[All Fields] OR "prevention"[All Fields])) AND fha[FILTER] AND y_10[FILTER] AND humans[FILTER]

**Table S1.** The mapping of machine reading predicate sources to RO and BFO ontology targets.

| Machine reading system | Predicates from machine reading systems | RO/BFO predicate |
|---|---|---|
| **SemRep** | AFFECTS | RO_0002211 |
| | ASSOCIATED_WITH | RO_0002610 |
| | AUGMENTS | augments |
| | CAUSES | RO_0002501 |
| | COEXISTS_WITH | RO_0002490 |
| | COMPLICATES | complicates |
| | DISRUPTS | disrupts |
| | INHIBITS | RO_0002212 |
| | INTERACTS_WITH | RO_0002434 |
| | PART_OF | RO_0000050 |
| | PRECEDES | RO_0000063 |
| | PREDISPOSES | RO_0003302 |
| | PREVENTS | RO_0002599 |
| | STIMULATES | RO_0002213 |
| | TREATS | RO_0002606 |
| **INDRA (EIDOS)** | influence | RO_0002501 |
| | association | RO_0002610 |
| **INDRA (REACH)** | modification | RO_0002436 |
| | regulateactivity | RO_0002436 |
| | regulateamount | RO_0002436 |
| | phosphorylation | RO_0002447 |
| | dephosphorylation | RO_0002436 |
| | ubiquitination | RO_0002436 |
| | modification | RO_0002436 |
| | regulateactivity | RO_0002436 |
| | regulateamount | RO_0002436 |
| | phosphorylation | RO_0002447 |
| | dephosphorylation | RO_0002436 |
| | ubiquitination | RO_0002436 |
| | deubiquitination | RO_0002436 |
| | sumoylation | RO_0002436 |

| | | |
|---|---|---|
| | desumoylation | RO_0002436 |
| | hydroxylation | RO_0002436 |
| | dehydroxylation | RO_0002436 |
| | acetylation | RO_0002436 |
| | deacetylation | RO_0002436 |
| | glycosylation | RO_0002436 |
| | deglycosylation | RO_0002436 |
| | farnesylation | RO_0002436 |
| | defarnesylation | RO_0002436 |
| | geranylgeranylation | RO_0002436 |
| | degeranylgeranylation | RO_0002436 |
| | palmitoylation | RO_0002436 |
| | depalmitoylation | RO_0002436 |
| | myristoylation | RO_0002436 |
| | demyristoylation | RO_0002436 |
| | ribosylation | RO_0002436 |
| | deribosylation | RO_0002436 |
| | methylation | RO_0002436 |
| | demethylation | RO_0002436 |
| | activation | RO_0002429 |
| | inhibition | RO_0002212 |
| | increaseamount | RO_0002429 |
| | decreaseamount | RO_0002212 |

**Table S2.** UMLS CUI definitions for depression and AD.

| **Depression** | |
|---|---|
| **Concept Name** | **CUI** |
| Depressive disorder | C0011581 |
| Mild depression | C0588006 |
| Major Depressive Disorder, Recurrent, Unspecified | C0154409 |
| Treatment Resistant Depression | C0871546 |
| Major Depressive Disorder, Single Episode, Unspecified | C0024517 |
| Major Depressive Disorder | C1269683 |
| Severe depression | C0588008 |

| | |
|---|---|
| Unipolar Depression | C0041696 |
| Recurrent depression | C0221480 |
| Major depression, single episode | C0024517 |
| **Alzheimer's Disease** | |
| Alzheimer's disease | C0002395 |
| Alzheimer's disease, familial | C0276496 |
| Alzheimer's disease, late-onset | C0494463 |

**Table S3.** Breakdown of Concepts (Count) by Ontology.

| **Ontology** | **Count** |
|---|---|
| GO | 685 |
| UBERON | 133 |
| HP | 472 |
| CHEBI | 1068 |
| PW | 5 |
| CL | 20 |
| DOID | 101 |
| CLO | 7 |
| EFO | 12 |
| PR | 639 |
| PATO | 16 |
| VO | 18 |
| BFO | 5 |
| NCBITaxon | 44 |
| OBI | 2 |
| RO | 1 |
| IAO | 3 |
| cellline#human | 1 |
| SO | 1 |
| NCIT | 1 |
| PRO | 1 |

**Table S4.** Comparison of confounders - search results belonging to the gene or enzyme, protein, or semantic categories. Comparison of search results belonging to the gene or enzyme semantic categories. Bold indicates an exact match between the knowledge graph and the complete literature. Italicized bold signifies a closely related synonym. Italicized bold with an asterisk means that it was identified but that the match is in the opposite table (e.g., that a geneProtein mapping has a match in the drug table).

| | Knowledge Graph | SemMedDB |
|---|---|---|
| **Confounders that are genes, proteins, or enzymes** | A2M gene\|A2M, **adiponectin**, alanineb, ***amino_acids (branched chain)***, ***angiotensin coverting enzyme inhibitors***, **apolipoprotein C-III**, **apolipoprotein E**, **apolipoproteins**, **APP protein (human)**, arachidonic acid, aromatase, aromatase, BIN1_gene, butyrylcholinesterase, cathepsin D, CD33 gene, clusterin, **connexins**, CS gene, epitopes, FRAP1 protein (human), FRMD4A_gene, **homocystein**, **interleukin-1 beta**, **interleukin-10**, **interleukin-6**, large gene, **leptin**, ***lipoproteins***, matrix metalioproteinase-3, **P-Glycoprotein\|ABCB1**, Presenilin-1, preslbumin, **valine**, **VEGF protein (human)** | acetylcholine, acetylcholinesterase, adenosine, adiponectin, adrbk1 gene\|grk2, alanine, alpha-fetoproteins, alpha7 nicotinic acetylcholine receptor, alteplase, aluminum, amino acids, amino acids, branched-chain, amyloid, amyloid beta, ***angiotensin ii***, antibiotics, **apolipoprotein e**, ***apolipoprotein e-4\|apoe***, **apolipoprotein e\|apoe**, ***apolipoprotein e4***, ***app gene***, **app protein (human)**, arginine, arylesterase, bdnf gene\|bdnf, bilirubin, binding protein, brain natriuretic peptide, brain-derived neurotrophic factor, brain-derived neurotrophic factor\|bdnf, c-reactive protein, ca1\|s100a10, ca2, cacna1d, cadmium, calcium, calcium channel, calcium ion, calmodulin-dependent protein kinase ii\|camk2g, cancer-predisposing gene, candidate disease gene, cannabinoid receptor, carbamylated erythropoietin, caspase-1, cd69 protein, human\|cd69, cell adhesion molecules, ***ceramides****, chemokine, **cholesterol**, cholinergic system, clock gene\|clock, **connexins**, copper, corticotropin-releasing hormone, cpg islands, cyclic amp-responsive dna-binding protein\|creb1, cyclooxygenase 2, cycloserine, cyp2d6 gene\|cyp2d6, cysteine, cytokine, disc1 gene\|disc1, dna, mitochondrial, docosahexaenoic acids, dopamine, dopamine transporter, eicosapentaenoic acid, elavl2 gene\|elavl2, elk3 gene\|elk3\|kcnh8, endocannabinoids, epha4 gene\|epha4, epidermal growth factor\|egf, estrogen receptor alpha, fas, fatty acids, fatty acids, monounsaturated, fatty acids, omega-3, fgfr3 gene\|fgfr3, fto, functional rna, gabpa\|nfe2l2, gamma-aminobutyric acid, gene clusters, ghrelin, gja1 gene\|gja1, glial cell-line derived neurotrophic factor, glucose, glutamate, glutamate receptor, glutamates, glutathione, glycogen synthase kinase 3 beta, growth factor, gtpase-activating proteins, ***high density lipoprotein cholesterol***, histamine, **homocysteine**, homologous gene, human leukocyte interferon\|ifna1, hyperforin, ***il17a protein***, human\|il17a, impact gene, inflammasomes, insulin, insulin-like growth factor i, **interleukin-1 (beta)**, **interleukin-10**, interleukin-15, **interleukin-6**, ion channel, ions, irs1 gene\|irs1, jnk mitogen-activated protein kinases\|jun, kynurenine, lcn2 protein, human\|lcn2, lecithin, **leptin**, leptin\|lep, levodopa, linoleic acid, lysine, **magnesium***, mapt, mapt protein, human\|mapt, membrane transport proteins, mercury, metabotropic glutamate receptor 5, micrornas, minocycline, molecular chaperones, monoamine oxidase a, mthfr gene\|mthfr, muscarinic acetylcholine receptor, n-acetylaspartate, n-methyl-d-aspartate receptors, nefl gene\|nefl, **neopterin*,** nerve growth factors, nerve growth factors\|ngf, netrin-1, neuropeptide y\|npy, neuropeptides, neurosteroids, neurotransmitters, neurotrophic factor, neurotrophic tyrosine kinase receptor type 2\|ntrk2, nf-kappa b, nitric oxide, nitric oxide synthase type i, norepinephrine, opioid receptor, oralit, oxidoreductase, oxygen, **p-glycoprotein**, palmitic acid, peptidyl-dipeptidase a, per2 protein, mammalian, phenylpyruvate tautomerase, pink1, pituitary adenylate cyclase activating polypeptide, plasma proteins, plasminogen activator inhibitor 1, polyunsaturated fatty acids, promoter regions (genetics), protein kinases, psmb6\|yy1\|dll1, reactive oxygen species, receptor, receptor for advanced glycation endproducts, receptors, metabotropic glutamate, regulatory sequences, nucleic acid, reln gene\|reln, ros1, selenium, serine, serotonin, serotonin transporter, serum proteins, signaling molecule, slc30a3, slc6a4, ***sphingomyelins****, sri, streptozocin, sucla2 gene\|sucla2, sugars, tenascin, threonine, tie gene, timp1 protein, human\|timp1, tnf protein, human\|tnf, today, toll-like receptors, toxin, trace elements, transferrin receptor, transferrin\|tf, triglycerides, tumor necrosis factor-alpha\|tnf, uric acid, urine protein, **valine**, vascular factor, vegf protein, **human\|vegfa, vgf**, wnt3 gene\|wnt3, wwc1, zinc |

**Table S5.** Comparison of confounders - search results belonging to the drug or hormone semantic categories.

| | **Knowledge Graph** | **SemMedDB** |
|---|---|---|
| **Confounders** | anthocyanins, **anti-anxiety agents**, **antioxidants**, **antipsychotic agents**, **ascorbic acid**, butyrolactone, **caffeine**, ***ceramides****, ***cholesterol****, **copper**, diuretics, **docosahexaenoic acids**, **estradiol**, **estrogens**, **ethanol**, **folic acid**, **glucocorticoids**, **glucose**, | 2-cyclopentyl-5-(5-isoquinolylsulfonyl)-6-nitro-1h-benzo(d)imidazole, 3,4-methylenedioxyamphetamine, acetyl-l-carnitine, acetylcholinesterase inhibitors, acetylcysteine, active ingredients, adrenal cortex hormones, adrenergic beta-antagonists, air pollutants, alar, alba, alpha tocopherol, alpha-amino-3-hydroxy-5-methyl-4-isoxazolepropionic acid, alpha-amylase, alteplase, amides, amino acids, amisulpride, amsonic acid, analgesics, androgens, anesthetics, **anti-anxiety agents**, anti-inflammatory agents, anti-inflammatory agents, non-steroidal, anticholinergic agents, anticonvulsants, antidepressive agents, antidiabetics, antiepileptic agents, antihypertensive agents, **antioxidants**, **antipsychotic agents**, antithrombin iii, antiviral agents, arachidonic acid, arginine, aripiprazole, aromatase inhibitors, **ascorbic acid**, aspirin, astaxanthin, atorvastatin, atypical antipsychotic, basis, benzodiazepines, benzoic acid, beta-asarone, betaine, beverages, biogenic amines, biological response modifiers, brain-derived neurotrophic factor, human\|bdnf, butenolide, **caffeine**, calcifediol, calcium homeostasis, calcium ion, cannabidiol, cannabinoids, carbazole, catechin, celecoxib, chaihu-shugan-san, channel blockers, chelating agents, chinese herbs, chlorpyrifos, cholecalciferol, choline, cholinesterase inhibitors, chrysin, cilostazol, cisplatin, citalopram, citicoline, clozapine, coenzyme q10, coffee, compete, **copper**, corticosterone, corticotropin, corticotropin-releasing hormone, crocin, curcumin, cyclooxygenase 2 inhibitors, cyclooxygenase inhibitors, cysteinyl-leukotriene, dehydroepiandrosterone, dehydroepiandrosterone sulfate, deliver, dexamethasone, |

| | | |
|---|---|---|
| | glycocorticoids, **hydrocortisone**, hydroxide ion, **hydroxymethylglytaryl (CoA reductase inhibitors)**, ketones, **lipids**, ***low-density lipoproteins***, **magnesium***, **melatonin**, **metformin**, **neopterin***, **nicotine**, **polyphenols**, **polyunsaturated fatty acids**, **progesterone**, ramipril, **resveratrol**, ***saturated fat***, **scopolamine**, **selenium**, **simvastatin**, ***sphingomyelins****, ***steroids***, **testosterone**, ***tryglycerides***, **vitamin E** | diet, dihydromyricetin, dihydroxyacetone sulfate, dimebolin, dimethyl fumarate, **docosahexaenoic acids,** donepezil, dopamine, dopaminergic agents, drugs, investigational, drugs, non-prescription, duloxetine, duration, edaravone, eicosapentaenoic acid, ellagic acid, enoxaparin, epidermal growth factor\|egf, ergocalciferol, erythropoietin\|epo, escitalopram, **estradiol**, **estrogens**, etanercept, **ethanol**, experimental drug, fatty acids, omega-3, ferulic acid, fiber, fish oils, flavonoids, fluimucil\|xk\|nlrp1, fluoxetine, folate, **folic acid**, ***folic acid supplementation***, food, fruit, fty-720, galantamine, gamma hydroxybutyrate, general anesthetic drugs, geniposide, genistein, ginkgo biloba extract, ginsenoside rg1, glp-1 receptor agonist [epc], **glucocorticoids**, **glucose**, glutamine, gonadal steroid hormones, guanosine, hallucinogens, harmaline, harman, harmine, herbal drugs, histamine antagonists, honokiol, humanin, **hydrocortisone**, hydroxychloroquine, **hydroxymethylglutaryl-coa reductase inhibitors**, hypnotics, hypoglycemic agents, icariin, immunologic adjuvants, immunomodulators, inflammation mediators, infliximab, inositol, insulin, insulin\|ins, interferons, interleukin-1, beta, iron, isoflurane, kai-xin-san, ketoprofen, khat, kynurenic acid, lactate, lamotrigine, leptin, leptin\|lep, levetiracetam, levocarnitine, levothyroxine, linalool, **lipids**, lipopolysaccharides, liraglutide, lithium, lithium carbonate, lithium chloride, low-dose aspirin, magnolol, marihuana, medium chain triglycerides, mefloquine, melanin-concentrating hormone, **melatonin**, memantine, **metformin**, methamphetamine, methionine, methylate, methylphenidate, metric, mifepristone, modafinil, monoamine oxidase b inhibitor, monoamine oxidase inhibitors, n-methylaspartate, n,n-dimethylarginine, naproxen, naringenin, natalizumab, neuroprotective agents, new medications, nicergoline, **nicotine**, nicotinic agonists, nimodipine, nmda receptor antagonist, nonesterified fatty acids, nootropic agents, norepinephrine, nutraceuticals, nutrients, oils, volatile, omega-3 fatty acids, oralit, other medicated shampoos in atc, oxygen, paclitaxel, palmidrol, pesticides, phenylmethylpyrazolone, phosphatidylethanolamines, phosphodiesterase 5 inhibitor, phosphodiesterase inhibitors, phosphorus, pioglitazone, pk 11195, plant preparations, **polyphenols**, **polyunsaturated fatty acids**, pramipexol, pregnenolone, prescribed medications, **progesterone**, prolactin, propofol, psychostimulant, psychotropic drugs, pyruvates, quercetin, quetiapine, quetiapine fumarate, ramelteon, reboxetine, recombinant interferon-gamma, reproductive hormone, reserpine, **resveratrol**, reuptake inhibitors, riluzole, risperidone, rivastigmine, rosiglitazone, rosuvastatin, royal jelly, rutin, s-adenosylmethionine, salvianolic acid b, salvin, **scopolamine**, selective estrogen receptor modulators, selective norepinephrine re-uptake inhibitor, selective serotonin re-uptake inhibitor, selective serotonin reuptake inhibitors, selegiline, **selenium**, serine, serotonin, serotonin antagonists, serotonin uptake inhibitors, sertraline, silymarin, **simvastatin**, small molecule, sodium chloride, sodium valproate, **steroid hormone**, sulfur dioxide, synthetic drug, tadalafil, telmisartan, **testosterone**, tetrahydrocannabinol, thalidomide, theanine, therapeutic agent (substance), thiazolidinediones, thymoquinone, thyroid hormones, thyrotropin, thyroxine, timeline, tobacco, topiramate, trazodone, tretinoin, triiodothyronine, troxerutin, tryptophan, valproic acid, vegetables, verapamil, vilazodone, vitamin a, vitamin b 12, vitamin b complex, vitamin b12, vitamin d, vitamin e, vitamin supplementation, vitamins, zinc, zolpidem |

**Table S6.** Comparison with variables organized in rows by role (confounder, collider, or mediator) and columns organized by (knowledge graph (KG) and the two subsets of SemMedDB. The subsets (restricted to articles published in 2010 or after) include MachineReadingDB$_{\text{Scoped}}$ and SemMedDB$_{\text{Complete}}$. Concepts in bold appear in both the KG and SemMedDB$_{\text{Complete}}$. Underlined concepts appear in both SemMedDBs. To save space, we restrict our analysis of concepts retrieved to the semantic categories relating to findings and phenotypes. Strikethrough indicates that the variable appears in either the KG or SemMedDB$_{\text{Complete}}$ but not in the identical role(s).

| | **KG** | **machineReadingDB$_{\text{Scoped}}$** | **SemMedDB$_{\text{Complete}}$** |
|---|---|---|---|
| **Confounders** | amyloidosis, amyotrophic lateral sclerosis, atherosclerosis, atrial fibrillation, brain hemorrhage, cerebral amyloid angiopathy, **cerebral atrophy**, <u>**cerebrovascular accident**</u>, ***chronic infectious disease***, COPD, <u>**diabetes mellitus, diabetes mellitus (non-insulin -dependent)**</u>, encephalopathies, **hypercholesterolemia**, **hyperglycemia**, **hyperinsulinism**, **hypertensive disease**, **hypoglycemia**, **hypotension**, ***hypotension (orthostatic)***, ***increase in blood pressure***, infection recurrent, inflammatory response, **insulin resistance**, **ischemic stroke**, leuko-encephalopathy, ***low tension glaucoma***, **malnutrition**, **migraine disorders**, **myocardial infarction**, nerve degeneration, non-alcoholic fatty liver disease, **obesity**, **osteoporosis**, **overweight**, <u>***Parkinson's disorders***</u>, ***periodontal diseases***, rickets, **sleep apnea syndrome**, **sleep apnea (obstructive)**, unconscious state, **vitamin D deficiency** | **cerebrovascular accident**, deep brain stimulation, **diabetes mellitus, non-insulin-dependent***, diet (Mediterranean), hearing impairment, impaired cognition, metabolic syndrome, ***Parkinson's disease*** | agitation, amputation, ~~anemia~~, blind vision, body weight decreased, caloric restriction, cancer treatment, cardiovascular diseases, cardiovascular risk factor, **cerebral atrophy**, <u>**cerebrovascular accident**</u>, cerebrovascular disorders, ***chronic disease***, **chronic infectious disease**, chronic stress, convulsions, coronary arteriosclerosis, coronary artery bypass surgery, death (finding), debridement, <u>deep brain stimulation</u>, degenerative disorder, degenerative polyarthritis, depressed mood, depressive symptoms, <u>***diabetes***</u>, **diabetes mellitus,** <u>**diabetes mellitus (non-insulin- dependent)**</u>, <u>diet, mediterranean</u>, difficulty sleeping, disability, disrupted sleep, drowsiness, dyslipidemias, dyssomnias, endocrine system diseases, endothelial dysfunction, exercise, family history, feeling hopeless, fitness, folic acid deficiency, folic acid supplementation, frailty, functional disability, **glaucoma**, group exercise program, <u>hearing impairment</u>, heart diseases, heart failure, hyperactive behavior, **hypercholeresterolemia**, **hyperglycemia**, hypercysteinemia, **hyperinsulinism**, hyperlipidemia, hypersensitivity, **hypertensive disease**, hyperthyroidism, hypertrophy, **hypoglycemia**, hypotension, hypothyroidism, hysterectomy, illiteracy, <u>impaired cognition</u>, impulsivity, indifferent mood, infarction, lacunar, **insulin resistance**, intellectual functioning disability, **ischemic stroke**, kidney failure (chronic), lack of sensation, leisure activities, malaise, **malnutrition**, maternal history, memory training, memory skills training, <u>metabolic syndrome</u>, **migraine disorders**, moderate drinking, motor symptoms, multiple sclerosis, music therapy, **myocardial infarction**, narcotic use, need for isolation, neurodegenerative disorders, neurotic disorders, **obesity**, oestrogen therapy, **osteoporosis**, ***overweight***, oxidative stress, pain, <u>***Parkinson's disease***</u>, ***periodontitis***, personal satisfaction, phototherapy, physical activity/inactivity, physical therapy, planning suicide, pollution, porphyromonas gingivalis, pre-eclampsia, psoriasis, psychotherapy, regular exercise, reproductive history, rest, rheumatoid arthritis, sedentary lifestyle, seizures, sensory stimulation, **sleep apnea syndromes**, **sleep apnea (obstructive)**, sleep deprivation, sleep disorders, sleep disturbances, sleep disturbances, sleeplessness, smoking, spinal puncture, stress, stress disorders, post-traumatic, stress, acute, transcranial alternating/direct current stimulation, transplantation, underweight, vascular diseases, virus diseases, visual impairment, **vitamin D deficiency**, ***waist circumference***, widowhood, work stress, worried |
| **Colliders** | ~~anemia~~, apraxias, atherosclerosis, brain hemorrhage, cerebral amyloid angiopathy, **cerebrovascular accident**, congestive heart failure, deglutition disorders, **diabetes mellitus**, ***diabetes mellitus (non-insulin-dependent)***, encephalopathies, falls, fronto- temporal | NA | at risk for suicide, cardiovascular diseases, cerebral atrophy, **cerebrovascular accident**, **diabetes, diabetes mellitus**, epilepsy, feeling suicidal, ~~hypertensive disease~~, **ischemic stroke**, malignant neoplasms, **malnutrition**, ~~obesity~~, osteoporosis, ***Parkinson's disease***, **psychotic disorders**, rheumatoid arthritis, semantic impairment, sleep disturbances, suicide attempt |

| | | | |
|---|---|---|---|
| | dementia, immune response, ~~insulin resistance~~, **ischemic stroke**, **malnutrition**, neurofibrillary degeneration, **osteoporosis**, ***Parkinson disorders***, **pneumonia**, **psychotic disorders**, senile plaques, tauopathies | | |
| **Mediators** | amyotrophic lateral sclerosis, ~~anemia~~, atherosclerosis, brain hemorrhage, ***cerebral amyloid angiopathy***, **cerebrovascular accident**, ***congestive heart failure***, deglutition disorders, **diabetes mellitus**, ***diabetes mellitus (non-insulin-dependent)***, ~~insulin resistance~~, **ischemic stroke**, **malnutrition**, **osteoporosis**, ***Parkinson's disorders***, tauopathies | NA | blind vision, ***brain diseases***, cardiovascular diseases, cerebral atrophy, **cerebrovascular accident**, convulsions, ***diabetes***, **diabetes mellitus**, ***heart failure***, ~~hypercholesterolemia~~, ~~hypertensive disease~~, immune system diseases, **ischemic stroke**, **malnutrition**, metabolic syndrome, ~~migraine disorders~~, ~~obesity~~, **osteoporosis**, ***Parkinson's disease***, psoriasis, rheumatoid arthritis, semantic impairment, septicemia, sleep disturbances, stress disorders (post-traumatic), virus diseases, Wernicke encephalopathy |

**Table S7.** Comparison of search results belonging to the semantic categories of findings and phenotypes organized in rows by variable role, including role combinations ("Confounders only," "Colliders only," or "Mediators only") and columns organized by (knowledge graph (KG) and SemMedDB$_{\text{Complete}}$. We ignore MachineReadingDB$_{\text{Scoped}}$ for this analysis since it did not yield colliders or mediators. Concepts in bold appear in both the KG and SemMedDB$_{\text{Complete}}$. Closely related hypernymic/hyponymic concepts (near matched) are italicized in bold. Concepts are underlined if they appear in both SemMedDBs. A strikethrough (e.g., ~~anemia~~) indicates disagreement between knowledge bases.

| | **KG** | **SemMedDB$_{\text{Complete}}$** |
|---|---|---|
| **Confounders only** | amyloidosis, atrial fibrillation, ~~cerebral atrophy~~, **chronic infectious disease**, COPD, encephalopathies, ~~hypercholesterolemia~~, **hyperglycemia**, **hyperinsulinism**, ~~hypertensive disease~~, **hypoglycemia**, **hypotension**, ***hypotension (orthostatic)***, increase in blood pressure, ***infection recurrent***, inflammatory response, leukoencephalopathy, ***low tension glaucoma***, ~~migraine disorders~~, **myocardial infarction,** nerve degeneration, non-alcoholic fatty liver disease, ~~obesity~~, **overweight**, ***periodontal disease***, rickets, **sleep apnea syndrome**, **sleep apnea (obstructive)**, subarachnoid hemorrhage, unconscious state, **vitamin D deficiency** | agitation, amputation, ~~anemia~~, body weight decreased, caloric restriction, cancer treatment, cardiovascular risk factor, cerebrovascular disorders, ***chronic disease***, **chronic infectious disease**, chronic stress, coronary arteriosclerosis, coronary artery bypass surgery, death (finding), debridement, deep brain stimulation, degenerative disorder, degenerative polyarthritis, depressed mood, depressive symptoms, ~~diabetes mellitus, non-insulin-dependent, diet (mediterranean)~~, difficulty sleeping, disability, disrupted sleep, drowsiness, dyslipidemias, dyssomnias, endocrine system diseases, endothelial dysfunction, exercise, family history, feeling hopeless, fitness, folic acid deficiency, folic acid supplementation, frailty, functional disability, ***glaucoma***, group exercise program, hearing impairment, heart diseases, hyperactive behavior, **hyperglycemia**, hypercysteinemia, **hyperinsulinism**, hyperlipidemia, hypersensitivity, hyperthyroidism, hypertrophy, **hypoglycemia**, **hypotension**, hypothyroidism, hysterectomy, illiteracy, impaired cognition, impulsivity, indifferent mood, infarction, lacunar, ~~insulin resistance~~, intellectual functioning disability, kidney failure, chronic, lack of sensation, leisure activities, malaise, maternal history, memory training, memory skills training, moderate drinking, motor symptoms, multiple sclerosis, music therapy, **myocardial infarction**, narcotic use, need for isolation, neurodegenerative disorders, neurotic disorders, oestrogen therapy, **overweight,** oxidative stress, pain, ***periodontitis***, personal satisfaction, phototherapy, physical activity/inactivity, physical therapy, planning suicide, pollution, porphyromonas gingivalis, pre-eclampsia, psychotherapy, regular exercise, reproductive history, rest, sedentary lifestyle, seizures, sensory stimulation, **sleep apnea syndromes**, **sleep apnea, obstructive**, sleep deprivation, sleep disorders, sleep disturbances, sleeplessness, smoking, spinal puncture, stress, stress disorders, transcranial alternating/direct current stimulation, transplantation, underweight, vascular diseases, visual impairment, **vitamin D deficiency**, ***waist circumference***, widowhood, work stress, worried |
| **Colliders only** | apraxias, encephalitis, falls, frontotemporal dementia, immune response, neurofibrillary degeneration, pneumonia, **psychotic disorders**, senile plaques | at risk for suicide, epilepsy, feeling suicidal, malignant neoplasms, **psychotic disorders**, suicide attempt |
| **Mediators only** | NA | immune system diseases, septicemia, Wernicke encephalopathy |
| **Confounder/ mediators** | amyotrophic lateral sclerosis | blind vision, convulsions, ~~heart failure~~, ~~hypercholesterolemia~~, metabolic syndrome, ~~migraine disorders~~, psoriasis, stress disorders (post-traumatic), virus diseases |
| **Collider/ confounders** | encephalopathies | NA |
| **Collider/ mediators** | ~~anemia~~, ~~congestive heart failure~~, deglutition disorders, tauopathies | brain diseases, semantic impairment |

| | | |
|---|---|---|
| **Collider/ confounder/ mediators** | amyloid angiopathy, atherosclerosis, brain hemorrhage, **cerebral cerebrovascular accident**, **diabetes mellitus,** ***diabetes mellitus (non-insulin -dependent)***, ~~insulin resistance~~, **ischemic stroke**, **malnutrition**, **osteoporosis**, ***Parkinson disorders****, stroke* | cardiovascular diseases, ~~cerebral atrophy~~, **cerebrovascular accident**, ***diabetes***, **diabetes mellitus**, ~~hypertensive disease~~, **ischemic stroke**, **malnutrition**, ~~obesity~~, **osteoporosis**, ***Parkinson's disease***, rheumatoid arthritis, sleep disturbances |

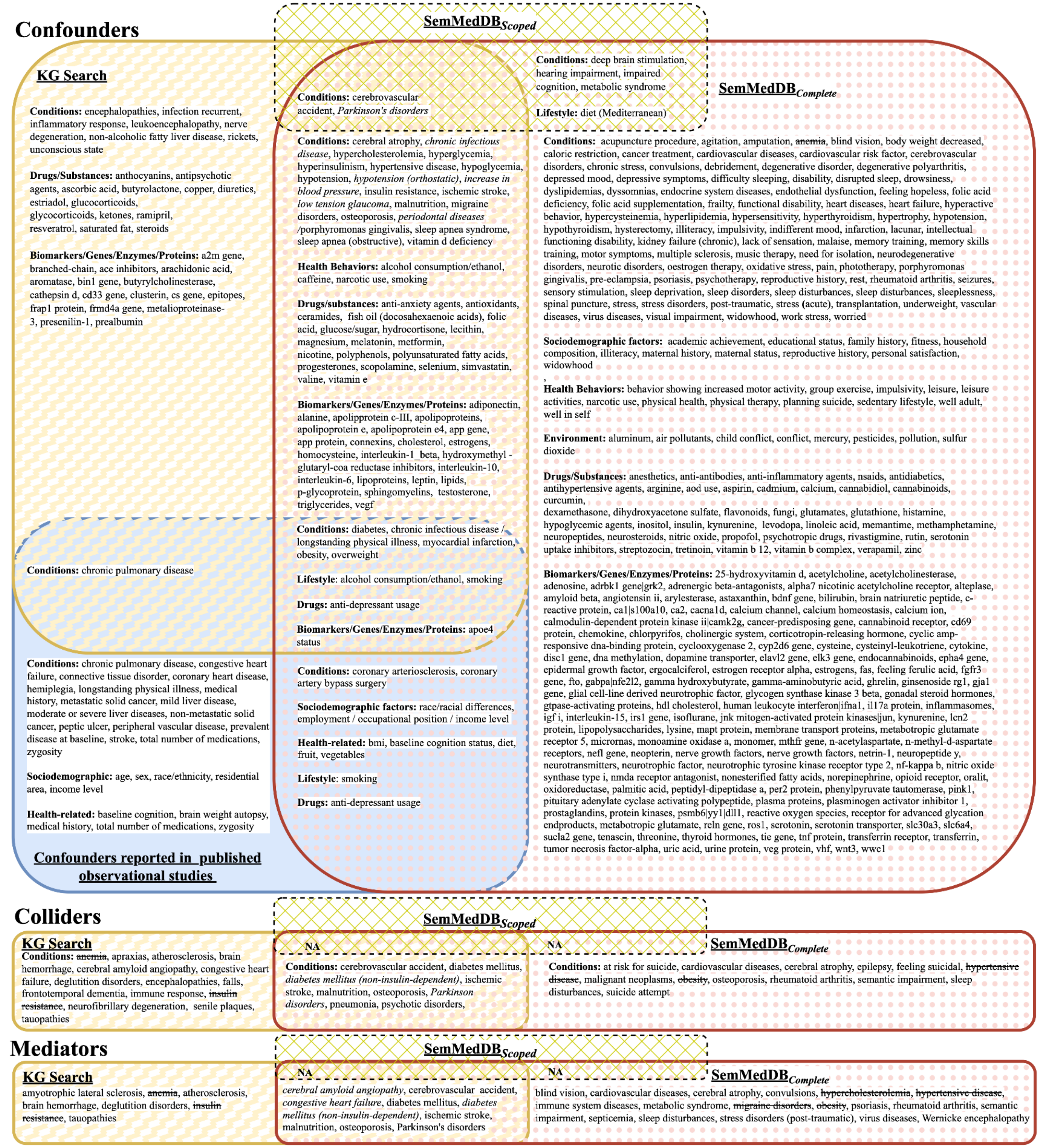


**Figure S1.** This figure shows the complete variables/concepts for simple (not combined) causal roles from KG search in yellow, $\text{SemMedDB}_{\text{Scoped}}$ in green diagonal and $\text{SemMedDB}_{\text{Complete}}$ in rose polka dot, and the confounders from the pilot-metareview in solid blue as a Venn diagram.

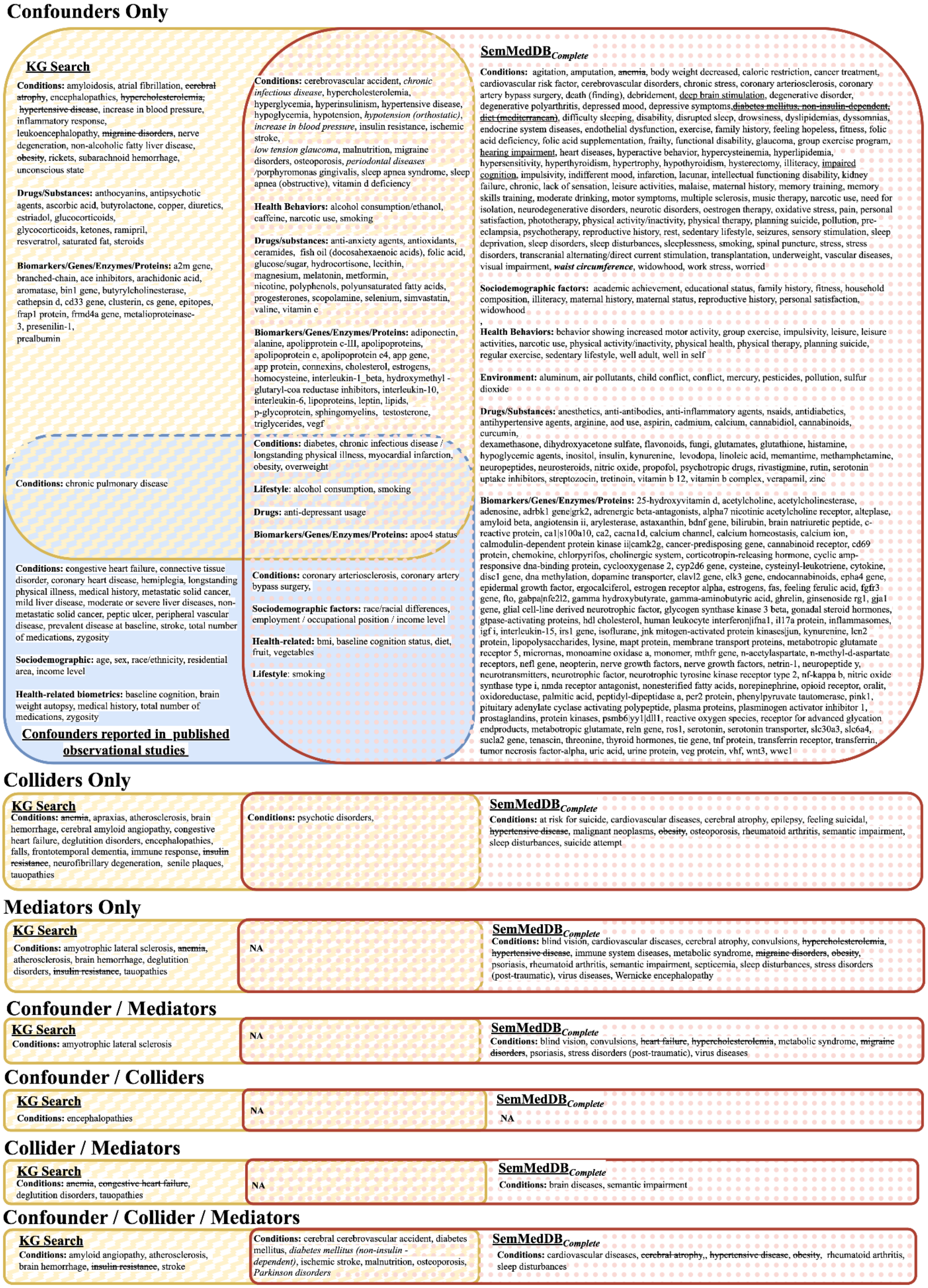


**Figure S2.** This figure shows the complete variables/concepts for combined causal roles from KG search in yellow, $\text{SemMedDB}_{\text{Scoped}}$ in green diagonal and $\text{SemMedDB}_{\text{Complete}}$ in rose polka dot, and the confounders from the pilot-metareview in solid blue as a Venn diagram.
