## Supplementary material for "Causal feature selection using a knowledge graph combining structured knowledge from the biomedical literature and ontologies: a use case studying depression as a risk factor for Alzheimer’s disease": File VII. Appendix_C.docx. This file contains documentation for access the Zenodo archive containing Jupyter notebooks for knowledge hygiene operation

### Supplementary Material Appendix C.

**File VII.** ***Appendix_C.docx***.

**Documentation for key contents of the project code archive on Zenodo** **1**
Key files and their locations in the Zenodo archive. 1
1. Accessing the Zenodo archive 1
2. Data pre-processing: UMLS-to-OBO mappings 1
3. Data pre-processing: Jupyter notebook with code for mappings and logical closure 1
4. KG search: SPARQL queries 2
a. Confounder SPARQL query 2
b. Collider SPARQL query 2
c. Mediator SPARQL query 2
5. Comparing KG search with SemMedDB: SemMedDB queries 3

#### Documentation for key contents of the project code archive on Zenodo

##### Key files and their locations in the Zenodo archive.

###### 1. Accessing the Zenodo archive

Steps to accessing the Zenodo archive

1. Visit the Zenodo archive: https://doi.org/10.5281/zenodo.6785307
2. Download the zip and unzip file: **`alz_lbgcm-master.zip`**
3. Follow the instructions below to access particular research data components.

###### 2. Data pre-processing: UMLS-to-OBO mappings

This section is pertinent to sections 2.3 and 2.4 of the paper. The research data produced for the terminology mapping and logical closure components of the workflow are available in the following folder:

```
alz_lbgcm/clipspy-KG/
```

UMLS to OBO ontology mappings are available here:

```
alz_lbgcm/clipspy-KG/cui_to_ontology_maps
```

The code for executing the mappings is in the Jupyter notebook in #3.

#### 3. Data pre-processing: Jupyter notebook with code for mappings and logical closure

The research data produced for the terminology mapping and logical closure components of the workflow are available in the following folder:

`alz_lbgcm/clipspy-KG/`

The OBO to UMLS and logical closure Jupyter notebook is available here:

`alz_lbgcm/clipspy-KG/ad-kg-v1.ipynb`

Inside that folder, there are other files, some of which are required to run the Jupyter notebook. In order to execute these functions, you will need to install CLIPS, available at this link: https://clipsrules.net/.

#### 4. KG search: SPARQL queries

##### a. Confounder SPARQL query

The SPARQL query for confounders is available in the folder:

`alz_lbgcm/combinedGraphAnalysis/adConfounderSearch`

Inside this folder, the SPARQL query for confounders is here:

`depression-AD-confounder-search.sparql`

The raw output from running the confounder SPARQL query on the combined KG is here:

`MR-and-lit-review-DEPRESSION-confounder-set-ALL.txt`

##### b. Collider SPARQL query

The SPARQL query for colliders is available in the folder:

`alz_lbgcm/combinedGraphAnalysis/adColliderSearch`

Inside this folder, the SPARQL query for colliders is here:

`depression-AD-collider-search.sparql`

The raw output from running the collider SPARQL query on the combined KG is here:

`MR-and-lit-review-DEPRESSION-collider-set-ALL.txt`

##### c. Mediator SPARQL query

The SPARQL query for mediators is available in the folder:

**`alz_lbgcm/combinedGraphAnalysis/adMediatorSearch`**

Inside this folder, the SPARQL query for mediators is here:

**`depression-AD-mediator-search.sparql`**

The raw output from running the mediator SPARQL query on the combined KG is here:

**`MR-and-lit-review-DEPRESSION-mediator-set-ALL.txt`**

#### 5. Comparing KG search with SemMedDB: SemMedDB queries

The SemMedDB code is in R and SQL and queries a version of SemMedDB converted to PostgreSQL relational database system.

1. Searching SemMedDBScoped: https://zenodo.org/record/6785307/files/getKnowledgeD2AD_scopedSemRep.R?download=1
2. Searching SemMedDBComplete: https://zenodo.org/record/6785307/files/getKnowledge_depression2AD.R?download=1
